# Structural similarities reveal an expansive conotoxin family with a two-finger toxin fold

**DOI:** 10.1101/2025.07.03.662903

**Authors:** Muhammad Saad Khilji, Celeste M. Hackney, Thomas L. Koch, Arik J. Hone, Aymeric Rogalski, Maren Watkins, Jortan Tun, J. Michael McIntosh, Baldomero Olivera, Helena Safavi-Hemami, Kaare Teilum, Lars Ellgaard

## Abstract

Venomous animals have evolved a diverse repertoire of toxins with considerable pharmaceutical potential. The rapid evolution of peptide toxins, such as the conotoxins produced by venomous marine cone snails, often complicates efforts to infer their evolutionary relationships based solely on sequence information. Structural bioinformatics, however, can provide robust support. Here, we first solve the NMR structure of a macro-conotoxin from the MLSML superfamily, Tx33.1, which is composed of 124 residues, including 12 cysteines. We then apply deep learning-based methods for structure prediction and comparison to identify structural similarities between this toxin and five additional, previously uncharacterized conotoxin superfamilies. Although only three of these superfamilies exhibit sequence homology, a combined approach incorporating structure prediction, structure comparison, and gene structure analysis supports the conclusion that all six superfamilies share a common evolutionary past. The Tx33.1 NMR structure displays similarity to the first two domains of Argos, a secretory protein from *Drosophila melanogaster* that comprises three domains, each harboring two short β-stranded loops (“fingers”). Consequently, we propose the name “two-finger toxin (2FTX)” fold for this type of domain. Finally, using structure similarity searches, we identify a wide range of 2FTX proteins in protostomes, including non-venom-derived, secretory cone snail proteins. This study demonstrates how structural bioinformatics can be employed to uncover evolutionary relationships among rapidly evolving genes. It simultaneously identifies a large, previously unrecognized group of protostome 2FTX proteins, many of which exhibit close structural similarity to Argos and may perform a similar function in regulating EGFR signaling.

## Introduction

Animal venoms are rich in bioactive molecules that are used to kill prey and deter predators. Peptide toxins specifically target molecules such as ion channels, G-protein-coupled receptors, and transporters in their recipients. Due to high target specificity and potency, peptide toxins are valuable compounds that can be explored for use as research tools and potential drug leads. Currently, 11 venom-derived peptides originating from cone snails, snakes, leeches, and lizards have been approved for clinical applications (Bordon, et al. 2020).

Often, toxin evolution involves duplication of a gene encoding a protein with an endophysiological function followed by the “recruitment” of the duplicated copy into venom in a process accompanied by neo- and/or sub-functionalization (Jackson and Koludarov 2020). Small peptide toxins generally lack a classical hydrophobic core – instead, most are structurally stabilized by disulfide bonds involving conserved cysteine residues. This characteristic feature provides a stable scaffold that can evolve novel functional properties through variation on inter-cysteine positions, which correspond to surface-exposed residues often exhibiting high evolutionary rates driven by positive selection (Woodward, et al. 1990).

Given the often high mutational rates of peptide toxins combined with their small size, inferring evolutionary histories based solely on sequences can be challenging. Because three-dimensional structure is more conserved through evolution than sequence (Illergard, et al. 2009), structural analysis can aid in establishing distant evolutionary relationships (Undheim, et al. 2016), e.g., between toxins and their ancestral endophysiological counterparts or among evolutionarily related toxins with highly diverged sequences. For instance, structural homology of the spider and centipede toxins Ta1a and Ssm6a revealed the convergent recruitment of these peptides from endophysiological peptides of the crustacean hyperglycemic hormone (CHH) family into venom (Undheim, et al. 2015). Conversely, the AlphaFold (AF)-predicted structure of a newly identified cone snail toxin was used in a structure similarity search to identify the first endogenous molluscan peptide of the CHH family (Koch, et al. 2023). In another example, we recently established an evolutionary relationship between the con-ikot-ikot and Mu8.1 conotoxins that have highly divergent mature sequences but similar three-dimensional and gene structures (Hackney, et al. 2023).

Conotoxins represent a group of particularly fast-evolving gene products (Duda and Palumbi 1999). Each of the approximately 1000 extant species of cone snails expresses hundreds of mainly disulfide-rich toxins. Conotoxins are produced as precursor sequences encoding a signal sequence for targeting to the endoplasmic reticulum (ER), often followed by a propeptide that is proteolytically removed upon exit from the ER, and finally, the mature sequence. Although larger toxins are also known, most characterized conotoxins are short (approx. 15–45 residues for the mature peptide region). Conventionally, conotoxins are grouped into superfamilies based on similarity in their signal sequence regions. To date, approximately 70 gene superfamilies of conotoxins have been described (Robinson, et al. 2014; Safavi-Hemami, et al. 2018; Koch, et al. 2024).

Conotoxins can be classified into structurally diverse classes that typically share specific biological activities towards target proteins, although exceptions exist. Known structures include, e.g., the inhibitor cystine knot (ICK) (Undheim, et al. 2016), Kunitz (Mourao and Schwartz 2013), insulin (Laugesen, et al. 2022), saposin-like (Hackney, et al. 2023), and mini-granulin folds (Nielsen, et al. 2019; Raffaelli, et al. 2024). Structural motifs such as the ICK fold, conserved cysteine-rich domain, and Kunitz-type folds are found across a wide range of venomous animals. By contrast, despite their prevalence in caenophidian snakes (Koludarov, et al. 2023), the three-finger toxins (3FTXs) have apparently not been found in other venomous animals. 3FTXs belong to the Ly6/uPAR superfamily of proteins that share the same overall fold consisting of a disulfide-bonded globular core from which three β-stranded loops (or “fingers”) protrude (Kessler, et al. 2017; Leth, et al. 2019; Koludarov, et al. 2023).

In recent years, technological advances in the fields of transcriptomics and proteomics have greatly expanded the number of available conotoxin sequences (Terrat, et al. 2012; Safavi-Hemami, et al. 2014; Peng, et al. 2016; Degueldre, et al. 2017; Gao, et al. 2018; von Reumont, et al. 2022; Fedosov, et al. 2023). It has thus become apparent that several previously uncharacterized toxin superfamilies encode proteins exceeding 50 amino acid residues (Koch, et al. 2024). These “macro-conotoxins” have been challenging to produce and characterize due to their complicated structures, often comprising more than three disulfide bonds. However, the development of various systems for recombinant expression of disulfide-rich proteins (Klint, et al. 2013; Nielsen, et al. 2019; Rivera-de-Torre, et al. 2021; Hackney, et al. 2023), along with advances in chemical synthesis and protein ligation methods (recently reviewed in (Ho, et al. 2025)), has made these proteins amenable to biochemical and pharmacological characterization. Since macro-conotoxins are less well-characterized than short conotoxins, their investigation is bound to provide new insights into conotoxin structure, function, and evolution.

Here, we use structure- and sequence-based analyses of six previously uncharacterized macro-conotoxin superfamilies to uncover their evolutionary relationships. Using sequences and structures from these superfamilies as queries, we extend our analysis beyond cone snails to identify a large group of protostome proteins that adopt a fold similar to Argos, a secretory protein from *Drosophila melanogaster* that binds the epidermal growth factor (EGF) ligand called Spitz and harbors three two-finger toxin (2FTX) domains. Our findings demonstrate that combining structure determination and prediction with bioinformatics analyses provides a powerful approach for uncovering evolutionary connections between seemingly unrelated sequences. Moreover, this work identifies a previously unrecognized and extensive group of protostome 2FTX proteins of unknown functions.

## Results

### Recombinant production of a newly identified macro-conotoxin, Tx33.1, in the fully oxidized form

To identify novel classes of macro-conotoxins, we analyzed the transcriptome of the venom gland of the mollusk-hunter *Conus textile,* a species with well-characterized short conotoxins (Ueberheide, et al. 2009). This led to the identification of an abundant transcript (transcript per million (TPM) value: 2,188) encoding a peptide with an N-terminal signal sequence and a predicted mature toxin of 124 residues (Fig. 1A and Fig. S1A). The precursor sequence of this toxin lacks a pro-peptide region. This toxin was named Tx33.1 (Robinson, et al. 2017), where “Tx” denotes the two-letter species abbreviation for *C. textile*, “33” indicates the cysteine framework, and “1” signifies that it is the first toxin described from this gene family. To identify orthologs of Tx33.1 in other cone snail species we employed similarity searches against assembled cone snail venom gland transcriptomes, the NCBI non-redundant protein database, and NCBI transcriptome shotgun assembly cone snail databases. Using this approach, we identified 63 sequences (of which 50 were full-length; see Supporting File 1) from 25 species representing snail-, worm- and fish-hunting *Conus* species. The signal sequences contain a “MLSMLAWTLMTAMVVMNA” consensus motif (Fig. S1A). Based on the first five residues of the signal sequence, this superfamily of conotoxins is designated as the MLSML superfamily (Koch, et al. 2024). All identified sequences contain 12 conserved cysteine residues in the mature toxin region (Fig. 1A and Fig. S1A).

**Figure 1.**
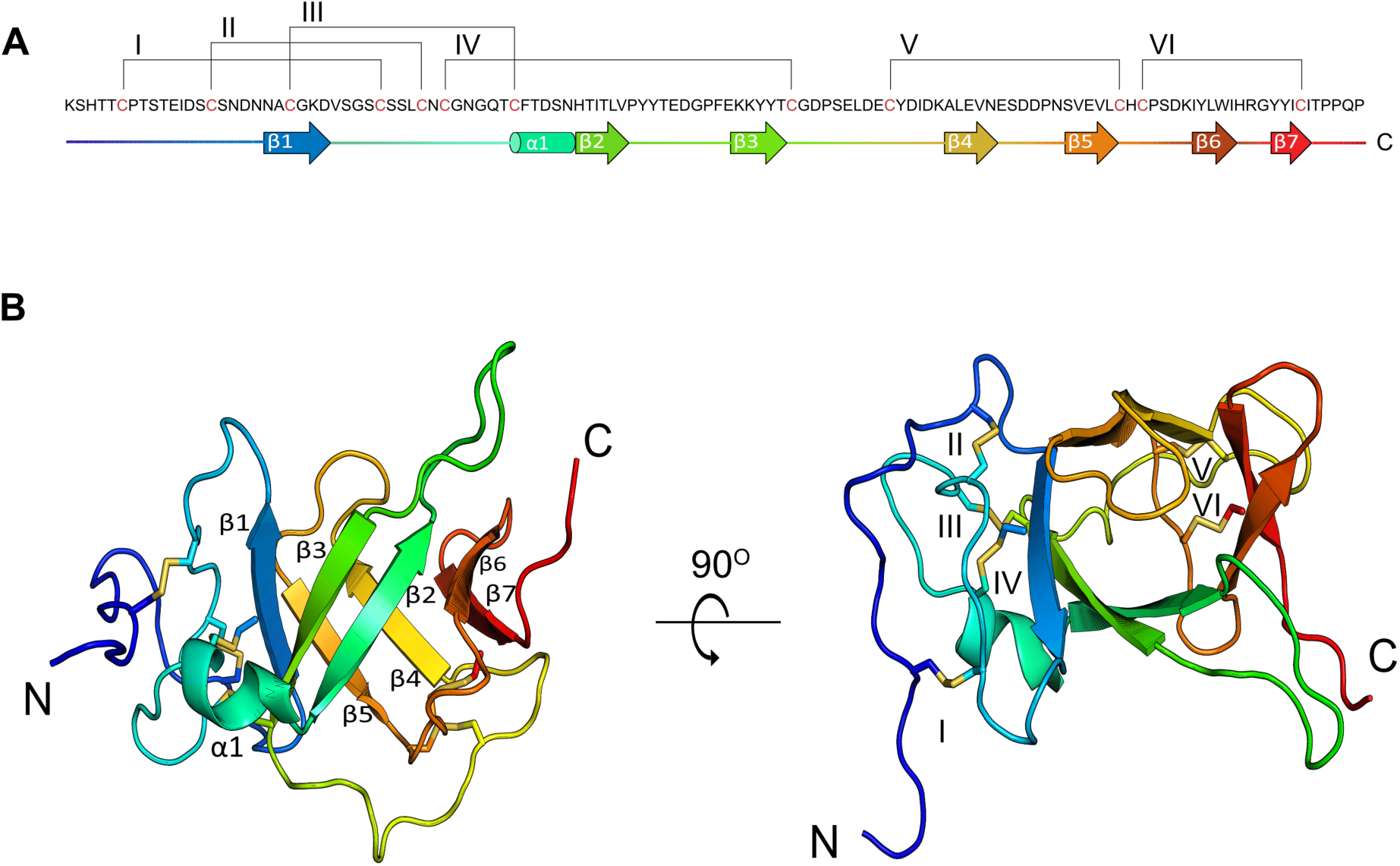
NMR structure of Tx33.1. **(A)** Sequence of mature Tx33.1 with graphic representation of regular secondary structure elements numbered by Arabic numerals and represented as arrows (β-strands) and cylinders (α-helix). Disulfide bonds are represented by brackets and numbered by Roman numerals. Cysteine residues are colored red. **(B)** Cartoon representation of the Tx33.1 NMR structure containing seven β-strands (β1-β7) and one α-helix (α1) shown in two different orientations. The disulfide bonds are shown as yellow stick models and labelled with Roman numerals as in Panel A.

To further investigate this toxin, Tx33.1 was recombinantly expressed in BL21(DE3) *E. coli* cells using the csCyDisCo system (Nielsen, et al. 2019; Bertelsen, et al. 2021), a derivative of the original CyDisCo system (Gaciarz, et al. 2017). Briefly, this systems allows for production of correctly folded, fully oxidized disulfide-rich proteins in the bacterial cytosol by co-expression of three foldases – the Erv1p oxidase, as well as human protein disulfide isomerase (hPDI) and a conotoxin-specific PDI (csPDI) found to be highly expressed in the venom gland of many cone snail species (Safavi-Hemami, et al. 2016). This platform has previously been employed to produce other conotoxins in their correctly folded forms (Nielsen, et al. 2019; Hackney, et al. 2023; Müller, et al. 2023). Tx33.1 was expressed as a C-terminal fusion to an engineered variant of ubiquitin (Ub) containing 10 consecutive histidines (Ub-His_10_) with a cleavage site for Tobacco Etch Virus (TEV) protease, allowing for the release of the mature toxin (Rogov, et al. 2012).

SDS-PAGE analysis showed that co-expression of Ub-His_10_-Tx33.1 with Erv1p, hPDI, and csPDI significantly increased the yield of soluble protein, with the large majority of Ub-His_10_-Tx33.1 appearing in the soluble fraction (Fig. S2A). Subsequent purification steps, involving immobilized metal affinity chromatography (IMAC), TEV protease cleavage, reverse IMAC, and anion exchange chromatography, yielded highly pure protein (>95% as judged by SDS-PAGE analysis; Fig. S2B), with a final yield of approximately 8 mg/L of culture.

Containing 12 cysteine residues, Tx33.1 is predicted to form six disulfide bonds. Full oxidation was confirmed using Q-TOF mass spectrometry, which measured a monoisotopic molecular mass of 13,703.80 Da, closely matching the theoretical mass of 13,703.95 Da for the fully oxidized protein (Fig. S3A). Additional evidence that Tx33.1 adopts a folded conformation was provided by a well-dispersed [^1^H–^15^N]-heteronuclear single quantum coherence spectrum (Fig. S3B).

### Tx33.1 displays a previously unknown toxin fold

We next determined the three-dimensional structure of Tx33.1 in solution using NMR spectroscopy. In total, 83% of all chemical shifts were assigned, including 88% of backbone ^1^H, ^13^C, and ^15^N resonances, and 80% of sidechain resonances. A total of 1,320 short-range distance restraints derived from NOESY spectra, together with 170 dihedral angle restraints calculated from chemical shifts, were used in structure calculations (Table 1).

**Table 1.**
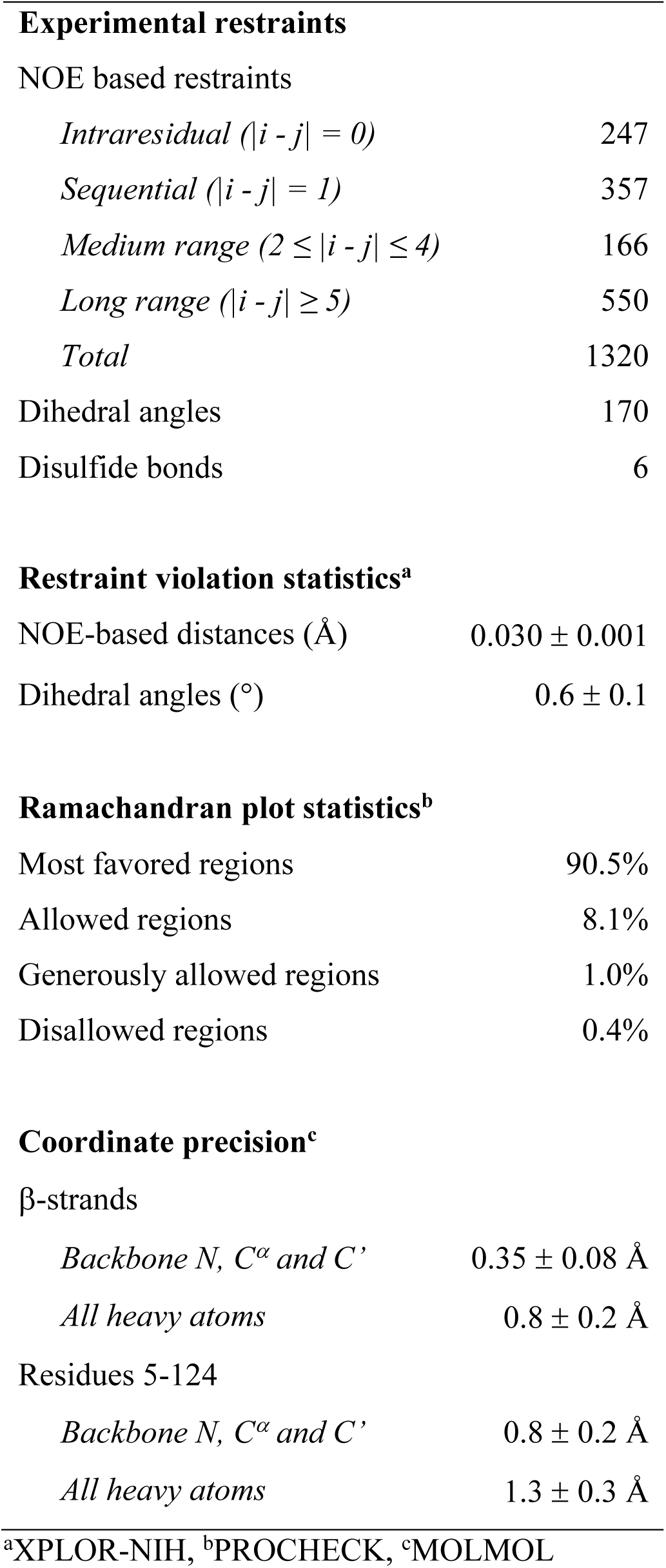
Statistics for the ensemble of 20 conformers representing Tx33.1 (PDB: 9FRQ)

The Tx33.1 structure is well defined from residue 5 to residue 124, with a pairwise root-mean-square deviation (RMSD) of 0.8 ± 0.2 Å for the 20 models representing the structure (Fig. S4). The core of the Tx33.1 structure constitutes a seven-stranded β-barrel (Fig. 1B), with a pairwise RMSD of 0.35 Å. The residues prior to and immediately following β1 constitute a loop region held together by the first four disulfide bonds (Cys6–Cys30, Cys15–Cys34, Cys22– Cys42 and Cys36–Cys69; Fig. 1). A small alpha helix (α1) comprising six residues is present between β1 and β2. The last two disulfide bonds (Cys78–Cys100 and Cys102–Cys118; Fig. 1) connect loop regions in the C-terminal region of the protein. A notably long, yet well-defined loop of 10 residues is present between β2 and β3. The loop between β4 and β5 comprises six residues and constitutes a lid-like structure placed at one end of the β-barrel. Three residues, Gly23, His48, and Thr57, are strictly conserved within the MLSML family, suggesting they may play a functional role.

In summary, the NMR structure of Tx33.1 revealed a fold rich in β-strand structure that has not previously been observed in known venom proteins.

### Bioinformatics uncovers six structurally and evolutionarily related macro-conotoxin superfamilies

Using the sequence and structure of Tx33.1, we next investigated the potential presence of related families of macro-conotoxins. We used structural homology searching against predicted structures of a nonredundant set of 3,412 conotoxin precursors extracted from venom gland RNAseq data from 42 cone snail species (Koch, et al. 2024). Using this approach, we identified five additional uncharacterized conotoxin families – the MMLFM, “Unknown”, MARFL, unk2 and E superfamilies – with a near-identical cysteine framework (Fig. 2A) that exhibit structural similarity to Tx33.1 (Fig. 2B). Structure-based phylogenetic analysis supported this notion, demonstrating high similarity among the families (Fig. 2C). However, on a sequence-level, only the MMLFM and Unknown superfamilies show similarity to the MLSML conotoxins, as demonstrated using sequence-based CLANS (CLuster ANalysis of Sequences) clustering (Frickey and Lupas 2004) (Fig. 2D) – a method for visualizing protein families based on all-against-all pairwise protein sequence similarities – and maximum likelihood tree reconstruction (Fig. 2E). Overall, the phylogenetic reconstruction based on the structural predictions (Fig. 2C) revealed a much stronger association among the six conotoxin superfamilies compared to sequence-based analyses (Fig. 2E). While the superfamilies largely formed monophyletic groups in the structural analysis (Fig. 2E), the short branch lengths of the deep branches in the tree shown in Fig. 2C suggest a common ancestor for all six superfamilies. Multiple sequence alignments of the proteins from the MLSML, MMLFM, Unknown, MARFL, unk2, and E superfamilies are provided in Fig. S1. All protein sequences used for the analysis in Fig. 2 are included in Supporting File 2.

**Figure 2.**
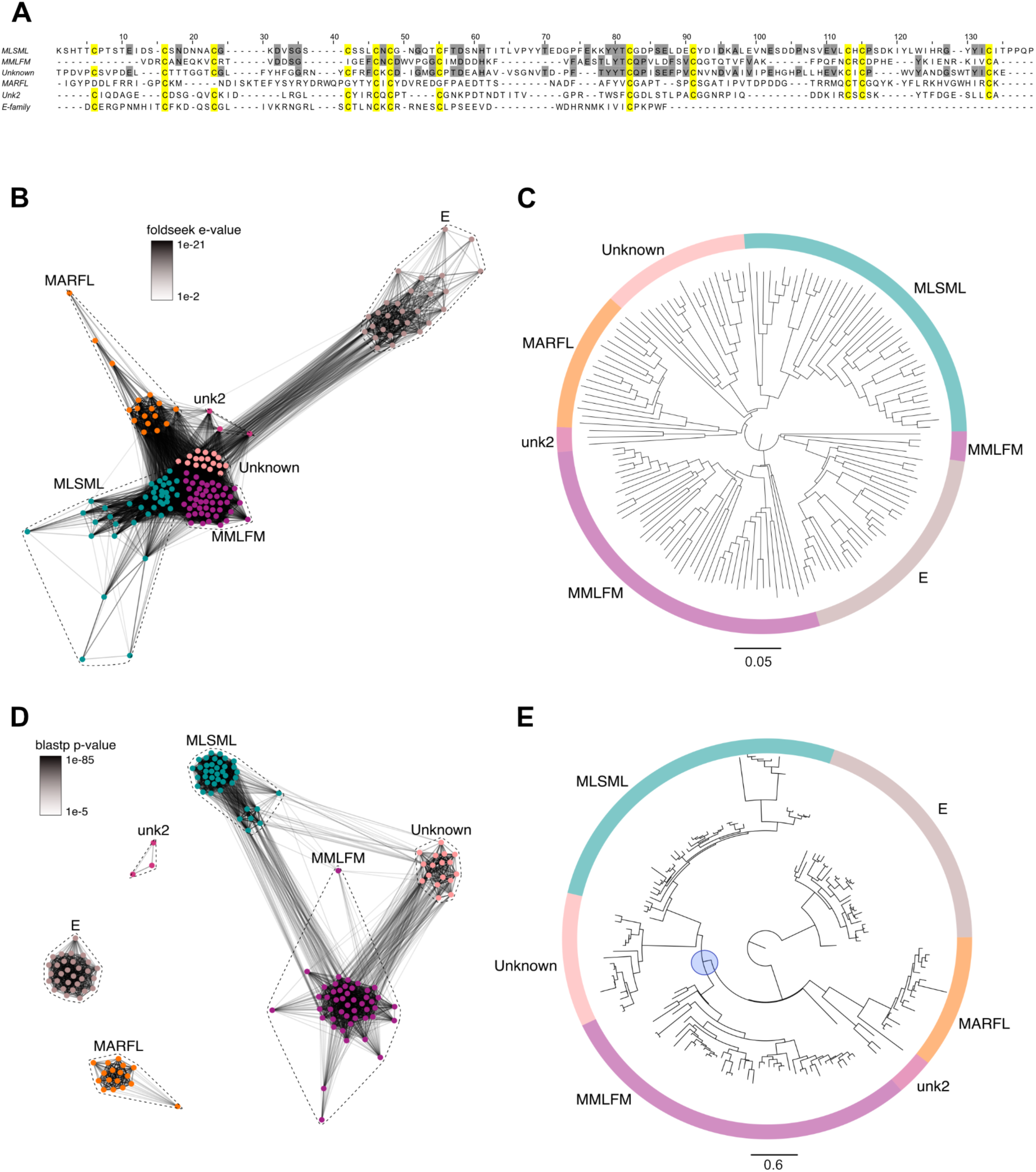
Sequence and structural similarity among six conotoxin superfamilies. **(A)** Multiple sequence alignment between proteins belonging to the MLSML (Tx33.1 from *Conus textile*), MMLFM (from *Conus coronatus*), Unknown (from *Conus raulsilvai*), MARFL (from *Conus ventricosus*), unk2 (from *Conus rattus*), and E-superfamilies (from *Conus gloriamaris*). Cysteine residues are highlighted in yellow. Additional identical residues in the top three sequences are highlighted in gray. **(B)** Foldseek structural similarity CLANS cluster of conotoxins from the MLSML, MMLFM, Unknown, MARFL, unk2, and E-superfamilies. The structures were predicted for each of the toxin precursor sequences using AlphaFold 2, and structural similarities were calculated using Foldseek easy-cluster. The e-values were extracted and used to attract similar structures. **(C)** Foldtree reconstruction of conotoxin phylogeny performed with the predicted AlphaFold 2 structures used as input in Panel B. **(D)** BLOSUM62 cluster map of conotoxins from the MLSML, MMLFM, Unknown, MARFL, unk2, and E-superfamilies. The nodes represent individual precursor sequences, edges correspond to blastp p-values above 1e-5. Nodes are color coded according to conotoxin superfamily as indicated. **(E)** Maximum-likelihood gene tree of the same sequences used as input for Panel D. The tree was rooted with the E-superfamily, which had the longest branch. The blue circle marks the node indicating that the MLSML, Unknown and MMLFM superfamilies form a monophyletic clade. All sequences used for the analyses in this figure are included in Supporting File 2.

To further investigate whether the six macro-conotoxin superfamilies share characteristic features with classical conotoxins, we examined their expression levels, signatures of positive selection, and their degree of sequence variation. Analysis of the sequences of the six identified superfamilies revealed that in addition to high transcriptional levels (Fig. S5 and Supporting File 3), five of the six superfamilies show signs of positive selection (due to the limited sample size, the unk2 superfamily was not included in this analysis). Estimated pairwise dN/dS ratios showed mean dN/dS values of 6.91, 1.93, 2.48, 2.40, and 3.31 across the MLSML, MMLFM, Unknown, MARFL, and E-superfamilies, respectively. Site-models of selection (comparing M7 and M8 models in the Phylogenetic Analysis by Maximum Likelihood (PAML) package (Yang 1997)) further supported the presence of positive selection at multiple sites throughout the sequences of these toxins (72 of 179 codons in MLSML, 66 of 165 codons in MMLFM, 81 of 153 codons in Unknown, 44 of 162 codons in MARFL, and 43 of 96 codons in E). Plotting the sequence conservation along the precursors further revealed that several inter-cysteine regions exhibit a high degree of variation (Fig. S6). These sequence features, typical of conotoxin-encoding genes, support the role of these proteins as functional venom components. In addition to the structural similarity, gene structure analysis using the published genomes of *Conus ventricosus, Conus betulinus*, and *Conus canariensis*, demonstrated that five of the six superfamilies (MLSML, MMLFM, Unknown, MARFL, and unk2) share similar gene structures, each comprising four precursor-encoding exons with conserved intron phases (0, 2, and 0, respectively, with a single exception) (Fig. S7). E-superfamily genes deviate slightly from this structure and are encoded on three exons that share intron phases (0 and 2) with the first two introns of the other superfamilies (Fig. S7). The positions of the conserved cysteines involved in disulfide-bond formation are well preserved across the exons in all superfamilies, further supporting a common evolutionary origin (Fig. S7).

Collectively, these findings reveal that the six conotoxin superfamilies share a common evolutionary origin and possess similar three-dimensional structures, despite strong divergence at non-cysteine positions in three of the superfamilies.

### Comparative structural analysis of Tx33.1 and AF-predicted conotoxin structures

The results in Fig. 2B demonstrate structural similarity among proteins from the six superfamilies at an overall level based on their AF-predicted structures. Consequently, we conducted a more detailed comparison of the NMR structure of Tx33.1 with the predicted structures of the other five superfamilies.

First, we note that the NMR and AF-predicted structures of Tx33.1 are largely identical with an RMSD of 1.5 Å (Fig. 3A), demonstrating AlphaFold’s accuracy in predicting this fold. The main differences are observed in the N-terminal region, where the predicted structure includes an additional β-strand. Moreover, β1 in the NMR structure exhibits a slightly different orientation compared to the corresponding strand in the predicted structure. These subtle differences lead to the formation of the β-barrel in the NMR structure, whereas the predicted structure rather displays two domains, each containing two short β-hairpins. This same overall fold is also observed in the predicted structures of proteins from the MMLFM (Fig. 3B), Unknown, MARFL, and unk2 superfamilies (Fig. S8), although the number of predicted β-strands varies slightly. In the E-superfamily, the mature sequences consist of approximately 65 residues that align with the N-terminal half of the proteins from the five other conotoxin superfamilies (Fig. 2A and Fig. S2E). Accordingly, the AF-predicted structures of E-superfamily proteins are similar to the structure of the N-terminal half of Tx33.1 (Fig. S8D).

**Figure 3.**
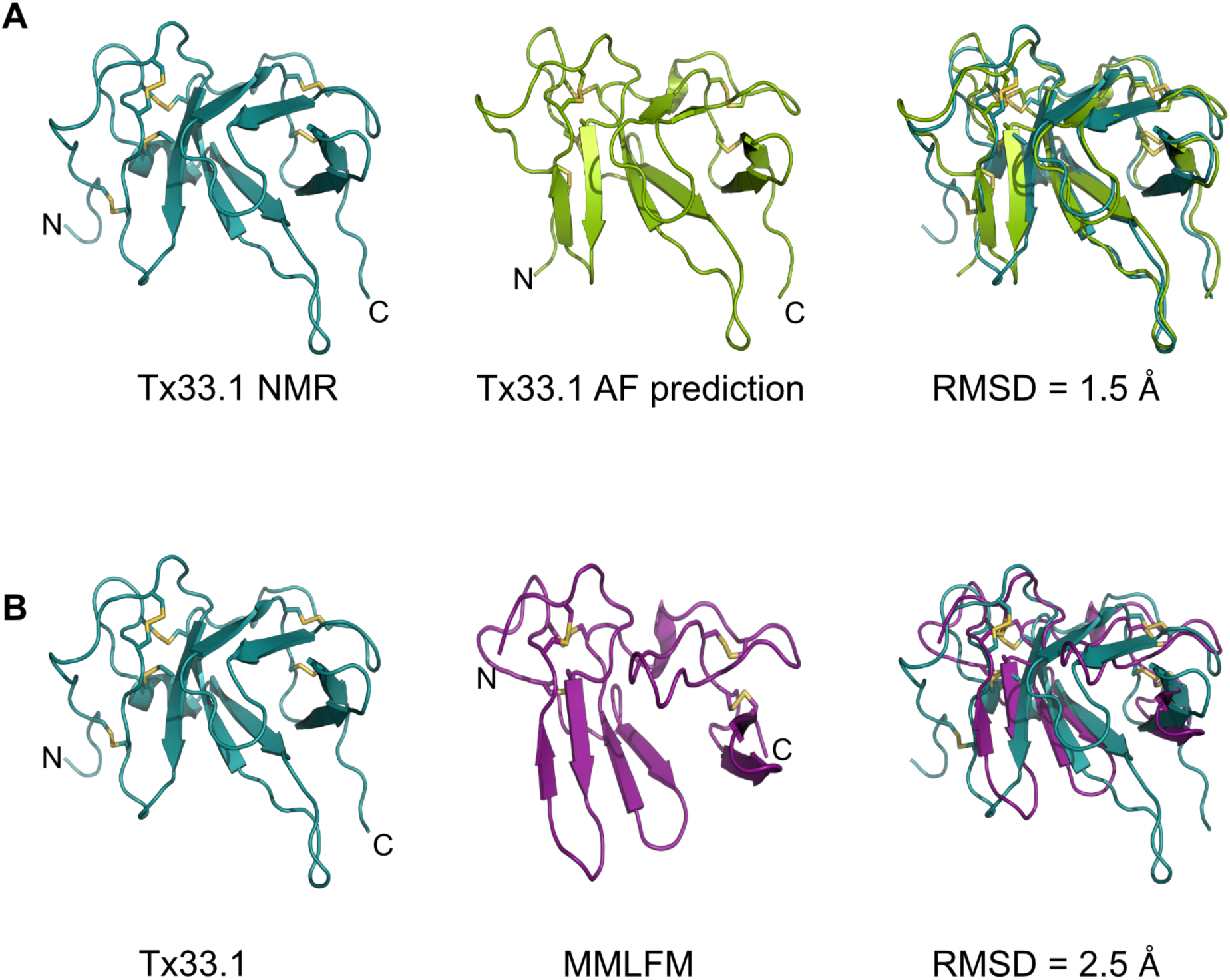
Validation of AlphaFold predictions from structural comparison with the Tx33.1 NMR structure. Structural comparison between the NMR structure of Tx33.1 with the AlphaFold-predicted structures of **(A)** Tx33.1 and **(B)** the MMLFM protein from *Conus coronatus*. Disulfide bonds are shown by yellow sticks and the RMSD values for the two overlays are provided.

Overall, the NMR structure determination of Tx33.1 and the comparison with the AF-predicted structures of proteins from the MMLFM, Unknown, MARFL, unk2, and E superfamilies revealed the details of the structural similarity apparent from Fig. 2B. We found that only the Tx33.1 NMR structure displays a β-barrel structure, while small variations in the orientation of a few residues cause the predicted structures to instead display two domains, each containing two short β-hairpins. Moreover, the long protruding loop between β2 and β3 is a feature exclusively observed in the proteins of the MLSML superfamily, such as Tx33.1.

### Identification of a non-venom cone snail gene encoding a secretory protein with structural similarity to the conotoxins of the six superfamilies

Toxins are often recruited into the venom gland from genes encoding endophysiological proteins. To potentially identify the cone snail gene from which conotoxins of the six investigated superfamilies were originally recruited, we first employed the NMR structure of Tx33.1 in an online structure similarity search using Foldseek (van Kempen, et al. 2024). This search identified a predicted protein with a similar structure in *Pomacea canaliculata*, a freshwater snail for which a reference genome sequence is available (Lu, et al. 2024). Using the amino acid sequence of the Tx33.1 structural homolog from *P. canaliculata*, we employed tblastn to extract the genomic sequences of homologous genes identified in *C. betulinus* and *C. ventricosus*.

The amino acid sequences of these cone snail proteins contain a predicted signal sequence and show the same cysteine pattern as the MMLFM conotoxins, which were consequently selected for further comparison (Fig. 4). The endogenous cone snail protein sequences exhibit substantially greater sequence conservation than the toxin sequences (Fig. 4A). The identified endogenous cone snail proteins are longer by approximately 55 residues than the MMLFM conotoxins. As expected, AF-predicted structures of MMLFM family toxins and the endogenous cone snail proteins exhibit the same overall fold (Fig. 4B), although the β-strands in the conotoxins are generally shorter. The newly identified proteins display three structural domains, each comprising two short β-hairpins (D1, D2, and D3), whereas the conotoxins lack the region encoding D3 (Fig. 4A and B).

**Figure 4.**
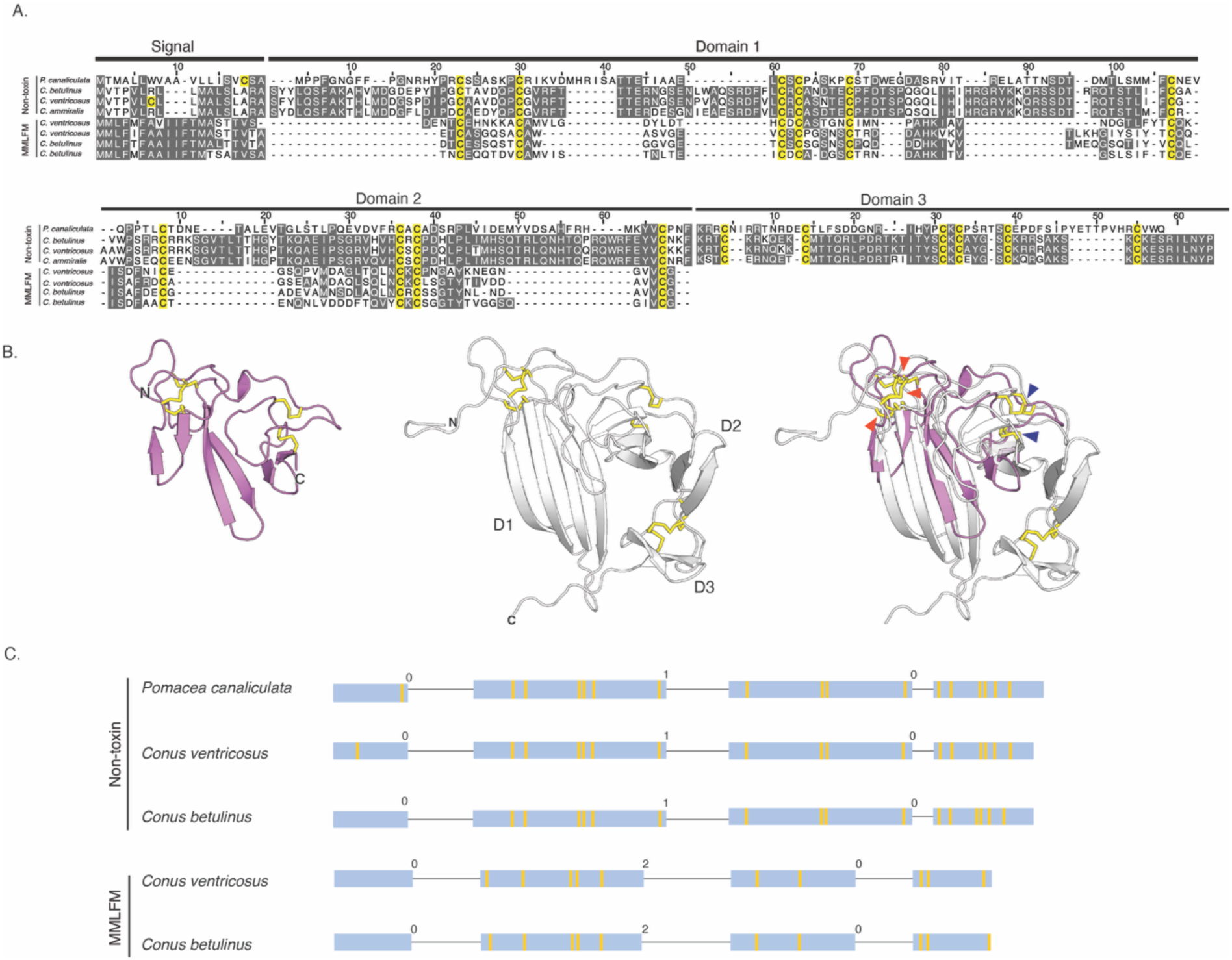
Cone snails encode an endogenous protein with structural similarity to the conotoxins of the six superfamilies. **(A)** Multiple sequence alignment of four non-toxin protein sequences from the freshwater golden apple snail (*P. canaliculata*), cone snails *C. betulinus*, *C. ventricosus* and *C. ammiralis* and four conotoxin protein sequences from the MMLFM superfamily. Domain boundaries were determined based on AlphaFold3 predictions. Alignments were then carried out for each domain individually using MAFFT and visualized in Jalview. Amino acid residues are shaded in gray according to a 30% identity threshold, while all cysteine residues are colored yellow regardless of conservation. The signal sequence and Domains 1 through 3 are delineated by black bars according to the *P. canaliculata* sequence. **(B)** Cartoon representations of the AlphaFold 3 predictions for a *C. betulinus* conotoxin from the MMLFM superfamily (magenta, left) and the non-toxin protein identified from *C. betulinus* (gray, middle). The domains are labelled D1-D3, and disulfide bridges are represented by yellow sticks. In the structural superposition of the two predicted structures (right), positions of structurally equivalent disulfides are marked with arrowheads (orange, D1; dark blue, D2). (**C**) Gene structures of three non-toxin protein sequences from *P. canaliculata*, *C. betulinus*, and *C. ventricosus* compared to the gene structures from conotoxins from the MMLFM superfamily as described in Fig. S7. The exons are depicted as boxes proportional to the length of the sequences, while introns are represented as thin lines (not proportional to sequence length). The intron phases (0, 1, or 2) are given above each intron. The yellow lines within exons indicate cysteine positions.

Both groups of proteins are encoded by four exons (Fig. 4C). However, the two gene structures display different intron phases (“0, 1, 0” *versus* “0, 2, 0”). Closer examination of the gene structures encoding the newly identified proteins reveals that D1 is encoded by the second exon and the beginning of the third, which also encodes the entirety of D2. By contrast, the signal sequence and D3 are each encoded exclusively on separate exons (exon one and exon four, respectively). In contrast, MMLFM conotoxin gene structures show that the signal sequence is exclusively encoded by the first exon, while the mature portion of these proteins is encoded by the three remaining exons, with no clear correlation between intron-exon boundaries and the domain architecture of the protein. The observed dissimilarity in gene structure suggests that the toxin and endogenous protein either do not share a common origin or that their gene structures have diverged significantly since their last common ancestor (see Discussion).

Overall, cone snails encode a predicted secretory protein that is structurally similar in terms of the general fold to the conotoxins of the six investigated superfamilies, but with highly diverged sequences and different phases and positions of the introns.

### Tx33.1 is structurally similar to Argos, a secretory protein from *D. melanogaster*

The discovery of an endogenous cone snail protein and conotoxins displaying similar structural folds prompted us to seek further insight into the prevalence of the observed fold and its associated function. When using the Tx33.1 structure as a query in a Foldseek search, the top hit identified in the Protein Data Bank (PDB) was the crystal structure of Argos, also known as giant lens protein, from *D. melanogaster* (Fig. 5A). Argos is a secretory protein that modulates epidermal growth factor receptor (EGFR) signaling by sequestering the EGF-like ligand called Spitz, thereby regulating diverse developmental processes in *Drosophila* (Freeman, et al. 1992; Klein, et al. 2004). The crystal structure of Argos consists of three separate β-sheet domains (each comprising four strands) of which Domain 2 (D2) and Domain 3 (D3) constitute a clamp-like structure that binds Spitz (Fig. S9) (Klein, et al. 2008). Notably, to enable crystallization, two predicted flexible regions of the Argos sequence were deleted: the first 122 residues and a region of 119 residues located within D1 (Fig. S10). As previously noted (Klein, et al. 2008), the individual domains of Argos adopt the same fold as 3FTXs, such as α-bungarotoxin, with the notable distinction that each domain comprises only two “fingers” (Fig. 5B).

**Figure 5.**
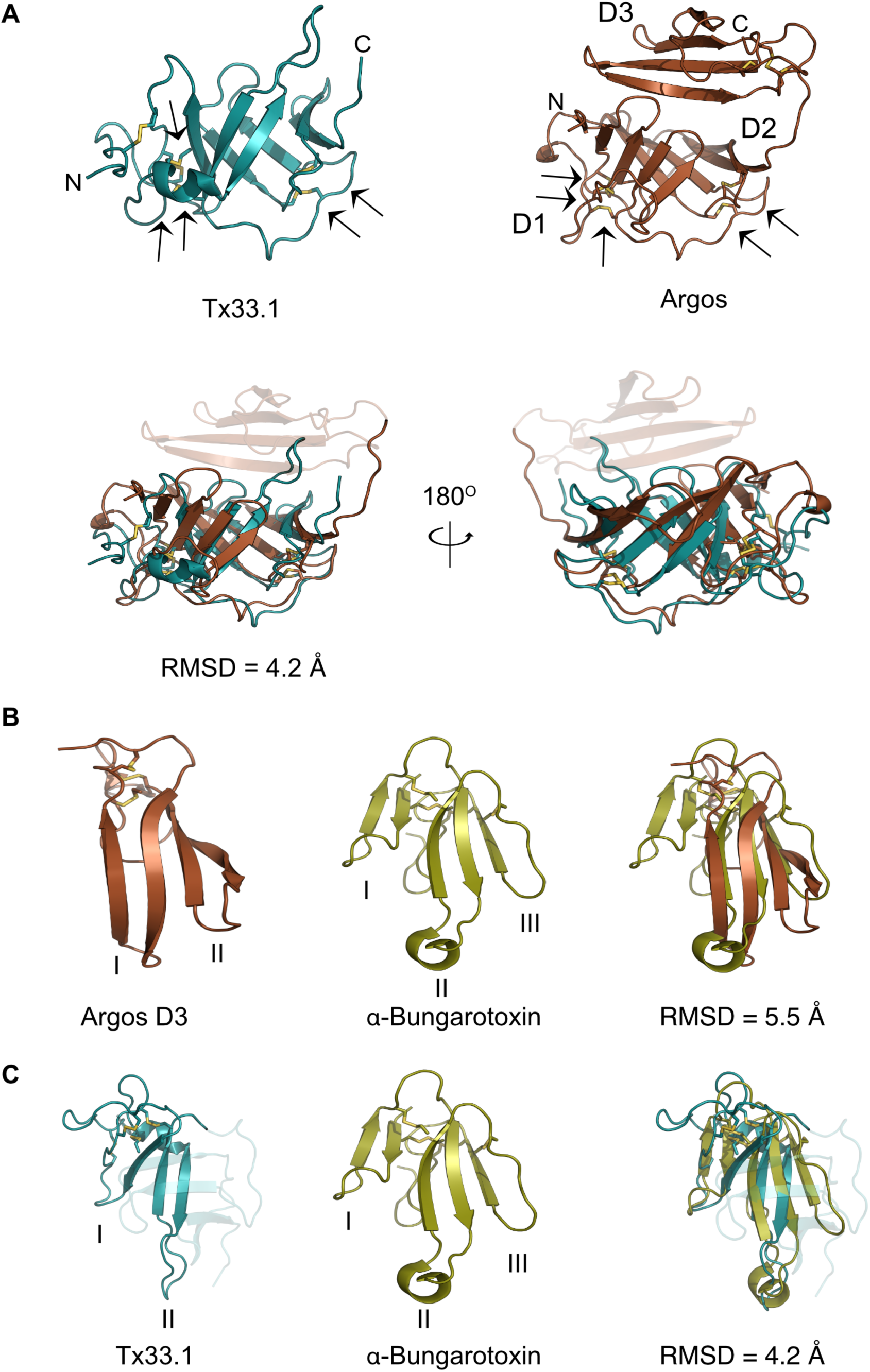
Tx33.1 exhibits a two-finger toxin (2FTX) fold. **A)** Top, left: NMR structure of Tx33.1. Top, right: Crystal structure of Argos (PDB: 3C9A). Arrows point to the five structurally conserved disulfide bonds between the two proteins. D1, D2, and D3 label the three domains in Argos. Bottom: Superimposition of Tx33.1 with the D1-D2 region of Argos shown from two different angles. **(B)** Superimposition of Argos D3 with α-bungarotoxin (PDB: 8DA1). (**C**) Superimposition of the N-terminal region of Tx33.1 (residues 1–71) with α-bungarotoxin. Yellow stick models show disulfide bonds. The roman numerals I–III label the “fingers” in the respective structures.

The structural comparison of Tx33.1 and Argos (Fig. 5A) shows a good structural correlation (RMSD = 4.2 Å) between the D1-D2 region of Argos and Tx33.1. Moreover, the positions of the last five of the six disulfide bonds in Tx33.1 are structurally conserved in Argos (Fig. 5A). The overlay of the region comprising residues 1-71 of Tx33.1 with α-bungarotoxin (Fig. 5C) shows an overall structural similarity with an RMSD = 4.2 Å in the region of the snake toxin comprising the first two fingers. Notably, the position of the extended loop in Finger II of α-bungarotoxin, which constitutes an important binding site for several types of nicotinic acetylcholine receptors (nAChRs) (Zouridakis, et al. 2014; Kessler, et al. 2017), corresponds to the position of the long loop between β2 and β3 in Tx33.1. The regions comprising the N-terminal half of the AF-predicted structures of the MMLFM, Unknown, MARFL, unk2 and E-superfamily toxins also exhibit structural similarity to 3FTXs (not shown). Due to their overall structural similarity to 3FTXs, but with the notable difference of a “missing” finger, we propose the collective name “two-finger toxins” (2FTXs) for the conotoxins of the six superfamilies investigated here.

### Identification of a large group of protostome 2FTX proteins

In addition to detecting Argos, the Foldseek search with Tx33.1 also detected a wide variety of 2FTX structures predicted by AlphaFold. This included venom proteins from centipedes containing three 2FTX domains, as well as proteins from turrids (venomous marine snails that, like cone snails, belong to the superfamily Conoidea) that contain two 2FTX domains (see, e.g., UniProt entries P0DQA2 and A0A098LWA8). Using such proteins as query sequences in PSI-BLAST searching uncovered from a single search approximately 1,800 hits among protostomes, e.g., from mollusks, nematodes, tardigrades, and arthropods (including ticks, mites, spiders, ants, crustaceans, wasps, bees, beetles, butterflies, and scorpions). Manual inspection of these sequences showed that a clear majority contain a predicted signal sequence, indicating that they are secretory proteins like Argos. To the best of our knowledge, none of these many 2FTX proteins have previously been described in the literature and at present their function remains unknown. A single exception is the identification of a male-specific protein from the orb-weaving spider *Tetragnatha versicolor* that exhibits weak sequence similarity to Argos (Zobel-Thropp, et al. 2018).

A CLANS clustering analysis of the sequences obtained from PSI-BLAST searches revealed that they segregate into 12 distinct groups based on sequence similarity (Cluster A-L; Fig. S11A). The AF-predicted structures of a representative sequence from each cluster (Fig. S11B), show that two of the clusters (A and D) encode two-domain 2FTX proteins, while the remaining clusters all encode three-domain 2FTX proteins (see Supporting File 4 for input sequences used in this panel). Not surprisingly, the two clusters that diverge most from the others (K and L) also display structural features rarely observed in the other clusters, such as a greater proportion of loop and α-helix regions.

Argos from *D. melanogaster* is found in Cluster C, which includes proteins similar to Argos with predicted flexible regions in addition to the three 2FTX domains (Fig. S10), as well as proteins comprising only three 2FTX domains (Cluster C; Fig. S11B). In fact, across clusters the most common overall structure consists of three 2FTX domains without intervening predicted flexible regions (Fig. S11B). Sequence comparisons among a selection of such proteins from a small, but diverse set of species revealed conservation of the cysteine residues in a pattern matching that of the MMLFM conotoxins (Fig. S12). Pairwise sequence identities between Argos and the various proteins in the alignment range from 12% to 26% (25%-38% sequence similarity) in the overlapping region. Despite this relatively low sequence identity across different phyla, certain AF-predicted structures of representative proteins closely resembled the structure of Argos (Fig. 6A), exhibiting the same three-domain architecture of the 2FTX fold (Fig. 6B).

**Figure 6.**
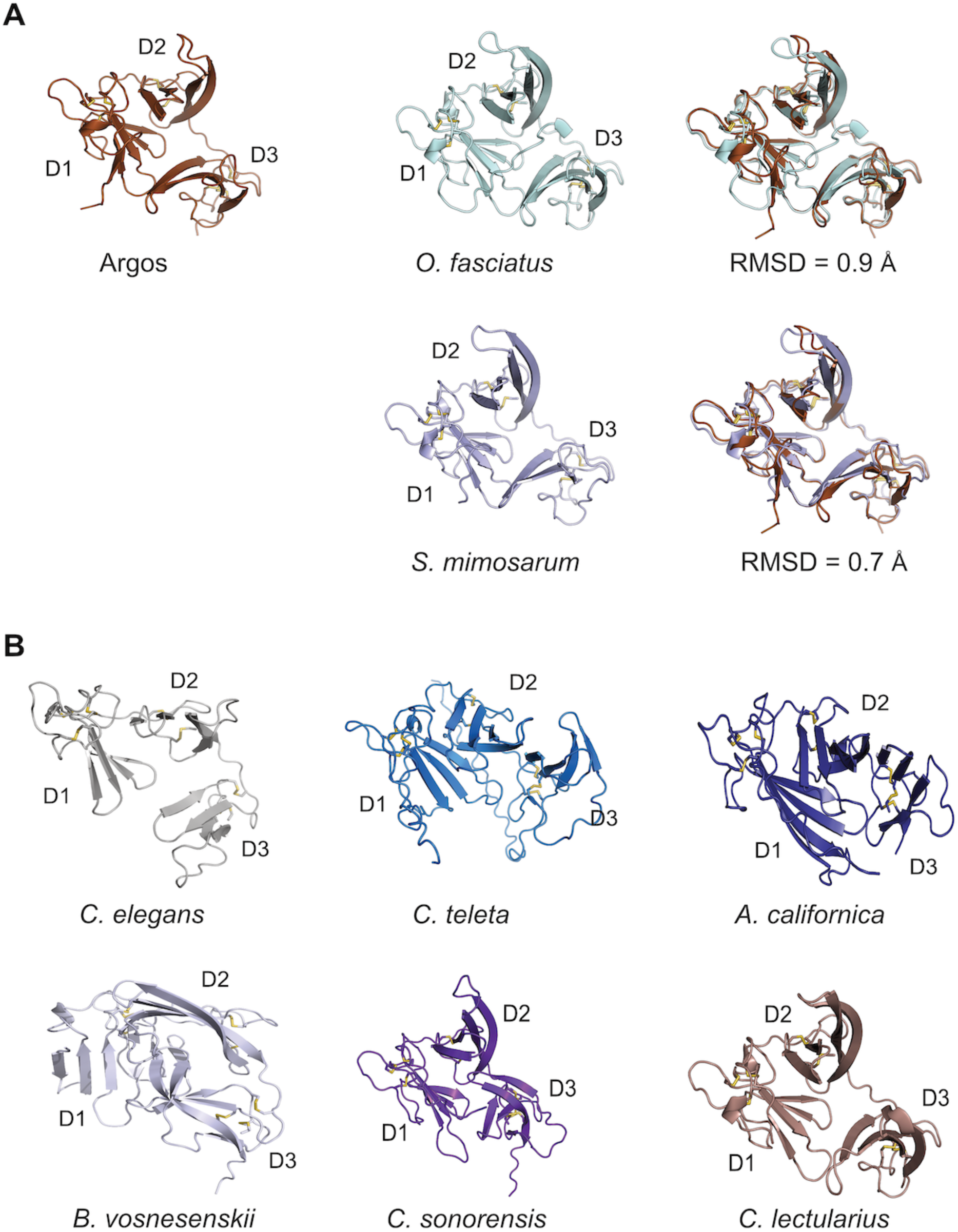
AlphaFold-predicted structures of selected protostome proteins with the same fold as Argos. **(A)** Structural comparison of Argos with Argos-like proteins from *Oncocephalus fasciatus* (slender meadow katydid; A0A2R7VVN0) and *Stegodyphus mimosarum* (African social velvet spider; A0A087V027). **(B)** Examples of AlphaFold-predicted structures of Argos-like protostome proteins from *Caenorhabditis elegans* (fruit fly; Q2157s7), *Capitella teleta* (bristle worm; R7U7Q8), *Aplysia californica* (California sea hare; NCBI-accession XP_005110430.1), *Bombus vosnesenskii* (yellow-faced bumble bee; A0A6J3LNI1), *Culicoides sonorensis* (biting midge; A0A336M5L5), and *Cimex lectularius* (common bed bug; A0A7E4RRV1). Common species names and Uniprot entry numbers are given in parentheses.

Two proteins exhibiting Argos-like structures were identified in model organisms. The *C. elegans* M6.4 protein is of unknown function. According to its Wormbase entry (WBGene00019776), there is enriched tissue expression in the anchor cell and neurons, highest developmental expression in the L1 larval stage, and no phenotype observed for gene knockouts. In *A. californica* we obtained an Argos-like sequence from a Foldseek hit and using this sequence we searched tissue-specific transcriptome datasets from *A. californica* (Koch, et al. 2023) to generate a tissue expression profile (Fig. S13). The results showed the highest expression in pedal ganglia with generally much lower expression in all other tissues.

Overall, these findings reveal a large, previously unrecognized group of protostome-specific 2FTX proteins, several of which show close structural similarity to Argos.

### Investigations of Tx33.1 bioactivity

Finally, we tested bioactivity of Tx33.1 in three experimental systems. First, based on the structural similarity with 3FTXs such as α-bungarotoxin, we tested the activity of Tx33.1 on human α7 and α9α10 nAChRs heterologously expressed in *Xenopus* oocytes using two-electrode voltage-clamp (TEVC) electrophysiology. Tx33.1 showed minimal inhibitory activity at the concentration (30 µM) tested; the ACh-evoked currents amplitudes mediated by α7 nAChRs were 99 +/- 8% (n=4) of control values and 89 +/- 11% (n=4) for α9α10 nAChRs (data not shown).

In a next set of experiments, we assessed the bioactivity of Tx33.1 using the constellation pharmacology assay, which enables monitoring of Ca^2+^ influx in primary cell cultures from mouse dorsal root ganglia (DRG) (Teichert et al, 2012). During each experiment, intracellular calcium levels were simultaneously monitored in ∼1,000 cells using the Fura-2-AM dye. Dissociated DRG neurons were depolarized using 15 s pulses of high-potassium extracellular solution (30 mM), typically resulting in Ca^2+^ elevation, as indicated by an increase in the Fura-2 signal. For activity screening, Tx33.1 was applied in the extracellullar solution (at 1 μM and 10 μM) between KCl pulses, and activity was assessed based on significant changes (block or amplification) in the response to the subsequent KCl pulse. No effects were observed at either concentration tested (data not shown).

Lastly, given the structural resemblance between Tx33.1 and the Argos-like protein identified in *A. californica*, we hypothesized that Tx33.1 might exert a functional effect upon injection into this mollusk. However, we observed no phenotypic effect when assaying gross locomotion in a 30-minute period following injection (data not shown).

## Discussion

Here, we analyzed six previously uncharacterized superfamilies of large conotoxins. Using NMR structure determination and AlphaFold prediction, we demonstrate that these conotoxins exhibit a unique structure distinct from other known toxins. The 2FTX fold identified here is also present in a vast number of protostome proteins, suggesting that the Argos-mediated regulatory mechanism for EGFR signaling may be more widespread in nature than previously recognized.

Cone snails belong to the large superfamily Conoidea, which primarily consists of venomous marine snails, including turrids and terebrids. Based on their prey preference for either fish, worms or other mollusks, cone snails are classified as piscivores, vermivores or molluscivores, respectively. The conotoxins of the six superfamilies investigated here are widely distributed among cone snail species and are expressed in cone snails that prey on fish, worms, and mollusks alike. The presence of several putative toxins most closely related to the MMLFM superfamily in the turrid *Gemmula speciosa* (Watkins, et al. 2006; Gonzales and Saloma 2014) indicates that these toxins were present in the common ancestor of extant cone snails, suggesting their origin within the superfamily Conoidea.

The 2FTX-based structures identified here are reminiscent of the 3FTX fold as previously noted (Klein, et al. 2008). 3FTXs belong to the Ly6/uPAR superfamily of proteins (Kessler, et al. 2017; Leth, et al. 2019; Koludarov, et al. 2023). Several mammalian members of the Ly6/uPAR superfamily are transmembrane proteins belonging to the transforming growth factor (TGF)-β receptor family (Leth, et al. 2019), including the urokinase plasminogen activator receptor and activin type II receptors. Like Argos, these proteins bind growth factor-like domains in their binding partners to exert a variety of cellular functions. Given the predicted structural resemblance to Argos (as illustrated in Fig. 6), we speculate that at least certain protostome Argos-like proteins may modulate EGFR signaling across diverse species. However, because these proteins remain undescribed in the literature, further studies are necessary to verify this hypothesis. The protostome clade encompasses a wide range of organisms across phyla such as arthropods, nematodes, mollusks, flatworms, and annelids, all of which have characterized EGFR signaling pathways (Barberan, et al. 2016). Although it was originally speculated that a human homologue of Argos might exist (Klein, et al. 2008), our PSI-BLAST and FoldSeek searches did not identify any Argos-like proteins in deuterostomes. Given that Argos-binding of Spitz relies on Domain 3, which is absent from the conotoxins of all six superfamilies investigated here, any conotoxin function involving EGF-type ligand binding would require a markedly different binding mode. Therefore, we consider growth factor binding unlikely for the conotoxins in this study. However, we note the existence of a family of EGF-like conotoxins (unpublished observation) and the identification of the kappa-scoloptoxin Ssd1b from the venom gland transcriptome of the centipede, *Scolopendra dehaani*, which encodes an Argos-like protein containing three 2FTX domains (Liu, et al. 2012). Therefore, the predation strategy of at least certain cone snails and centipedes may involve disruption of EGFR signaling in their prey.

Based on the structural similarities with 3FTXs and Argos we tested the bioactivity of Tx33.1 on human α7 and human α9α10 nAChR, on mouse dorsal root ganglion cells, and by injection into *A. californica*. Given the slim chances of target availability for any particular toxin across these diverse systems, the negative outcome of these experiments is not surprising. Consequently, it remains difficult to assess the biological activity of this group of conotoxins at present. Although we did not observe activity on human α7 or α9α10 nAChR, this does not exclude the possibility that these toxins target the cholinergic system, similar to 3FTXs. In any case, for Tx33.1 and other members of the MLSML family, it is tempting to speculate that the distinctive long loop in Finger II plays a functional role, given its close resemblance to the corresponding loop in several long α-neurotoxins that target nAChRs.

Often, weaponization of endophysiological signaling peptides is accompanied by structural adaptations. For instance, 3FTXs lost the transmembrane domain present in their immediate ancestor (Koludarov, et al. 2023), whereas toxins derived from neuropeptides of the CHH family lack an α-helix present in the ancestral genes (Undheim, et al. 2015). Similarly, if the 2FTXs indeed evolved from an endogenous Argos-like protein, these conotoxins may have lost the C-terminal region of their sequences (corresponding to D3 of the cone snail Argos-like protein) during venom recruitment, while simultaneously diverging in sequence. However, due to the differences in intron phases and locations, as well as highly divergent sequences, an evolutionary relationship between the conotoxin families and the endogenous Argos-like protein is not readily apparent.

In conclusion, this study of six uncharacterized families of macro-conotoxins reveals a previously undescribed toxin structure and provides new insight into conotoxin evolution. Our findings also suggest that 2FTX proteins may constitute a large group of overlooked modulators of EGFR signaling in protostomes. Verification of this hypothesis will require further investigation through cellular and organismal studies, such as the examination of EGFR signaling perturbations in null variants of the putative Argos ortholog gene. Methodologically, we propose that combining modern deep learning-based tools for structure prediction, comparison and similarity searching, as applied here, provides a powerful approach for investigating the evolution of rapidly evolving animal venom peptides.

## Materials and methods

### Sequence mining and analysis

We searched published (Koch, et al. 2024) cone snail venom gland transcriptomes using blastp with standard settings. We also used the online blastp and tblastn algorithms to retrieve homologous sequences of Tx33.1 using standard settings. To discern their superfamily assignments, we clustered these using all-against-all blastp p-values in CLANS (Frickey and Lupas 2004). To determine the toxin gene structures, we predicted the coding regions of MLSML, MMLFM, Unknown, MARFL, unk2, and E-superfamily toxins in *C. betulinus, C. ventricosus,* and *C. canariensis* using the protein2genome (p2g) model in Exonerate (v.2.76.2) (Slater and Birney 2005). The annotated sets of known toxins served as queries, and the longest predicted model for each gene was seleceted for further analyses, including identification of intron location and phases.

The amino acid sequence from *P. canaliculata* encoding a protein structurally similar to Tx33.1 was used as a query in a blastp search against the non-redundant protein sequence database within *Conus* to identify orthologues. This search identified a precursor transcriptome sequence from *C. ammiralis*. To identify genomic scaffolds encoding homologous 2FTX proteins, the *C. ammiralis* precursor sequence was then queried against the published genomes of *C. betulinus* and *C. ventricosus* using p2g in Exonerate.

Signal sequences were predicted using SignalP v. 6.0 (Teufel, et al. 2022). Based on the absence of a basic site for proteolytic cleavage before the first cysteine in the sequence, Tx33.1 does not appear to encode a propeptide. The mature sequence is predicted to begin immediately after the signal peptidase cleavage site at Ala18. The predicted mature sequence of Tx33.1 is: KSHTTCPTSTEIDSCSNDNNACGKDVSGSCSSLCNCGNGQTCFTDSNHTITLVPYYTE DGPFEKKYYTCGDPSELDECYDIDKALEVNESDDPNSVEVLCHCPSDKIYLWIHRGY YICITPPQP. Sequences were aligned using the MAFFT server (v. 7) (Katoh, et al. 2002) and visualized using the program Jalview (v. 2.11) (Waterhouse, et al. 2009). To remove sequence redundancy, CD-HIT (Fu, et al. 2012) was used with a 90% identity cut-off.

For the structural similarity analysis in Fig. 2, each of the toxin precursor sequences was predicted using AlphaFold2 (Jumper, et al. 2021) prior to calculating all-against-all structural similarities using Foldseek easy-cluster (Barrio-Hernandez, et al. 2023). The obtained Foldseek e-values were then used as attractive values in the CLANS program. Finally, we reconstructed the phylogeny of the conotoxin superfamilies based on their structural predictions using FoldTree (Moi, et al. 2023), which utilizes Foldseek’s structural alphabet for tree reconstruction.

To determine the tissue distribution of the transcript encoding the Argos-like protein from *A. californica* (NCBI reference XP_005110430.1), we first identified the nucleotide sequence encoding the protein by performing an NCBI tBLASTn search against the *A. californica* transcriptome shotgun assembly database. The identified nucleotide sequence (NCBI reference GBDA01016051.1) was subsequently used as a query sequence in a BLASTn search against various neuronal tissue transcriptomes of *A. californica*, generated as previously described (Koch, et al. 2023). Transcriptomes from two non-neuronal tissues – salivary gland and spermatheca – served as negative controls.

### Selection analyses

The analyses were performed identically for the MLSML, MMLFM, Unknown, MARFL, and E-superfamily genes. For each of the cone snail species expressing these toxin genes, one random gene was selected to avoid comparisons between paralogous genes within the same species. The nucleotide sequences were translated with transeq v6.6.0.0, aligned with mafft v7.525 using the linsi algorithm. We converted the protein sequence alignment to the corresponding nucleotide alignment using pal2nal v14 (Suyama, et al. 2006). The gene tree was estimated using iqtree v2.2.2.3, with the nucleotide models of evolution selected based on the Bayesian Information Criterion as follows: JTTDCMut+G4, WAG+G4, WAG+G4, WAG+G4 for MLSML, MMLFM, Unknown, MARFL and E-superfamily, respectively. Both pairwise dN/dS estimations and site-models were run using paml v4.10.7 with codon frequency FmutSel assigning fitness to every codon.

### Plasmid generation

A codon-optimized sequence for expression in *E. coli* encoding mature Tx33.1 was synthesized and inserted into the pET39_Ub19 expression vector (Rogov, et al. 2012). The resulting plasmid (named pLE666; purchased from TWIST Bioscience^®^) encodes ubiquitin (Ub)-His_10_-tagged Tx33.1 with a cleavage site for TEV protease located immediately before the mature toxin sequence. The fusion protein produced from pLE666 was named Ub-His_10_-Tx33.1.

### Protein expression

Chemically competent *E*. *coli* BL21(DE3) cells were transformed with pLE666 alone or co-transformed with pcsCyDisCo (pLE577) (Nielsen, et al. 2019) encoding Erv1p, hPDI and csPDI to facilitate oxidative folding. The transformed cells were plated on lysogeny broth (LB) agar supplemented with kanamycin (100 µg/ml) (for transformation with pLE666 alone) or kanamycin and chloramphenicol (30 µg/ml) for co-transformations with pLE577. A single colony was transferred into 10 ml LB medium supplemented with the appropriate antibiotic(s) and incubated at 37 °C in an orbital shaker at 200 rpm for ∼16 hours. For small-scale expressions, the pre-culture was inoculated into 50 ml LB medium supplemented with appropriate antibiotic(s) to a starting OD_600_ value of 0.1 and grown at 37°C until the OD_600_ reached ∼0.6. Protein expression was induced by adding isopropyl ß-D-1-thiogalactopyranoside (IPTG) to a final concentration of 1 mM and the culture further propagated at 25°C with shaking (200 rpm) for ∼20 hours. For large-scale expression performed in either 0.5 L (M9 minimal medium) or 1 L (LB medium) culture volume, the overnight culture was diluted in LB or M9 minimal medium (for stable isotope labeling; composition: 3 g/liter KH_2_PO_4_, 15.1 g/liter Na_2_HPO_4_ 12H_2_O, 5 g/liter NaCl, 1 mM MgSO_4_, 1 ml/liter M2 Trace element solution (203 g/liter MgCl_2_ 6H_2_O, 2.1 g/liter CaCl_2_ 2H_2_O, 2.7 g/liter FeSO_4_ 7H_2_O, 20 mg/liter AlCl_3_ 6H_2_O, 10 mg/liter CoSO_4_ 7H_2_O, 2 mg/liter KCr(SO_4_)_2_ 12H_2_O, 2 mg/liter CuCl_2_ 2H_2_O, 1 mg/liter H_3_BO_4_, 20 mg/liter KI, 20 mg/liter MnSO_4_ H_2_O, 1 mg/liter NiSO_4_ 6H_2_O, 4 mg/liter Na_2_MoO_4_ 2H_2_O, 4 mg/liter ZnSO_4_ 7H_2_O, 21 g/liter citric acid monohydrate), 1 g/liter [^15^N]NH_4_Cl, and 4 g/liter [^13^C]glucose) to an OD_600_ = 0.1. For cells grown in LB medium, expression was induced as described above. For isotope labeling, cells were first grown in M9 minimal medium containing unlabelled NH_4_Cl and glucose. Upon reaching OD_600_ ∼ 0.6, cells were pelleted at 3,000 g for 20 minutes and resuspended in 0.5 L M9 medium containing ^15^NH_4_Cl and [^13^C]-glucose. The culture was incubated at 37°C for 30 minutes in an orbital shaker at 200 rpm before induction of protein expression with 1 mM IPTG and overnight incubation at 25°C.

### Cell harvest and preparation of lysate

Bacterial cultures were pelleted by centrifugation at 4,000 g for 15 minutes at 4°C. The cells were resuspended in 20 ml lysis buffer (300 mM NaCl, 50 mM NaH_2_PO_4_, 20 mM imidazole, pH 8.0) per liter of culture, incubated on a roller for 45 minutes at 4°C and frozen at -20°C overnight. After a freeze-thaw cycle, the lysate was sonicated for 8 cycles of 30 seconds with 30-second intervals in between using a UP200S ultrasonic processor (Hielscher) at 90% power while keeping the sample on ice. The lysate was centrifuged at 30,000 g for 45 minutes at 4°C before filtration through a 0.45 μm syringe filter. Cell pellets were resuspended in the same relative volume of lysis buffer containing 8 M urea to be used for SDS-PAGE analysis.

### Protein purification

Ub-His_10_-Tx33.1 was purified from the cleared cell lysate on an ÄKTA Start chromatography system using IMAC on a 5 ml pre-packed HisTrap column (Cytiva^®^). Upon application of the lysate, the column was washed with 20 column volumes of lysis buffer before elution of Ub-His_10_-Tx33.1 with a 0-100% gradient with IMAC elution buffer (300 mM NaCl, 50 mM NaH_2_PO_4_, 400 mM imidazole, pH 8.0). After dialysis into storage buffer (300 mM NaCl, 50 mM NaPi, pH 8.0), the Ub-His_10_-Tx33.1 fusion protein was incubated overnight at room temperature at a ratio of 1:8 with activated His_6_-TEV protease, produced and purified in-house as described previously (Nielsen, et al. 2019). The cleavage mixture was centrifuged to remove any precipitates and applied to a 5 ml gravity flow column packed with Ni-NTA material (QIAGEN^®^). The flow-through containing Tx33.1 was collected, whereas the column-bound Ub-His_10_ tag and His_6_-TEV protease were eluted with IMAC elution buffer. Tx33.1 was then dialyzed into anion exchange wash buffer (20 mM NaCl, 50 mM NaPi, pH = 5.3). The sample was loaded onto an 8 ml Source 15Q (Cytiva^®^) column attached to an ÄKTA Pure System, followed by washing for 3 column volumes with the same buffer. Finally, the protein was eluted using a 30-60 % gradient of anion exchange elution buffer (1 M NaCl, 50 mM NaPi, pH 5.3). The fractions containing pure Tx33.1 as judged by SDS-PAGE analysis were pooled and dialyzed into storage buffer.

### SDS-PAGE analysis

Protein samples were analyzed under non-reducing conditions on 15% glycine SDS-PAGE gels alongside the PageRuler pre-stained protein ladder (Thermofischer Scientific^®^). Gels were stained with Coomassie Brilliant Blue, and images were acquired on a ChemiDoc^TM^ MP Imaging System (BioRad^®^).

### Determination of protein concentration

To determine the concentration of the purified Tx33.1, the absorbance was measured at 280 nm on a NanoDrop spectrophotometer (ND-1000, Thermofischer Scientific^®^). The theoretical extinction coefficient (18,170 M^-1^ cm^-1^) was calculated using the web-based Expasy ProtParam tool (Wilkins, et al. 1999).

### Mass spectrometry

Mass spectrometry was performed on a Waters Xevo G3 QTof system equipped with a ACQUITY UPLC C4 reversed-phase liquid chromatography column (300 Å, 1.7 µm, 2.1 mm × 50 mm) using a sample of purified Tx33.1 at 5 μg/mL adjusted to pH =2 with trifluoroacetic acid as input.

### NMR spectroscopy

A sample of 500 µM ^13^C,^15^N-labelled Tx33.1 were prepared in 10 mM 2-(N-morpholino)ethanesulfonic acid (MES), 50 mM NaCl, 10% D_2_O, pH 6.5. For backbone chemical shift assignments ^15^N-HSQC, HNCA, HN(CO)CA, CBCA(CO)NH, CBCANH, CBCA(CO)NH, HNCO, and HN(CA)CO spectra were recorded on Bruker Avance III HD spectrometers operating at 600 and 750 MHz spectrometer with triple resonance cryoprobes.

Side chain assignments were achieved from HCCONH and HCCH-TOCSY. Distance restraints were obtained from ^15^N-NOESY-HSQC and ^13^C-NOESY-HSQC experiments recorded using a mixing time of 120 ms. The triple resonance spectra were recorded with non-uniform sampling at 25% and reconstructed with qMDD (Kazimierczuk and Orekhov 2011). All spectra were processed with nmrPipe (Delaglio, et al. 1995) and analyzed in CCPNMR analysis (Skinner, et al. 2016).

### NMR structure calculations

Automated NOE assignment was performed using Cyana (Güntert and Buchner 2015). The NOE list was manually refined. XPLOR-NIH (Schwieters, et al. 2003) was used for the final structural refinement, including a torsion angle database potential (Bermejo, et al. 2012) and an implicit solvent model (Tian, et al. 2017). For the final ensemble representing the solution structure of Tx33.1, the 20 structures with the lowest energy were chosen from 100 calculated structures. The Ramachandran-plot statistics was calculated using PROCHECK (Laskowski, et al. 1996) and the coordinate precisions were calculated by MOLMOL (Koradi, et al. 1996). Structure visualization was performed with PyMOL version 2.5.4 (DeLano Scientific). The structure has been deposited in the Protein Data Bank (PDB, wwpdb.org) with PDB code 9FRQ and chemical shifts and peaks lists from the NOESY spectra have been deposited in the Biological Magnetic Resonance Bank (http://www.bmrb.wisc.edu/) with ID 34921.

### Structure prediction and homology searching

Structural predictions using AlphaFold2 or AlphaFold3 were performed with the Google Colab or AlphaFold3 servers (Mirdita, et al. 2022; Abramson, et al. 2024), while structure similarity searches were done using Foldseek (van Kempen, et al. 2024). All AF-models used have pLDDT scores in the range between 0.65 and 0.9, with most above 0.8. Structure superpositions were performed using the “super” command in PyMOL and the listed RMSD values were provided by the same program.

### Electrophysiology

Detailed methods for conducting two-electrode voltage-clamp (TEVC) experiments of human nAChRs heterologously expressed in *X. laevis* oocytes have been previously described (Hone, et al. 2024). Briefly, the oocytes were continuously perfused by gravity at a rate of ∼3 ml/min with frog saline composed of 115 mM NaCl, 2.5 mM KCl, 1.8 mM CaCl_2_, 1.0 mM MgCl_2_, 1 mg/ml bovine serum albumin and buffered to pH 7.4 with 4-(2-hydroxyethyl)-1-piperazineethanesulfonic acid (HEPES). The perfusion system consisted of a series of solenoid valves (NResearch Inc, Caldwell, NJ, USA) controlled by LabVIEW software (National Instruments, Austin TX, USA) operating on a personal computer. To assess the activity of Tx33.1, the oocyte membranes were clamped at a holding potential of -70 mV using a Warner Instruments Oocyte Clamp OC-725C (Warner Instruments, Hamden, CT, USA) and stimulated with 100 µM acetylcholine (ACh) until stable baseline responses were observed. Oocytes expressing the α7 subtype were pulsed with ACh for 500 ms, to minimize open-channel block by ACh, and 1000 ms for those expressing the α9α10 subtype. Once stable responses were observed, Tx33.1, suspended in water, was manually applied to the oocyte in a static bath for 5 min. The ACh-evoked current amplitudes in the presence of Tx33.1 were compared to currents in response to control applications of water. The data were analyzed using Prism 10.2.4 (GraphPad, La Jolla, CA, USA) and the responses expressed as percent response ± SD. The ACh-evoked currents were acquired and sampled at 50 Hz using a USB-6009 digital acquisition system (National Instruments) and filtered at 5 Hz (FIB1; Frequency Devices, Ottawa, IL, USA).

### Calcium imaging

Experiments were performed as previously described (Teichert, et al. 2012). Briefly, lumbar DRG neurons from C57BL/6 mice >30 days old were dissociated by trypsin digestion and mechanical trituration and plated on polylysine-coated plates. The cells were incubated overnight in a 5% CO_2_ incubator at 37 °C in neuronal culture medium [minimal essential media supplemented with 10% (vol/vol) FBS, penicillin (100 U/mL), streptomycin (100 μg/mL), 1× Glutamax, 10 mM HEPES, and 0.4% (wt/vol) glucose, adjusted to pH 7.4]. One hour before the experiment, cells were loaded with 2.5 μM fura-2-acetoxymethyl ester (Fura-2-AM, Sigma-Aldrich) and incubated at 37 °C. Cells loaded with Fura-2-AM were excited intermittently with 340- and 380-nm light; fluorescence emission was monitored at 510 nm. An image was captured at each excitation wavelength, and the ratio of fluorescence intensities (340/380 nm) was acquired every 2 s to monitor the relative changes in intracellular calcium concentration in each cell as a function of time. Typically, ∼1,000 neurons were imaged during each experiment. The calcium transients were elicited by 15 s applications of depolarizing stimulus (30 mM KCl), followed by four washes with extracellular solution [145 mM NaCl, 5 mM KCl, 2 mM CaCl_2_, 1 mM MgCl_2_, 1 mM sodium citrate, 10 mM HEPES, and 10 mM glucose, adjusted to pH 7.4]. For activity screening, Tx33.1 was applied in extracellular solution at 1 and 10 μM between KCl pulses. Four different pharmacological agents were used for cell classification: conotoxin *κ*M-RIIIJ (1 μM), AITC (100 μM), menthol (400 μM) and capsaicin (300 nM). The data was acquired using NIS-Elements and further processed with CellProfiler v.3.1.9 (Jones, et al. 2009). Custom-built scripts in Python v.3.7.2 and R were used for further data analysis and visualization. The experimental methods were approved by the Institutional Animal Care and Use Committee (IACUC) of the University of Utah (Protocol number: 17–05017).

### Aplysia injections

Juvenile (5-20g) *Aplysia californica* purchased from the National Resource for Aplysia, University of Miami, were injected with 10 ng toxin as described previously (Espino, et al. 2024). The center-of-mass was tracked using ZebraZoom (https://github.com/oliviermirat/ZebraZoom; (Mirat, et al. 2013)) and the overall movement was quantified for a 30-minute period.

## Supplementary material

Supporting File 1: All 50 full-length MLSML superfamily sequences. (PDF)

Supporting File 2: All conotoxin amino acid sequences used for the analyses in Fig. 2. (PDF)

Supporting File 3: Source data related to the boxplot shown in Fig. S5. (XLSX)

Supporting File 4: Representative sequences from clusters A-L in Fig. S11A used as input for the AlphaFold predictions in Fig. S11B. (PDF)

## Author contributions

Conceptualization, M.S.K, C.M.H., T.L.K., H.S.-H., and L.E.; Data curation, M.S.K, C.M.H., T.L.K., M.W., K.T., and L.E.; Formal Analysis, M.S.K., T.L.K., A.J.H., and K.T.; Funding acquisition, K.T., B.O., J.M.M., H.S.-H., and L.E.; Investigation, M.S.K, C.M.H., T.L.K., M.W., A.J.H., J.T., A.R., K.T. and L.E.; Project administration, L.E.; Supervision, H.S.-H., K.T. and L.E.; Visualization, M.S.K, C.M.H., T.L.K., K.T., and L.E.; writing—original draft preparation, M.S.K, C.M.H., T.L.K., H.S.-H., K.T. and L.E.; writing—review and editing, all authors. All authors have read and agreed to the published version of the manuscript.

## Footnotes

The atomic coordinates for Tx33.1 are available with the Protein Data Bank under accession number 9FRQ and in the Biological Magnetic Resonance Data Bank under accession number 34921. Nucleotide sequence data for Tx33.1 is available in GenBank under the accession number PV796107.

## Abbreviations

AF: AlphaFold
nAChRs: nicotinic acetylcholine receptors
CD: circular dichroism spectroscopy
CLANS: CLuster ANalysis of Sequences)
csPDI: conotoxin-specific PDI
D: domain
DRG: dorsal root ganglia
DTT: dithiothreitol
EGF: epidermal growth factor
ER: endoplasmic reticulum
HEPES: 4-(2-hydroxyethyl)-1-piperazineethanesulfonic acid
hPDI: human PDI
IMAC: immobilized metal affinity chromatography
IPTG: isopropyl ß-D-1-thiogalactopyranoside
LB: lysogeny broth
MALDI-TOF: matrix-assisted laser desorption– ionization time of flight
MSA: multiple sequence alignment
NCBI: National Center for Biotechnology Information
PDB: Protein Data Bank
PDI: protein disulfide isomerase
RMSD: root mean square deviation
SDS-PAGE: sodium dodecyl sulfate-polyacrylamide gel electrophoresis
TEV: tobacco etch virus
TPM: transcripts per million
Ub: ubiquitin
Ub-His_10_: Ub containing 10 consecutive histidines
2FTX: two-finger toxin
3FTX: three-finger toxin.

## References

Abramson J, Adler J, Dunger J, Evans R, Green T, Pritzel A, Ronneberger O, Willmore L, Ballard AJ, Bambrick J, et al. 2024. Accurate structure prediction of biomolecular interactions with AlphaFold 3. Nature 630:493–500.

Barberan S, Martin-Duran JM, Cebria F. 2016. Evolution of the EGFR pathway in Metazoa and its diversification in the planarian Schmidtea mediterranea. Sci. Rep. 6:28071.

Barrio-Hernandez I, Yeo J, Janes J, Mirdita M, Gilchrist CLM, Wein T, Varadi M, Velankar S, Beltrao P, Steinegger M. 2023. Clustering predicted structures at the scale of the known protein universe. Nature 622:637–645.

Bermejo GA, Clore GM, Schwieters CD. 2012. Smooth statistical torsion angle potential derived from a large conformational database via adaptive kernel density estimation improves the quality of NMR protein structures. Protein Sci. 21:1824–1836.

Bertelsen AB, Hackney CM, Bayer CN, Kjelgaard LD, Rennig M, Christensen B, Sørensen ES, Safavi-Hemami H, Wulff T, Ellgaard L, et al. 2021. DisCoTune: versatile auxiliary plasmids for the production of disulphide-containing proteins and peptides in the E. coli T7 system. Microb. Biotechnol.

Bordon KCF, Cologna CT, Fornari-Baldo EC, Pinheiro-Junior EL, Cerni FA, Amorim FG, Anjolette FAP, Cordeiro FA, Wiezel GA, Cardoso IA, et al. 2020. From Animal Poisons and Venoms to Medicines: Achievements, Challenges and Perspectives in Drug Discovery. Front. Pharmacol. 11:1132.

Degueldre M, Verdenaud M, Legarda G, Minambres R, Zuniga S, Leblanc M, Gilles N, Ducancel F, De Pauw E, Quinton L. 2017. Diversity in sequences, post-translational modifications and expected pharmacological activities of toxins from four Conus species revealed by the combination of cutting-edge proteomics, transcriptomics and bioinformatics. Toxicon 130:116–125.

Delaglio F, Grzesiek S, Vuister GW, Zhu G, Pfeifer J, Bax A. 1995. NMRPipe: a multidimensional spectral processing system based on UNIX pipes. J. Biomol. NMR 6:277–293.

Duda TF, Jr., Palumbi SR. 1999. Molecular genetics of ecological diversification: duplication and rapid evolution of toxin genes of the venomous gastropod Conus. Proc. Natl. Acad. Sci. USA 96:6820–6823.

Espino S, Watkins M, Probst R, Koch TL, Chase K, Imperial J, Robinson SD, Florez Salcedo P, Taylor D, Gajewiak J, et al. 2024. chi-Conotoxins are an Evolutionary Innovation of Mollusk-Hunting Cone Snails as a Counter-Adaptation to Prey Defense. Mol. Biol. Evol. 41.

Fedosov A, Tucci CF, Kantor Y, Farhat S, Puillandre N. 2023. Collaborative Expression: Transcriptomics of Conus virgo Suggests Contribution of Multiple Secretory Glands to Venom Production. J. Mol. Evol. 91:837–853.

Freeman M, Klambt C, Goodman CS, Rubin GM. 1992. The argos gene encodes a diffusible factor that regulates cell fate decisions in the Drosophila eye. Cell 69:963–975.

Frickey T, Lupas A. 2004. CLANS: a Java application for visualizing protein families based on pairwise similarity. Bioinformatics 20:3702–3704.

Fu L, Niu B, Zhu Z, Wu S, Li W. 2012. CD-HIT: accelerated for clustering the next-generation sequencing data. Bioinformatics 28:3150–3152.

Gaciarz A, Khatri NK, Velez-Suberbie ML, Saaranen MJ, Uchida Y, Keshavarz-Moore E, Ruddock LW. 2017. Efficient soluble expression of disulfide bonded proteins in the cytoplasm of Escherichia coli in fed-batch fermentations on chemically defined minimal media. Microb. Cell Fact. 16:108.

Gao B, Peng C, Zhu Y, Sun Y, Zhao T, Huang Y, Shi Q. 2018. High Throughput Identification of Novel Conotoxins from the Vermivorous Oak Cone Snail (Conus quercinus) by Transcriptome Sequencing. Int. J. Mol. Sci. 19.

Gonzales DT, Saloma CP. 2014. A bioinformatics survey for conotoxin-like sequences in three turrid snail venom duct transcriptomes. Toxicon 92:66–74.

Güntert P, Buchner L. 2015. Combined automated NOE assignment and structure calculation with CYANA. J. Biomol. NMR 62:453–471.

Hackney CM, Florez Salcedo P, Müller E, Koch TL, Kjelgaard LD, Watkins M, Zachariassen LG, Tuelung PS, McArthur JR, Adams DJ, et al. 2023. A previously unrecognized superfamily of macro-conotoxins includes an inhibitor of the sensory neuron calcium channel Cav2.3. PLoS Biol. 21:e3002217.

Ho TN, Tran TH, Le HS, Lewis RJ. 2025. Advances in the synthesis and engineering of conotoxins. Eur. J. Med. Chem. 282:117038.

Hone AJ, Santiago U, Harvey PJ, Tekarli B, Gajewiak J, Craik DJ, Camacho CJ, McIntosh JM. 2024. Design, Synthesis, and Structure-Activity Relationships of Novel Peptide Derivatives of the Severe Acute Respiratory Syndrome-Coronavirus-2 Spike-Protein that Potently Inhibit Nicotinic Acetylcholine Receptors. J. Med. Chem. 67:9587–9598.

Illergard K, Ardell DH, Elofsson A. 2009. Structure is three to ten times more conserved than sequence - a study of structural response in protein cores. Proteins 77:499–508.

Jackson TNW, Koludarov I. 2020. How the Toxin got its Toxicity. Front. Pharmacol. 11:574925.

Jones TR, Carpenter AE, Lamprecht MR, Moffat J, Silver SJ, Grenier JK, Castoreno AB, Eggert US, Root DE, Golland P, et al. 2009. Scoring diverse cellular morphologies in image-based screens with iterative feedback and machine learning. Proc. Natl. Acad. Sci. USA 106:1826–1831.

Jumper J, Evans R, Pritzel A, Green T, Figurnov M, Ronneberger O, Tunyasuvunakool K, Bates R, Zidek A, Potapenko A, et al. 2021. Highly accurate protein structure prediction with AlphaFold. Nature 596:583–589.

Katoh K, Misawa K, Kuma K, Miyata T. 2002. MAFFT: a novel method for rapid multiple sequence alignment based on fast Fourier transform. Nucleic Acids Res. 30:3059–3066.

Kazimierczuk K, Orekhov VY. 2011. Accelerated NMR spectroscopy by using compressed sensing. Angew. Chem. Int. Ed. Engl. 50:5556–5559.

Kessler P, Marchot P, Silva M, Servent D. 2017. The three-finger toxin fold: a multifunctional structural scaffold able to modulate cholinergic functions. J. Neurochem. 142 Suppl 2:7–18.

Klein DE, Nappi VM, Reeves GT, Shvartsman SY, Lemmon MA. 2004. Argos inhibits epidermal growth factor receptor signalling by ligand sequestration. Nature 430:1040–1044.

Klein DE, Stayrook SE, Shi F, Narayan K, Lemmon MA. 2008. Structural basis for EGFR ligand sequestration by Argos. Nature 453:1271–1275.

Klint JK, Senff S, Saez NJ, Seshadri R, Lau HY, Bende NS, Undheim EA, Rash LD, Mobli M, King GF. 2013. Production of recombinant disulfide-rich venom peptides for structural and functional analysis via expression in the periplasm of E. coli. PLoS One 8:e63865.

Koch TL, Robinson SD, Salcedo PF, Chase K, Biggs J, Fedosov AE, Yandell M, Olivera BM, Safavi-Hemami H. 2024. Prey Shifts Drive Venom Evolution in Cone Snails. Mol. Biol. Evol. 41.

Koch TL, Torres JP, Baskin RP, Salcedo PF, Chase K, Olivera BM, Safavi-Hemami H. 2023. A toxin-based approach to neuropeptide and peptide hormone discovery. Front. Mol. Neurosci. 16:1176662.

Koludarov I, Senoner T, Jackson TNW, Dashevsky D, Heinzinger M, Aird SD, Rost B. 2023. Domain loss enabled evolution of novel functions in the snake three-finger toxin gene superfamily. Nat. Commun. 14:4861.

Koradi R, Billeter M, Wüthrich K. 1996. MOLMOL: A program for display and analysis of macromolecular structures. J. Mol. Graph. 14:51–55.

Laskowski RA, Rullmannn JA, MacArthur MW, Kaptein R, Thornton JM. 1996. AQUA and PROCHECK-NMR: programs for checking the quality of protein structures solved by NMR. J. Biomol. NMR 8:477–486.

Laugesen SH, Chou DH, Safavi-Hemami H. 2022. Unconventional insulins from predators and pathogens. Nat. Chem. Biol. 18:688–697.

Leth JM, Leth-Espensen KZ, Kristensen KK, Kumari A, Lund Winther AM, Young SG, Ploug M. 2019. Evolution and Medical Significance of LU Domain-Containing Proteins. Int. J. Mol. Sci. 20.

Liu ZC, Zhang R, Zhao F, Chen ZM, Liu HW, Wang YJ, Jiang P, Zhang Y, Wu Y, Ding JP, et al. 2012. Venomic and transcriptomic analysis of centipede Scolopendra subspinipes dehaani. J. Proteome Res. 11:6197–6212.

Lu Y, Luo F, Zhou A, Yi C, Chen H, Li J, Guo Y, Xie Y, Zhang W, Lin D, et al. 2024. Whole-genome sequencing of the invasive golden apple snail Pomacea canaliculata from Asia reveals rapid expansion and adaptive evolution. Gigascience 13.

Mirat O, Sternberg JR, Severi KE, Wyart C. 2013. ZebraZoom: an automated program for high-throughput behavioral analysis and categorization. Front. Neural Circuits 7:107.

Mirdita M, Schutze K, Moriwaki Y, Heo L, Ovchinnikov S, Steinegger M. 2022. ColabFold: making protein folding accessible to all. Nat. Methods 19:679–682.

Moi D, Bernard C, Steinegger M, Nevers Y, Langleib M, Dessimoz C. 2023. Structural phylogenetics unravels the evolutionary diversification of communication systems in gram-positive bacteria and their viruses. bioRxiv:2023.2009.2019.558401.

Mourao CB, Schwartz EF. 2013. Protease inhibitors from marine venomous animals and their counterparts in terrestrial venomous animals. Mar. Drugs 11:2069–2112.

Müller E, Hackney CM, Ellgaard L, Morth JP. 2023. High-resolution crystal structure of the Mu8.1 conotoxin from Conus mucronatus. Acta Crystallogr. F Struct. Biol. Commun. 79:240–246.

Nielsen LD, Foged MM, Albert A, Bertelsen AB, Soltoft CL, Robinson SD, Petersen SV, Purcell AW, Olivera BM, Norton RS, et al. 2019. The three-dimensional structure of an H-superfamily conotoxin reveals a granulin fold arising from a common ICK cysteine framework. J. Biol. Chem. 294:8745–8759.

Peng C, Yao G, Gao BM, Fan CX, Bian C, Wang J, Cao Y, Wen B, Zhu Y, Ruan Z, et al. 2016. High-throughput identification of novel conotoxins from the Chinese tubular cone snail (Conus betulinus) by multi-transcriptome sequencing. Gigascience 5:17.

Raffaelli T, Wilson DT, Dutertre S, Giribaldi J, Vetter I, Robinson SD, Thapa A, Widi A, Loukas A, Daly NL. 2024. Structural analysis of a U-superfamily conotoxin containing a mini-granulin fold: Insights into key features that distinguish between the ICK and granulin folds. J. Biol. Chem. 300:107203.

Rivera-de-Torre E, Rimbault C, Jenkins TP, Sorensen CV, Damsbo A, Saez NJ, Duhoo Y, Hackney CM, Ellgaard L, Laustsen AH. 2021. Strategies for Heterologous Expression, Synthesis, and Purification of Animal Venom Toxins. Front. Bioeng. Biotechnol. 9:811905.

Robinson SD, Li Q, Lu A, Bandyopadhyay PK, Yandell M, Olivera BM, Safavi-Hemami H. 2017. The Venom Repertoire of Conus gloriamaris (Chemnitz, 1777), the Glory of the Sea. Mar Drugs 15.

Robinson SD, Safavi-Hemami H, McIntosh LD, Purcell AW, Norton RS, Papenfuss AT. 2014. Diversity of conotoxin gene superfamilies in the venomous snail, Conus victoriae. PLoS One 9:e87648.

Rogov VV, Rozenknop A, Rogova NY, Lohr F, Tikole S, Jaravine V, Guntert P, Dikic I, Dötsch V. 2012. A universal expression tag for structural and functional studies of proteins. Chembiochem 13:959–963.

Safavi-Hemami H, Foged MM, Ellgaard L. 2018. Evolutionary Adaptations to Cysteine-rich Peptide Folding. In: Feige M, editor. Oxidative Folding of Proteins: Basic Principles, Cellular Regulation and Engineering: Royal Society of Chemistry. p. 99–128.

Safavi-Hemami H, Hu H, Gorasia DG, Bandyopadhyay PK, Veith PD, Young ND, Reynolds EC, Yandell M, Olivera BM, Purcell AW. 2014. Combined proteomic and transcriptomic interrogation of the venom gland of Conus geographus uncovers novel components and functional compartmentalization. Mol. Cell Proteomics 13:938–953.

Safavi-Hemami H, Li Q, Jackson RL, Song AS, Boomsma W, Bandyopadhyay PK, Gruber CW, Purcell AW, Yandell M, Olivera BM, et al. 2016. Rapid expansion of the protein disulfide isomerase gene family facilitates the folding of venom peptides. Proc. Natl. Acad. Sci. USA 113:3227–3232.

Schwieters CD, Kuszewski JJ, Tjandra N, Clore GM. 2003. The Xplor-NIH NMR molecular structure determination package. J. Magn. Reson. 160:65–73.

Skinner SP, Fogh RH, Boucher W, Ragan TJ, Mureddu LG, Vuister GW. 2016. CcpNmr AnalysisAssign: a flexible platform for integrated NMR analysis. J. Biomol. NMR 66:111–124.

Slater GS, Birney E. 2005. Automated generation of heuristics for biological sequence comparison. BMC Bioinformatics 6:31.

Suyama M, Torrents D, Bork P. 2006. PAL2NAL: robust conversion of protein sequence alignments into the corresponding codon alignments. Nucleic Acids Res. 34:W609–612.

Teichert RW, Smith NJ, Raghuraman S, Yoshikami D, Light AR, Olivera BM. 2012. Functional profiling of neurons through cellular neuropharmacology. Proc. Natl. Acad. Sci. USA 109:1388–1395.

Terrat Y, Biass D, Dutertre S, Favreau P, Remm M, Stocklin R, Piquemal D, Ducancel F. 2012. High-resolution picture of a venom gland transcriptome: case study with the marine snail Conus consors. Toxicon 59:34–46.

Teufel F, Almagro Armenteros JJ, Johansen AR, Gislason MH, Pihl SI, Tsirigos KD, Winther O, Brunak S, von Heijne G, Nielsen H. 2022. SignalP 6.0 predicts all five types of signal peptides using protein language models. Nat. Biotechnol. 40:1023–1025.

Tian Y, Schwieters CD, Opella SJ, Marassi FM. 2017. High quality NMR structures: a new force field with implicit water and membrane solvation for Xplor-NIH. J. Biomol. NMR 67:35–49.

Ueberheide BM, Fenyo D, Alewood PF, Chait BT. 2009. Rapid sensitive analysis of cysteine rich peptide venom components. Proc. Natl. Acad. Sci. USA 106:6910–6915.

Undheim EA, Grimm LL, Low CF, Morgenstern D, Herzig V, Zobel-Thropp P, Pineda SS, Habib R, Dziemborowicz S, Fry BG, et al. 2015. Weaponization of a Hormone: Convergent Recruitment of Hyperglycemic Hormone into the Venom of Arthropod Predators. Structure 23:1283–1392.

Undheim EA, Mobli M, King GF. 2016. Toxin structures as evolutionary tools: Using conserved 3D folds to study the evolution of rapidly evolving peptides. Bioessays 38:539–548.

van Kempen M, Kim SS, Tumescheit C, Mirdita M, Lee J, Gilchrist CLM, Soding J, Steinegger M. 2024. Fast and accurate protein structure search with Foldseek. Nat. Biotechnol. 42:243–246.

von Reumont BM, Anderluh G, Antunes A, Ayvazyan N, Beis D, Caliskan F, Crnkovic A, Damm M, Dutertre S, Ellgaard L, et al. 2022. Modern venomics-Current insights, novel methods, and future perspectives in biological and applied animal venom research. Gigascience 11.

Waterhouse AM, Procter JB, Martin DM, Clamp M, Barton GJ. 2009. Jalview Version 2--a multiple sequence alignment editor and analysis workbench. Bioinformatics 25:1189–1191.

Watkins M, Hillyard DR, Olivera BM. 2006. Genes expressed in a turrid venom duct: divergence and similarity to conotoxins. J. Mol. Evol. 62:247–256.

Wilkins MR, Gasteiger E, Bairoch A, Sanchez JC, Williams KL, Appel RD, Hochstrasser DF. 1999. Protein identification and analysis tools in the ExPASy server. Methods Mol. Biol. 112:531–552.

Woodward SR, Cruz LJ, Olivera BM, Hillyard DR. 1990. Constant and hypervariable regions in conotoxin propeptides. EMBO J. 9:1015–1020.

Yang Z. 1997. PAML: a program package for phylogenetic analysis by maximum likelihood. Comput. Appl. Biosci. 13:555–556.

Zobel-Thropp PA, Bulger EA, Cordes MHJ, Binford GJ, Gillespie RG, Brewer MS. 2018. Sexually dimorphic venom proteins in long-jawed orb-weaving spiders (Tetragnatha) comprise novel gene families. PeerJ 6:e4691.

Zouridakis M, Giastas P, Zarkadas E, Chroni-Tzartou D, Bregestovski P, Tzartos SJ. 2014. Crystal structures of free and antagonist-bound states of human alpha9 nicotinic receptor extracellular domain. Nat. Struct. Mol. Biol. 21:976–980.

