## Supplementary figures for "Structural similarities reveal an expansive conotoxin family with a two-finger toxin fold"

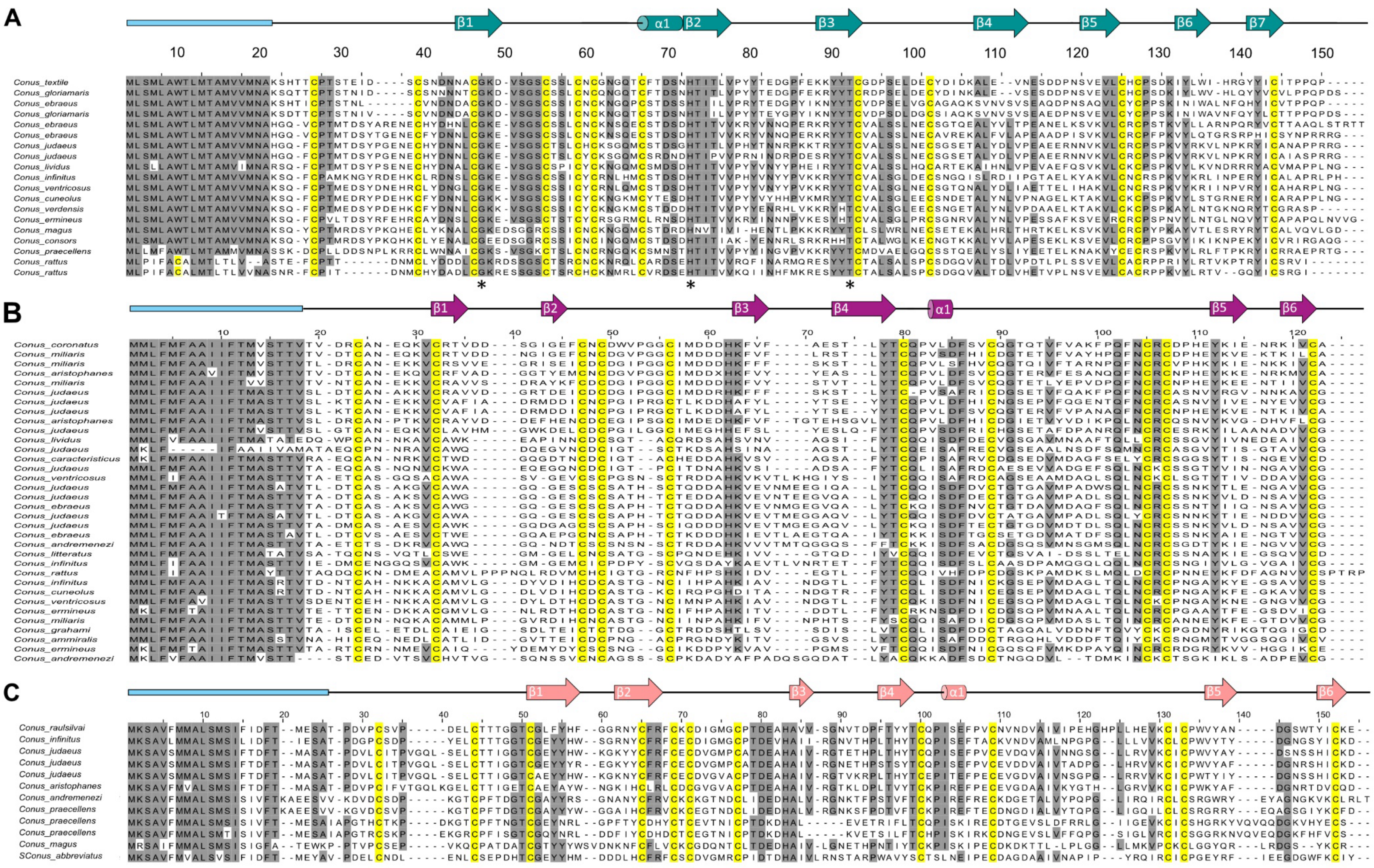
A
β1
α1
β2
β3
β4
β5
β6
β7
B
C

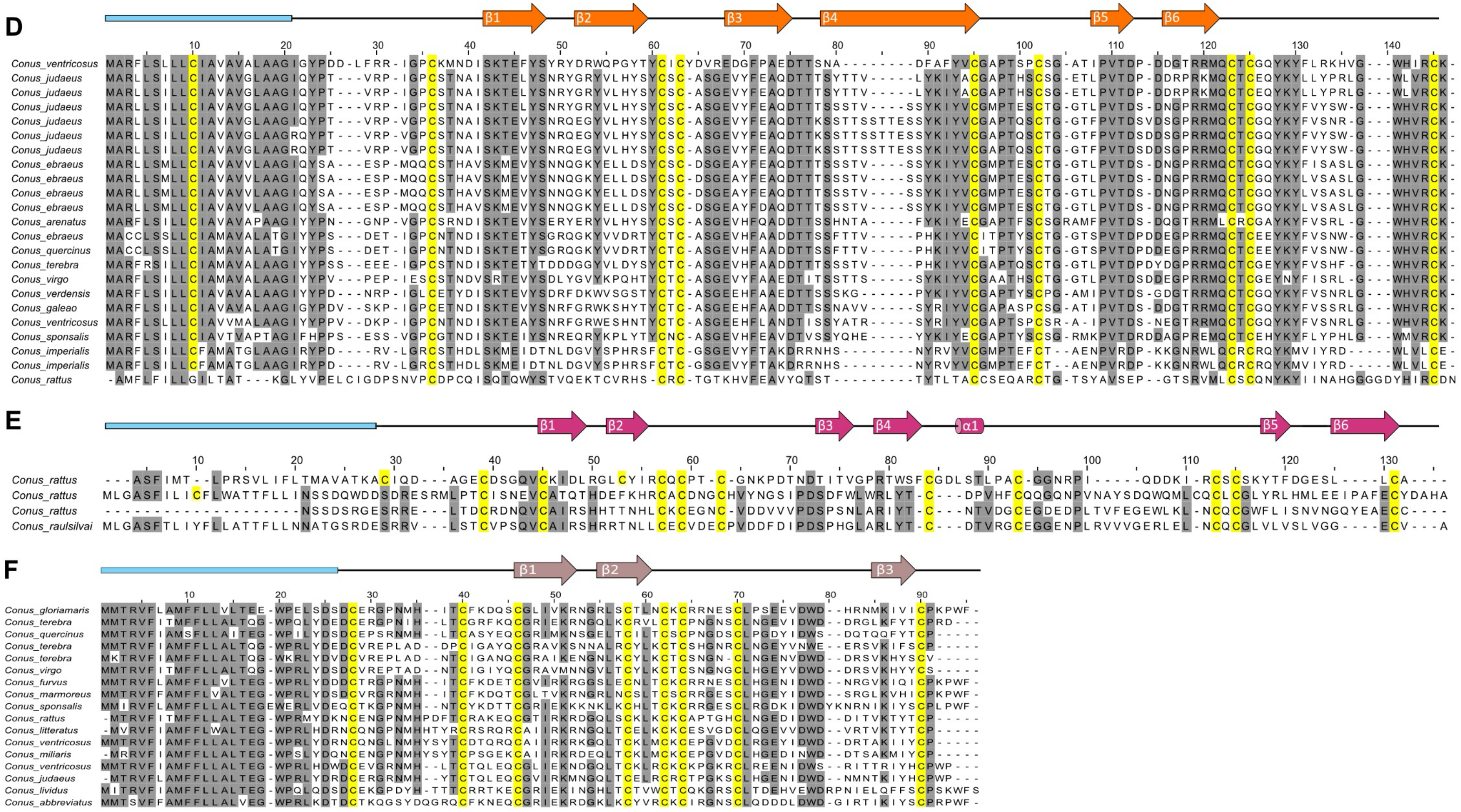
D
β1
β2
β3
β4
β5
β6
Conus_ventricosus
Conus_judaeus
Conus_judaeus
Conus_judaeus
Conus_judaeus
Conus_judaeus
Conus_judaeus
Conus_ebraeus
Conus_ebraeus
Conus_ebraeus
Conus_ebraeus
Conus_arenatus
Conus_ebraeus
Conus_quercinus
Conus_terebra
Conus_virgo
Conus_verdensis
Conus_galeao
Conus_ventricosus
Conus_sponsalis
Conus_imperialis
Conus_imperialis
Conus_rattus
E
β1
β2
β3
β4
α1
β5
β6
Conus_rattus
Conus_rattus
Conus_rattus
Conus_raulsilvai
F
β1
β2
β3
Conus_gloriamaris
Conus_terebra
Conus_quercinus
Conus_terebra
Conus_terebra
Conus_virgo
Conus_furvus
Conus_marmoreus
Conus_sponsalis
Conus_rattus
Conus_litteratus
Conus_ventricosus
Conus_miliaris
Conus_ventricosus
Conus_judaeus
Conus_lividus
Conus_abbreviatus

**Figure S1. Multiple sequence alignments for the six conotoxin superfamilies reveal intra- and interfamily similarities.**

Multiple sequence alignments for proteins of the (**A**) MLSML, **(B)** MMLFM, **(C)** Unknown, **(D)** MARFL, **(E)** unk2, and **(F)** E-superfamilies. The cyan N-terminal rectangle indicates the position of the predicted signal sequences, while the black line marks the mature toxin sequence. Arrows and cylinders above the sequences show the positions of β-strands and an α-helix. Cysteine residues are highlighted in yellow. Residues highlighted in grey indicate at least 70% identity at the given position in the alignment. Aligned sequences were shortlisted from a larger dataset based on percent similarity between the sequences, including only sequences with lower than 90 % similarity in the shown alignments. The asterisks in Panel A indicate strictly conserved residues across the entire family (including the sequences not shown in the alignment).

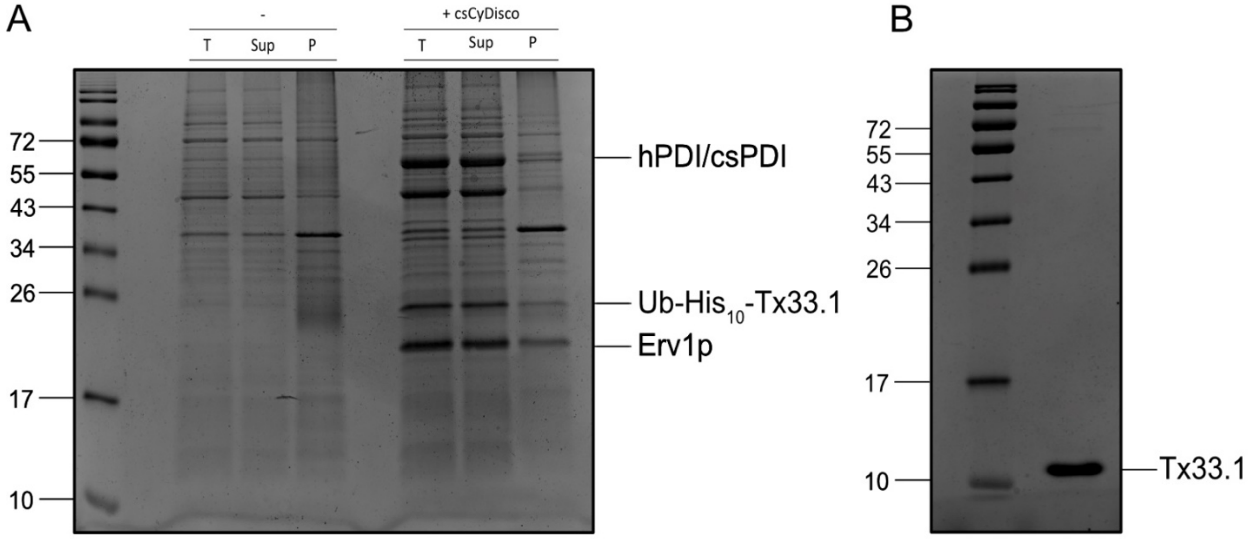


**Figure S2. Tx33.1 is efficiently produced in *E. coli* using the csCyDisCo system.** **(A)** SDS-PAGE analysis of Ub-His$_{10}$-Tx33.1 expressed in *E. coli* BL21(DE3) cells without (-) or with (+ csCyDisCo) the csCyDisCo system. For both expression conditions, samples of the total cell extract (T), the soluble fraction (Sup), and the resuspended pellet after lysis and centrifugation (P) have been loaded. Protein levels are directly comparable between lanes. **(B)** 15% SDS-PAGE gel showing polished Tx33.1 after several steps of purification. Values on the left side of the gels indicate the molecular mass (in kDa) of the indicated marker bands. The gels were stained with Coomassie Brilliant Blue.

**A**

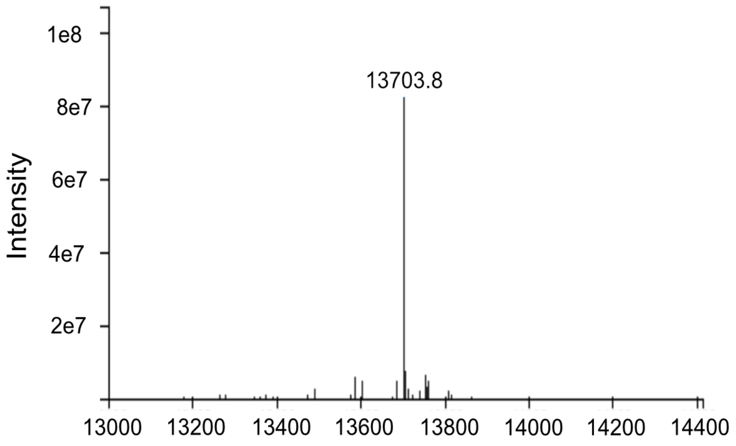


**B**

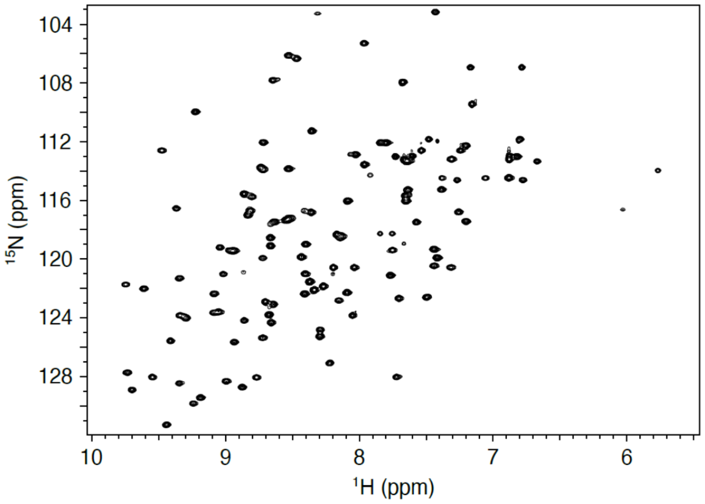


**Figure S3. Purified Tx33.1 is fully oxidized and adopts a folded conformation.**
**(A)** QTOF mass spectrum for Tx33.1 showing a single prominent peak at a monoisotopic mass of 13,703.8 Da. The theoretical monoisotopic mass for Tx33.1 is 13,703.95 Da. **(B)** $^{15}$N-HSQC spectrum for Tx33.1 recorded at 25°C demonstrating well-dispersed signals indicating that Tx33.1 adopts a folded conformation.

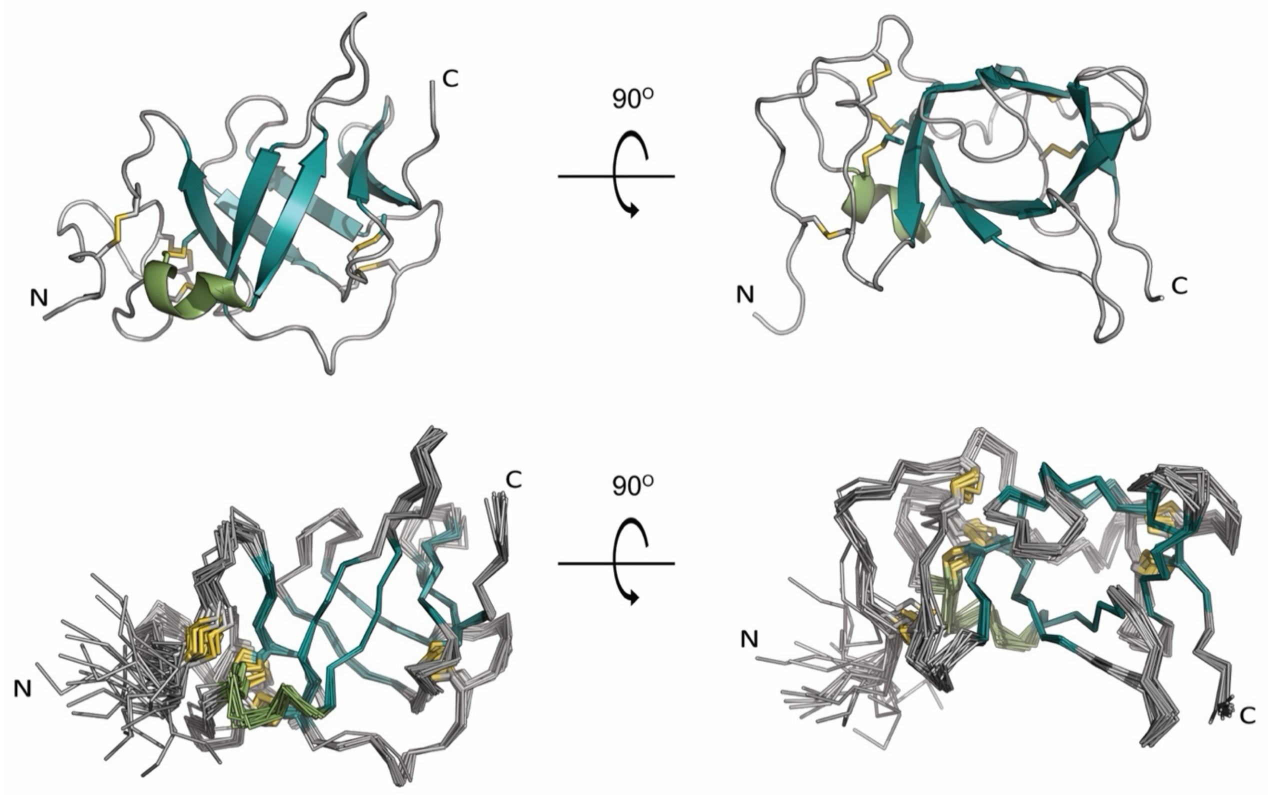


**Figure S4. The 20 lowest energy structures of Tx33.1 reveal a tight bundle.**
Top: Cartoon representation of the NMR structure of Tx33.1 shown from two different angles. Bottom: Superimposition of the 20 lowest energy structures of Tx33.1 shown in the same orientation as above. Cysteine residues are displayed in yellow, the $\alpha$-helix in light green and $\beta$-strands in cyan.

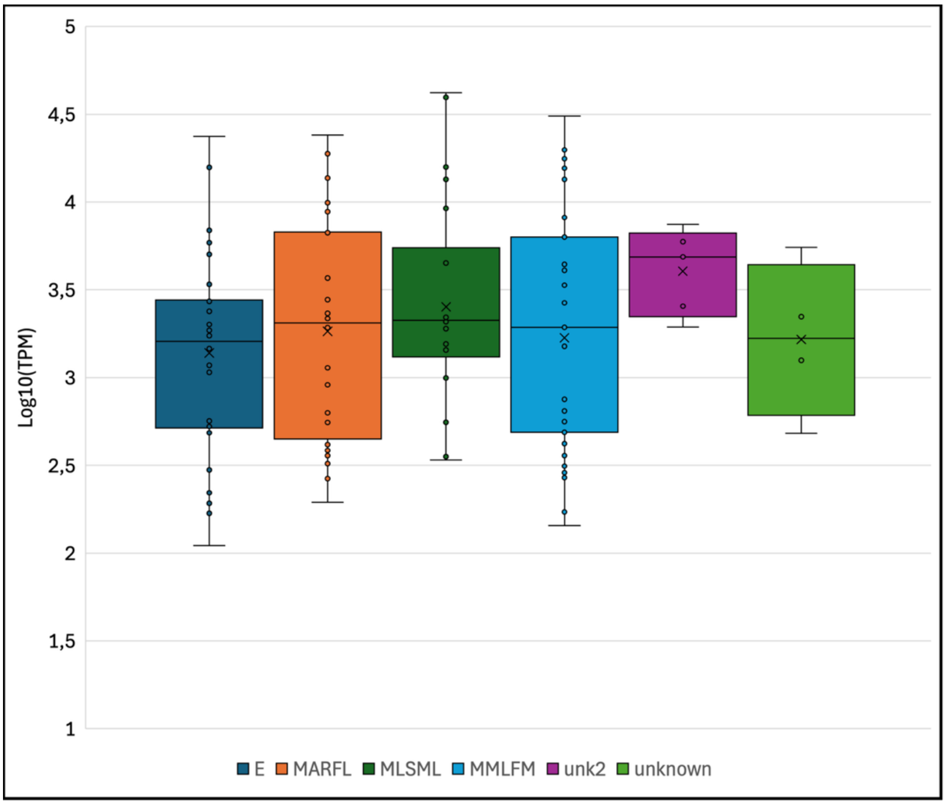


**Figure S5. The six identified conotoxin superfamilies show high transcriptional levels.**

Boxplot of log(10)-transformed transcript per million (TPM) values for transcripts of the E, MARFL, MLSML, MMLFM, unk2 and Unknown superfamilies. Calculations and all sequences analyzed (including their TPM counts) are provided in Supporting File 3.

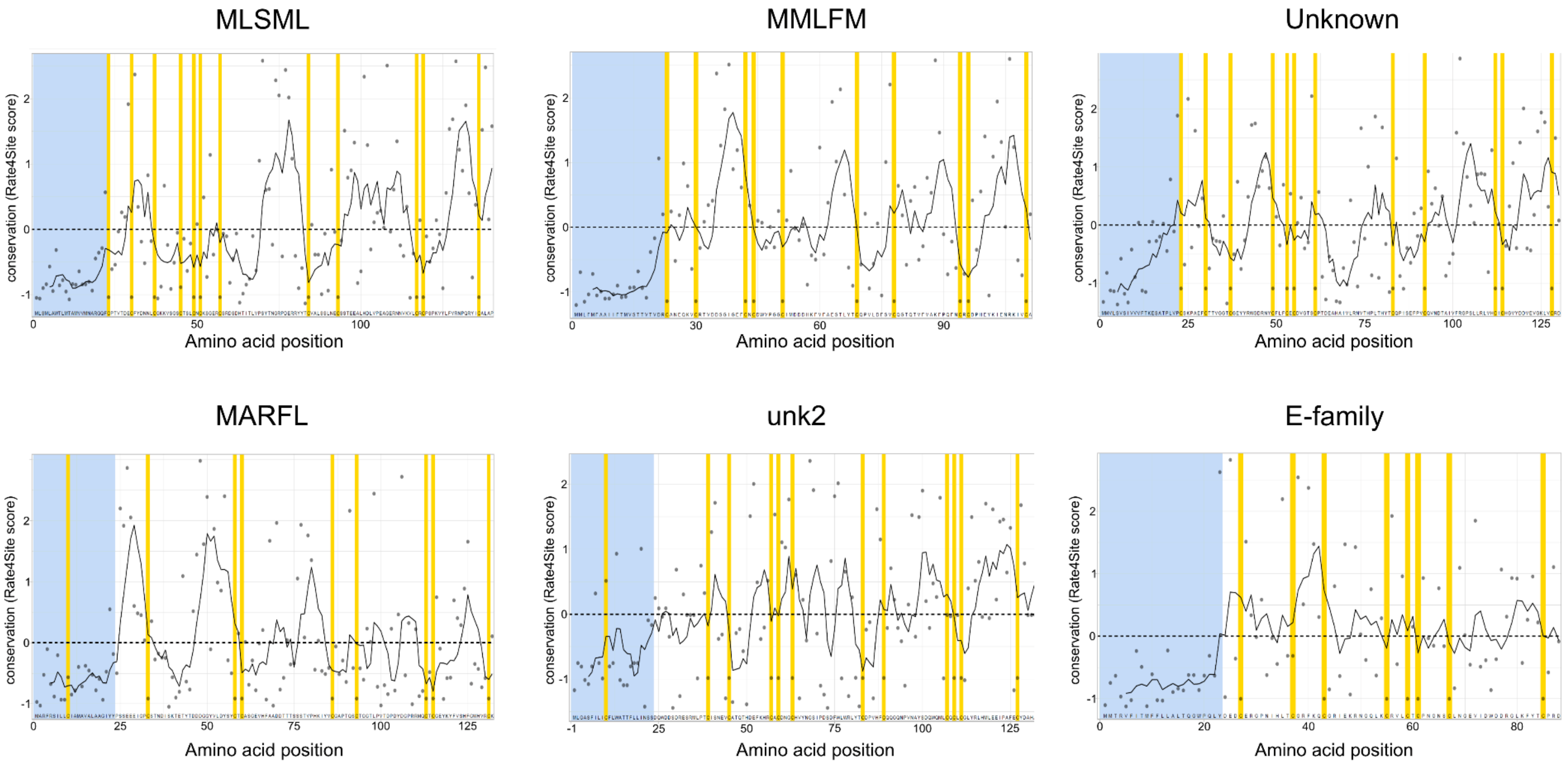


**Figure S6. All six superfamilies of conotoxins show inter-cysteine regions with a high degree of variation.** Position-specific evolutionary rates as determined with Rate4Site of the predicted signal sequence (light blue) and mature (white) toxin regions for the MLSML, MMLFM, Unknown, MARFL, unk2, and E-superfamilies. Using ggplot (Wickham 2016), the Rate4Site score is plotted as a function of the residue position (dots), with high scores indicating greater variation. Cysteine residues are highlighted in yellow. The black curve shows a running average of the five preceding residues.

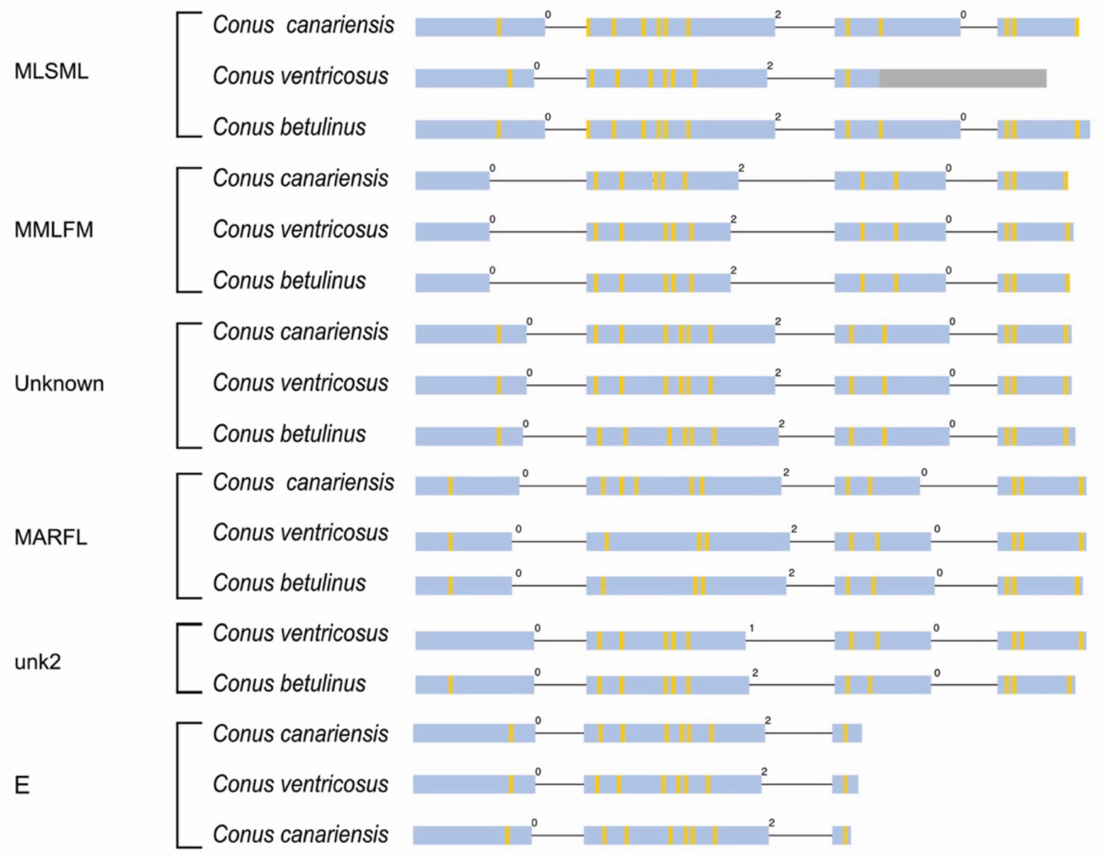


**Figure S7. Gene structures of *C. canariensis*, *C. ventricosus,* and *C. betulinus* conotoxins from the six superfamilies.**

Several identified transcripts from *C. canariensis*, *C. ventricosus* and *C. betulinus* were mapped successfully to their respective genomes. The transcripts were used to assess intron locations and phases from each of the superfamilies. The exons are depicted as boxes proportional to the length of the sequences, while introns are represented as thin lines (not proportional to sequence length). The intron phases (0, 1, or 2) are given above each intron. The yellow lines within exons indicate cysteine positions and the gray color indicates a non-translated region.

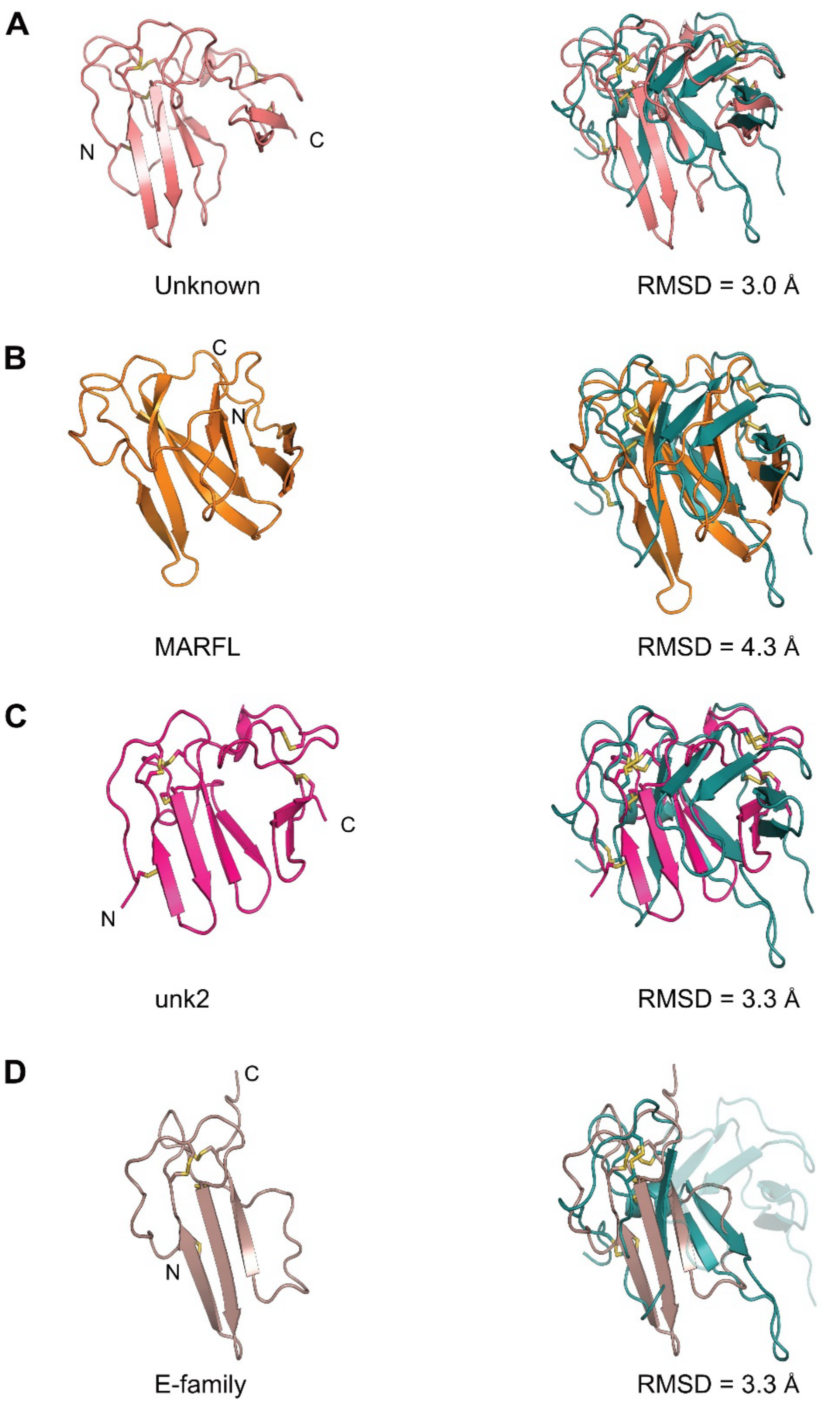


**Figure S8. Tx33.1 displays structural similarity with AlphaFold predictions of proteins from four additional conotoxin superfamilies.**

Structural comparisons of Tx33.1 with the AlphaFold-predicted structures of conotoxins from the (**A**) Unknown (the protein from *Conus raulsilvai*), (**B**) MARFL (the protein from *Conus ventricosus*), (**C**) unk2 (the protein from *Conus rattus*), and (**D**) E-superfamily (the protein from *Conus gloriamaris*). Disulfide bonds are shown by yellow sticks and the RMSD values for the overlays are provided.

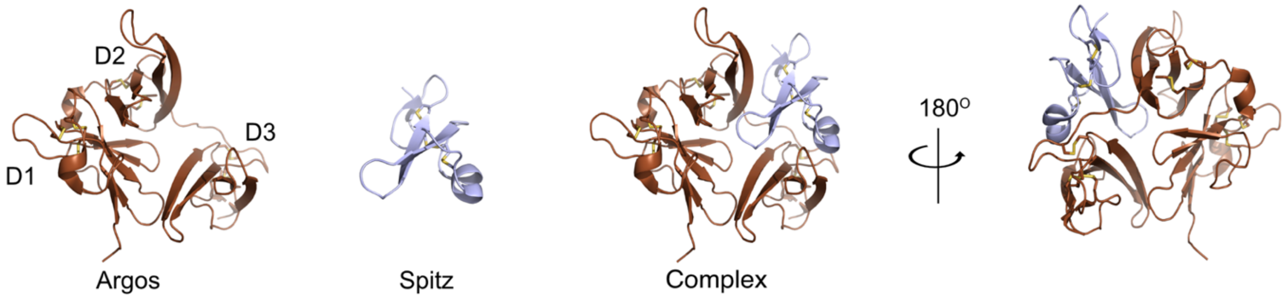


**Figure S9. The crystal structure of Argos in complex with Spitz.**

The crystal structure of Argos (left) in complex with Spitz (middle) (PDB: 3C9A) shown in two different orientations (right). D1, D2, and D3 label the three domains in Argos. Only D2 and D3 make direct contacts with Spitz, while D1 constitutes the backbone of this clamp-like structure.

**A**

RSIIGGKHGDRDVRILYQVGDSEEDLPVCAPNAVCSKIDLYETPWIERQCRCPDGRTCPSSLGVEDGH
TIADKTRHYKMCQPVHKLPVCTHFRDYTWTLTTAAELNVTEQIVHCRCPRNSVTYLTKREPIGNGSPGY
RYLFACSPLTRLRCQRKQPCKLFTVRKRQEFLDEVNINSLCQCPKGHRCPSHHTQSGVIAGESFLEDN
IQTYSGYCMANDHHHHHH

MPTTLMLLPCMLLLLLTAAAVAVGGTRLPLEVFEITPTTSTADKHKSLQYTVVYDAKDISGAAAATGVASS
TVKPATEQLTVVSISSTAAAEKDLAESRRHARQMLQKQQQHRSIIGGKHGDRDVRILYQVGDSEEDLPV
CAPNAVCSKIDLYETPWIERQCRCPESNRMPNNVIIHHHSHSSGSVDSLKYRNYYEREKMMQHKRM
LLGEFQDKKFESLHMKKLMQKLGAVYEDDLDHLDQSPDYNDALPYAEVQDNEFPRGSAHMRHSGH
RGSKEPATTFIGGCPSSLGVEDGHTIADKTRHYKMCQPVHKLPVCTHFRDYTWTLTTAAELNVTEQIVH
CRCPRNSVTYLTKREPIGNGSPGYRYLFACSPLTRLRCQRKQPCKLFTVRKRQEFLDEVNINSLCQCP
KGHRCPSHHTQSGVIAGESFLEDNIQTYSGYCMAND

**B**

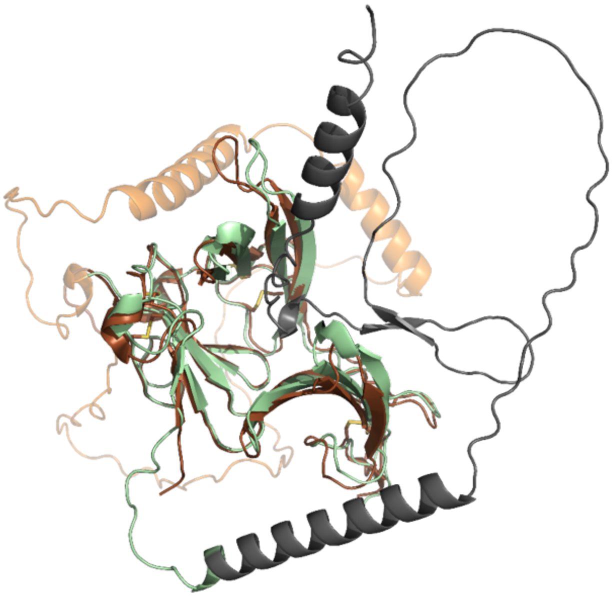

**Figure S10. Comparison between the crystal structure of Argos and the AF-predicted structure of full-length Argos.**

**(A)** Top: The amino acid sequence of the construct used to determine the crystal structure of Argos in complex with Spitz. The DGRT tetrapeptide sequence (white background) placed within D1 originates from *Apis mellifera* Argos. Bottom: The amino acid sequence of full-length Argos. The N-terminal region and the region with D1 not present in the construct used for crystallization are colored black and orange, respectively. **(B)** The crystal structure of Argos from the complex with Spitz (not shown) overlaid with the AF-predicted structure of full-length Argos and shown in the same orientation as used in Fig. S9. The structures are colored according to the color code used in Panel A.

**A**

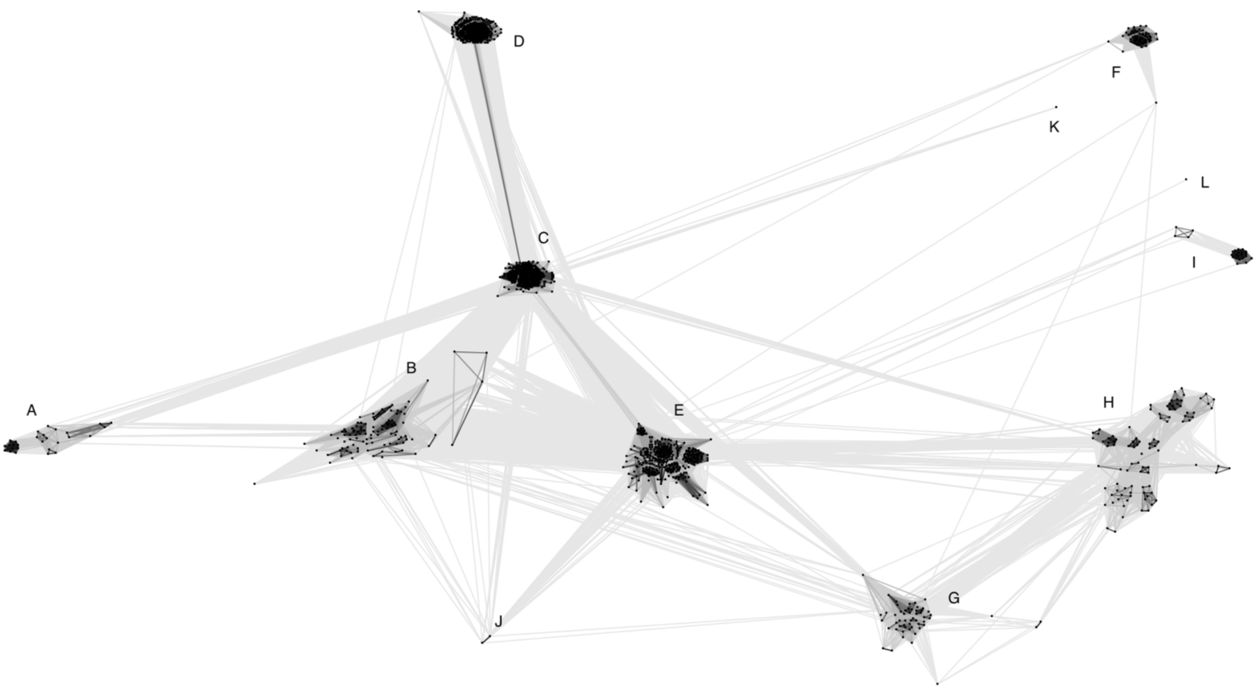


**B**

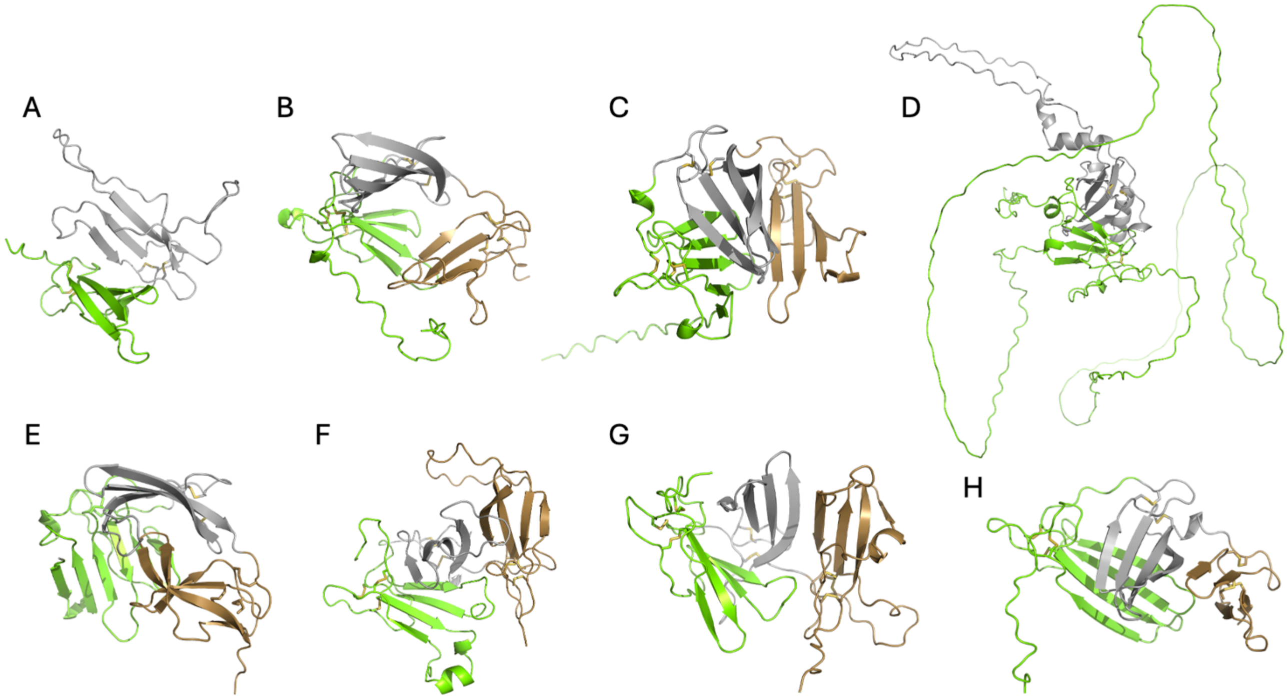

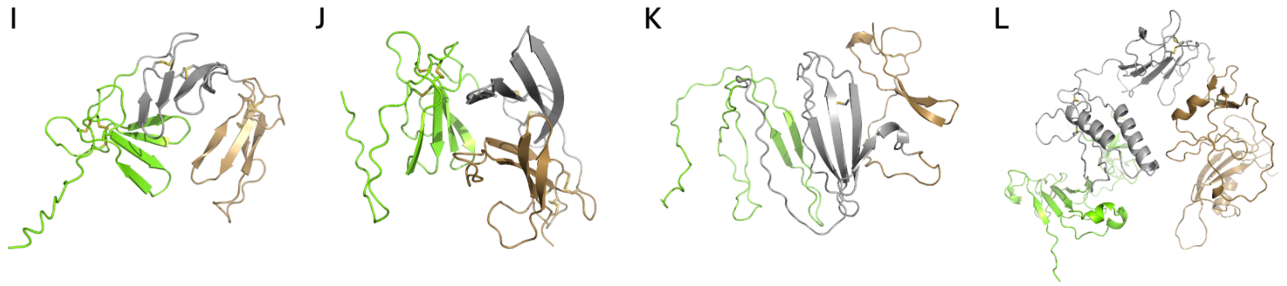


**Figure S11. CLANS clustering analysis and structure predictions for protostome 2FTX proteins.**
**(A)** BLOSUM62 cluster map of protostome 2FTX proteins. The nodes depict individual precursor amino acid sequences, and the edges correspond to the BLAST p-values < 1e-7 between the nodes. The twelve clusters are labelled A-L. **(B)** AlphaFold3 predictions of representative sequences for each of the clusters depicted in (A). Letters denote the different clusters. Regions comprising the first, second and third 2FTX domains are colored in light green, gray and brown, respectively. The sequences used for the predictions in this panel are provided in Supporting File 4 along with the sequence used for PSI-BLAST searching to identify the sequences used for the CLANS analysis in (A).

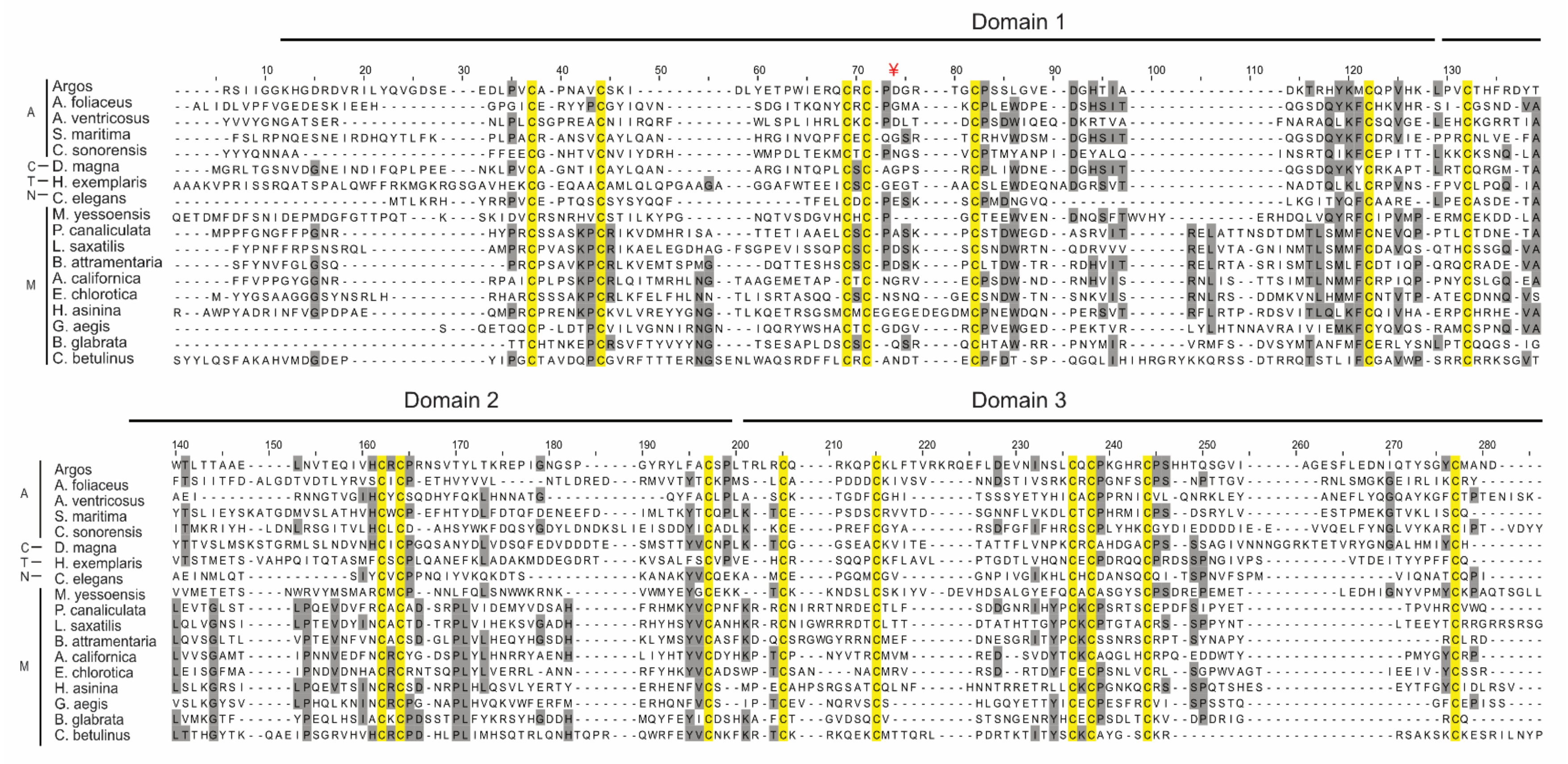


**Figure S12. Multiple sequence alignment of 2FTX proteins from diverse protostomes.**

Multiple sequence alignment of the mature 2FTX sequences from *Drosophila melanogaster* (the Argos protein sequence used for crystal structure determination), *Argulus foliaceus* (common fish louse; A0A7R9RVW3), *Araneus ventricosus* (orbweaver spider; A0A4Y2CDW2), *Strigamia maritima* (coastal centipede; T1J0Y9), *Culicoides sonorensis* (biting midge; A0A336M5L5), *Daphnia magna* (water flea; A0A0P6FFW7), *Hypsibius exemplaris* (water bear; A0A1W0WFR4), *Caenorhabditis elegans* (roundworm; Q21577), *Mizuhopecten yessoensis* (giant ezo scallop; A0A210PEV3) and mollusk species *P. canaliculata* (freshwater golden apple snail; XP 025114045.1), *L. saxatilis* (rough periwinkle snail; XP 070177296.1), *B. attramentaria* (Japanese mud snail; KAK7505081.1), *A. californica* (California sea hare; XP 005110430.1), *E. chlorotica* (green sea slug; RUS85914.1), *H. asinina* (ass's ear abalone; XP 067662371.1), *G. aegis* (giant deep-sea snail; XP 041370382.1), *B. glabrata* (ram's horn snail; XP 055898114.1) and the cone snail *C.*

*betulinus* (betuline cone; JADBJO010000357.1). Amino acid residues are shaded in gray according to a 20% identity threshold, while all cysteine residues are colored yellow. Signal sequences predicted by SignalP 6.0 were removed before alignment, and domain boundaries were determined based on AlphaFold3 predictions. Common species names and Uniprot/NCBI/GeneBank entry numbers are given in parentheses. The red “¥” placed at position 72 in the Argos sequence denotes the site where the loop in domain 1 has been replaced by DGRT for crystal structure determination (see Fig. S10). A = arthropoda, C = crustacean, T = tardigrade, N = nematoda, M = Mollusk.

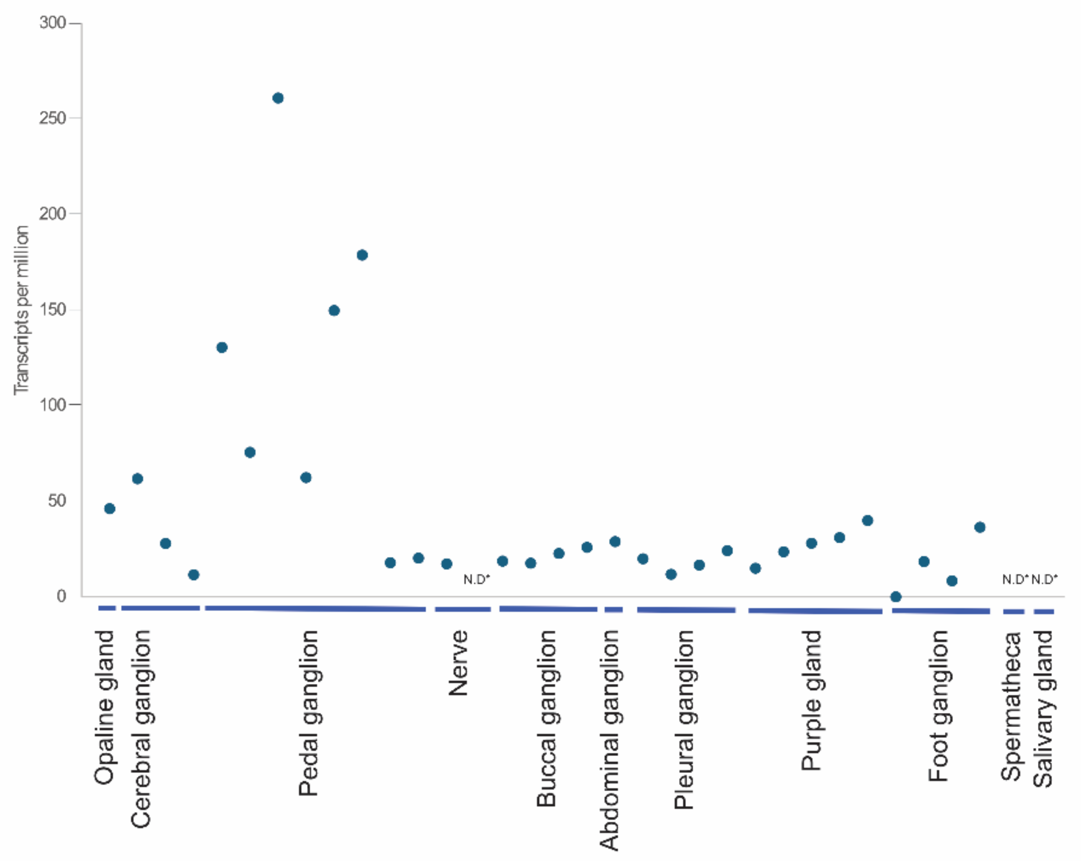


**Figure S13. Tissue distribution of the transcript encoding an *A. californica* Argos-like protein.**

A sequence encoding an Argos-like protein from *A. californica* (NCBI reference GBDA01016051.1) was used as a query sequence for BLASTn searching of various neuronal tissue transcriptomes of *A. californica*, generated as previously described (Koch, et al. 2023), to produce a tissue expression profile. Two non-neuronal tissues – salivary gland and spermatheca – were included as negative controls. N = 1, 2, or 4, depending on the number of tissues present in *A. californica*. Since multiple hits were identified within the same tissue sample for the pedal ganglion and purple gland, the number of hits identified were 6 and 5, respectively. Note the relatively high expression in six pedal ganglion transcriptome datasets compared to other tissues. *ND = not detected.

**References**

Koch TL, Torres JP, Baskin RP, Salcedo PF, Chase K, Olivera BM, Safavi-Hemami H. 2023. A toxin-based approach to neuropeptide and peptide hormone discovery. Front. Mol. Neurosci. 16:1176662.

Wickham H. 2016. ggplot2 : Elegant Graphics for Data Analysis. In. Use R!,. Cham: Springer International Publishing : Imprint: Springer,. p. 1 online resource (XVI, 260 pages 232 illustrations, 140 illustrations in color.
