## Supporting file 1 for "Structural similarities reveal an expansive conotoxin family with a two-finger toxin fold"

**Supporting File 1.** All 50 full-length MLSML superfamily sequences.

>ML.coronatus.TRINITY_DN558_c0_g1_i1_1 supFam_MLSML tpm_556.60 0
MLSMLAWTLMTAMVVMNARGQFCPTVTDECFYDNNLCGKKVSGSCTSLCNCKSGERCSRDSDHTITLVPSYT
NGRPDERRYYTCVALSSLNECSSTEEALHDLVPEAGERNNVKVLCRCPSPKVYLFVRNPQRYICALAP
>ML.ebraeus.TRINITY_DN659_c0_g1_i1_1 supFam_MLSML tpm_41951.58 1
MLSMLAWTLMTAMVVMNAHGQVCPTMTDSYTGENECFYDNNLCGKEVSGSCSSRCYCKNGGRCSTDSDHT
ITVVQSYINGYPEKRYYTCVALSSLNECAVREKALFVLAPEAADPISVKVLCRCPFPKVYLQTGRSRPHICSYNPR
RRG
>ML.ebraeus.TRINITY_DN659_c0_g1_i2_1 supFam_MLSML tpm_39415.85 2
MLSMLAWTLMTAMVVMNAHGQVCPTMTDSYARENECHYDHNLCGKEVSGSCSSLCNCKNSQECSTDSDH
TITVVKRYVNNQPERKRYYTCVALSSLNECSGTQEALYVLTPEANELKSVKVLCRCPSTKVYLLARNPQRYVCTTA
AQLSTRTT
>ML.lividus.TRINITY_DN566_c0_g1_i2_1 supFam_MLSML tpm_1555.75 3
MLSLLAWTLMTAMVIMNARSQFCPTMTDSYPSENECHYDNALCGKEVSGSCSPICYCKNGQMCSMDSDHTI
TVVPYYVNYYPHEIRYYTCVALSSLHQCARTEKAIHNLVPEVAEFQSVKVLCKCPSPKVYLKVNDRRRYACVMAP
PLNG
>ML.lividus.TRINITY_DN566_c0_g1_i3_1 supFam_MLSML tpm_4622.76 4
MLSLLAWTLMTAMVIMNARSQFCPTTTNECHYDNALCGKEVSGSCSPICYCKNGQMCSMDSDHTITVVPYYV
NYYPHEIRYYTCVALSSLHQCARTEKAIHNLVPEVAEFQSVKVLCKCPSPKVYLKVNDRRRYACVMAPPLNG
>ML.rattus.TRINITY_DN825_c0_g1_i1_1 supFam_MLSML tpm_2206.26 5
MLPIFACALMTLTLVASTEFCPTTDNMCLYDDDLCGKRDSSGSCTSRCNCKNRQLCARDSEHTITVVRQFINAR
MQRESYYTCTALSALSPCSDGQVALTDLVPDTLPLSSVEVLCACRPPRIYLRTVKPTRYICSRVI
>ML.rattus.TRINITY_DN825_c0_g1_i3_1 supFam_MLSML tpm_2127.53 6
MLPIFACALMTLTLVVNASNRFCPITDNMCHYDADLCGKRESSGSCTSRCHCKNMRLCVRDSEHTITVVKQIIN
HFMKRESYYTCTALSALSPCSDGQVALTDLVHETVPLNSVEVLCACRPPKVYLRTVGQYICSRGI
>ML.rattus.TRINITY_DN825_c0_g1_i5_1 supFam_MLSML tpm_9202.91 7
MLPIFACALMTLTLVVNASNRFCPITDNMCHYDADLCGKRESSGSCTSRCHCKNMRLCVRDSEHTITVVKQIIN
HFMKRESYYTCTALSALSPCSDGQVALRDLVHETVPLNSVEVLCACRPPKVYLRTVGHYICSTGI
>ML.rattus.TRINITY_DN654_c0_g1_i1_1 supFam_MLSML tpm_2197.66 8
MTLSVIKLTLITMITAVAVTVCSSSDACPNTASEACGLNRHACGKKTPWGTCEHLCRCPNNQPCLRDSKHTFQ
WKRQFLRDKETYYTCNDMSTLPACGNNAGLLHTSGEVTILCECLPPSYYAMESSKYICFRSG
>ML.virgo.TRINITY_DN10252_c0_g1_i1_1 supFam_MLSML tpm_4500.06 9
MLSMLAWTLMTAMVVMNAKSQYCPTMEEDSYPGEHRCFYANDLCGKDVSGSCSSICYCNDGQMCSTDSDH
TITIVPHYVNYFPVRQRYYTCVALSSLHECSRGERAIYDLVPETPELTSVEVLCKCSSPKVYLKVGTARYVCARPPR
LTPSARHLHDDPRRTS
>ML.SRR11807493.C.infinitus.VG.TRINITY_DN59_c0_g2_i3_1 supFam_MLSML tpm_15802.22 10
MLSMLAWTLMTAMVVMNAKSQFCPAMKNGYRDEHKCLRDNSLCGKEVSGSCSSICYCRNLHMCSTDSDHTI
TVVPHYVNYYPVKKRYYTCVALSGLDECSNGQISLRDIIPGTAELKYAKVLCNCRSPKVYKRLINPERYICALAPRL
PH
>ML.SRR11807496.C.cuneolus.VG.TRINITY_DN3685_c0_g1_i1_1 supFam_MLSML tpm_1900.93 12
MLSMLAWTLMTAMVVMNAKSQFCPTMEDRYPDEHKCFYDNNLCGKKVSGSCSSICYCKNGKMCYTESDHTI
TVVPYYVYHHPVKMRYYTCVALSGLEECSNDETALYNLVPNAGELKTAKVLCKCPSPKVYLSTGRNERYICARAP
PLNG
>ML.SRR11807497.C.boavistensis.VG.TRINITY_DN63_c0_g1_i1_1 supFam_MLSML tpm_2077.55 13
MLSMLAWTLMTAMVVMNAKSQFCPAMKNGYRDEHKCLRDNSLCGKEVSGSCSSICYCRNLHMCSTDSDHTI
TVVPHYVNYYPVKKRYYTCVALSGLDECSNGQISLRDIIPGTAELKYAKVLCNCRSPKVYKRLINPERYICALAPRL
PH
>ML.SRR11807498.C.verdensis.VG.TRINITY_DN2612_c0_g2_i1_1 supFam_MLSML tpm_2240.91 14

MLSMLAWTLMTAMVVMNAKSQFCPTMEDSYPDEHKCFYDNNLCGKKVSGSCSSICYCKNGKMCSTDDDHTI
TVVPYYENRHLVKKRYHTCVALSGLEECSNGETALYNLVPDAAELKTAKVLCKCPSPKAYLNTGKNQRYTCGRAS
P
>ML.SRR13740844.C.ventricosus.TRINITY_DN1909_c0_g2_i1_1 supFam_MLSML tpm_16462.67 15
MLSMLAWTLMTAMVVMNAKSQFCPTMEDSYDNEHRCLYDNGLCGKEVSGSCSSICYCRNLQMCSTDSDHTI
TVVPYYVNYYPVKKRYYTCVALSGLNECSGTQNALYDLIAETTELIHAKVLCNCRSPKVYKRIINPVRYICAHARPL
NG
>ML.SRR14407584.C.abbreviatus.TRINITY_DN11_c0_g1_i1_1 supFam_MLSML tpm_994.28 16
MLPIFACALMTLTLVVNASDQFCPTTDNMNQCFYDDALCGKRDSSQGCTSRCYCKNMQLCVRDSEHTITIVKR
IISGHMIKESYYTCTALSALSECDRMQKALEDLVPWTLELDSAKVLCACRPPKIYLRTINPTRYSCFF
>ML.SRR14407587.C.aristophanes.TRINITY_DN475_c0_g1_i1_1 supFam_MLSML tpm_339.89 17
MLSMLAWTLMTAMVVMNAHGQFCPTMTDECFYDNNLCGKKVSGSCTSLCNCKSGERCSRDSDHTITLVPSY
TNGRPDQRRYYTCVALSSLNECSSTEEALYDLVPEAGERNNVKVLCRCPSPKVYLFVFNPQRYICALAP
>ML.SRR15402271.C.textile.TRINITY_DN602_c0_g1_i1_1 supFam_MLSML tpm_2188.42 18
MLSMLAWTLMTAMVVMNAKSHTTCPTSTEIDSCSNDNNACGKDVSGSCSSLCNCGNGQTCFTDSNHTITLVP
YYTEDGPFEKKYYTCGDPSELDECYDIDKALEVNESDDPNSVEVLCHCPSDKIYLWIHRGYYICITPPQP
>ML.SRR15402271.C.textile.TRINITY_DN602_c0_g2_i4_1 supFam_MLSML tpm_1992.39 19
MLSMLAWTLMTAMVVMNAKSHTICPTSTNLCVNDNDACGKDVSGSCSSICNCKNGQPCSTDSSHTIILVPRYT
EDGPYIKNYYTCVDPSELVGCAGAQKSVNVSVSEAQDPNSAQVLCYCPPSKINIWALNFQHYICVTPPQP
>ML.SRR1544627.C.miliaris.TRINITY_DN3169_c0_g1_i1_1 supFam_MLSML tpm_511.66 20
MLSMFAWTLMTAMVVMNAHGQFCPTMTDINYPGENRCFYDNNLCGKEDSGSCTSLCYCKSGQMCSRDTD
HTITLVPRITNSGPDERRYYTCVALSSLNECSGTEKALHILSPEAEEPISVEVLCRCPSKVYRFILYPRPQRYICAPA
S
>ML.SRR17653514.C.judaeus.TRINITY_DN505_c0_g1_i2_1 supFam_MLSML tpm_1597.20 21
MLSMLAWTLMTAMVVMNAHGQFCPTMTDSRPGENKCLHDNYLCGKEDDSGSCTSLCYCKSGQMCFRDND
HTITAVPRIINDRPVEITYYTCVALSSLNECSGIQEALKVLSPEAEEGESSVEVLCRCPSPKKYRFVFIRPDINGDNP
RYVCANAPRRHG
>ML.SRR17653514.C.judaeus.TRINITY_DN740_c0_g1_i3_1 supFam_MLSML tpm_2116.13 22
MLSVFTVVWVLTTVMMMTDVTFQSTCDTDNLELCSEATHMCGKRISWDGCNGLCKCRTLQACTTDADHTVQ
VIPAPFQSNKTYYTCRSLSTMGACQSNNEAMSGNSEDTYKILCKCDETYQPHSLNNRTFVCR
>ML.SRR2124878.C.betulinus.TRINITY_DN401_c0_g1_i2_1 supFam_MLSML tpm_13467.75 23
MLSVFTVVWVLTTAMMMTDVTLQSTCDTDDLGLCSEDTRLCGKRTVWNRCNGLCKCPNQQACTTDTDHTVR
VRSAPFQLIQTYYTCRDVSTMDACQSNERAMDGHNEETYKILCKCDNIYQPNAPQNWYFICS
>ML.consors.TRINITY_DN5487_c0_g2_i1_1 supFam_MLSML tpm_360.65 24
MLSMLAWTLMTAMVVMNAKSQYCPTMRDSYPKQHKCLYENALCGEEDSGGRCSSICNCINGQMCSTDSDH
TITIAKYENNRLSRKRHHTCTALWGLKECNGTKKALYVLAPESEKLKSVEVLCRCPPSGVYIKIKNPEKYICVRIRG
AQG
>ML.MLSML-Gm1_gloriamaris.TRINITY_DN3174_c0_g1_i2_1 supFam_MLSML tpm_354.33 25
MLSMLAWTLMTAMVVMNAKSDTTCPTSTNIVSCVNDNDACGKDVSGSCSSICNCKNGQTCSTDSNHTIILVPY
YTEYGPYIKNYYTCVDPSDLDGCSIAQKSVNVSVSEAEDPNSVEVLCYCPPSKINIWAVNFQYYLCTTPPQPDS
>ML.MLSML-Gm2_gloriamaris.TRINITY_DN3174_c0_g1_i3_1 supFam_MLSML tpm_365.71 26
MLSMLAWTLMTAMVVMNAKSQTTCPTSTNIDSCSNNNNTCGKDVSGSCSSLCNCQNGQTCFTDSNHTITLV
PYYTEDGPFEKRYYTCRDPSELDECYDINKALEVSESDDPNSVEVLCHCPSDKIYLWVHLQYYVCVLPPQPDS
>ML.DAZ86972.1 TPA_inf: conotoxin precursor Tpra06 [Conus judaeus] 27
MLSMLAWTLMTAMVVMNAHGQFCPTMTDSYPGENECHYDNNLCGKEVSGSCTSLCHCKSGQMCSTDSGH
TITLVPHYTNNRPKKTRYYTCVALSSLNECSGSETALYDLVPEAEERNNVKVLCRCPSPKVYRKVLNPKRYICANA
PPRRG
>ML.UMA83964.1 conotoxin precursor Tpra06 [Conus judaeus] 28
MLSMLAWTLMTAMVVMNAHGQFCPTMTDSYPGENECHYDNNLCGKEVSGSCTSLCYCKSGQMCSTDSGH
TITLVPHYTNNRPKKTRYYTCVALSSLNECSGSETALYDLVPEAEERNNVKVLCRCPSPKVYRKVLNPKRYICAIAS
PRRG
>ML.DAZ86971.1 TPA_inf: conotoxin precursor Tpra06 [Conus judaeus] 29

MLSMLAWTLMTAMVVMNAHGQFCPTMTDSYPGENECHYDNNLCGKEVSGSCTSLCHCKSGQMCSRDND
HTIPVVPRNINDRPDESRYYTCVALSSLNECSGSETALYDLVPEAEERNNVKVLCRCPSPKVYRKVLNPKRYICA
NAPPRRG
>ML.UMA83620.1 conotoxin precursor Tpra06 [Conus judaeus] 30
MLSMLAWTLMTAMVVMNAHGQFCPTMTDSYPGENECHYDNNLCGKEVSGSCTSLCYCKSGQMCSRDNDH
TIPVVPRNINDRPDESRYYTCVALSSLNECSGSETALYDLVPEAEERNNVKVLCRCPSPKVYRKVLNPKRYICAN
APPRRG
>ML.UMA83315.1 conotoxin precursor Tpra06 [Conus judaeus] 31
MLSMLAWTLMTAMVVMNAHGQFCPTMTDSYPGENECHYDNNLCGKEVSGSCTSLCYCKSGQMCSRDNDH
TIPVVPRNINDRPDESRYYTCVALSSLNECSGSETALYDLVPEAEERNNVKVLCRCPSPKVYRKVLNPKRYICAIA
SPRRG
>ML.UMA82655.1 conotoxin precursor Tpra06 [Conus ebraeus] 32
MLSMLAWTLMTAMVVMNAHGQVCPTMTDSYTGENECFYDNNLCGKEVSGSCSSLCNCKNSQECSTDSDHT
ITVVKRYVNNQPERKRYYTCVALSSLNECSGTQEALYVLTPEANELKSVKVLCRCPSTKVYLLARNPQRYVCTTAA
PLSTRTT
>ML.UMA82382.1 conotoxin precursor Tpra06 [Conus ebraeus] 33
MLSMLAWTLMTAMVVMNAHGQVCPTMTDSYARENECHYDHNLCGKEVSGSCSSLCNCKNSQECSTDSDH
TITVVKRYVNNQPERKRYYTCVALSSLNECSGTQEALYVLTPEANELKSVKVLCRCPSTKVYLLARNPQRYVCTTA
APLSTRTT
>ML.UMA82383.1 conotoxin precursor Tpra06 [Conus ebraeus] 34
MLAWTSMTAMVVMNAHGQVCPTMTDSYARENECHYDHNLCGKEVSGSSSSLCNCKNSQECSTDSDHTITV
VKRYVNNQPERKRYYTCVALSSLNECTGTQEALYVLTPEANELKSVKVLCRCPSTKVYLLARNPQRYVCTTAAPL
STRTT
>ML.UMA83966.1 conotoxin precursor Tpra06 [Conus judaeus] 35
MLSMLAWTLMTAMVVMNAHGQFCPTMTDSYPGENECHYDNNLCGKEVSGSCTSLCYCKSGQMCSTDSGH
TITLVPHYTNNRPKKTRYYTCVALSSLNECSGIQEALKVLSPEAEEGESSVEVLCRCPSPKKYRFVFIRPDINGDN
PRYVCANAPRRHG
>ML.UMA82654.1 conotoxin precursor Tpra06 [Conus ebraeus] 36
MLSMLAWTLMTAMVVMNAHGQVCPTMTDSYTGENECFYDNNLCGKEVSGSCSSLCNCKNSQECSTDSDHT
ITVVKRYVNNQPERKRYYTCVALSSLNECAVREKALFVLAPEAADPISVKVLCRCPFPKVYLQTGRSRPHICSYNP
RRRG
>ML.UMA83967.1 conotoxin precursor Tpra06 [Conus judaeus] 37
MLSMLAWTLMTAMVVMNAHGQFCPTMTDSYPGENECHYDNNLCGKEVSGSCTSLCYCKSGQMCFRDNDH
TITAVPRIINDRPVEITYYTCVALSSLNECSGIQEALKVLSPEAEEGESSVEVLCRCPSPKKYRFVFIRPDINGDNPR
YVCANAPRRHG
>ML.DAZ86973.1 TPA_inf: conotoxin precursor Tpra06 [Conus judaeus] 39
MLSMLAWTLMTAMVVMNAHGQFCPTMTDSRPGENKCLHDNYLCGKEDDSGSCTSLCHCKSGQMCFRDN
DHTITAVPRIINDRPVEITYYTCVALSSLNECSGIQEALKVLSPEAEEGESSVEVLCRCPSPKKYRFVFIRPDINGD
NPRYVCANAPRRHG
>ML.AXL95365.1 conotoxin-like precursor unassigned superfamily 13 [Conus ermineus] 42
MLSMLAWTLMTAMVVMNAKSQFCPVLTDSYRFEHRCAYDNSLCGKEVSGSCTSTCYCRSGRMCLRNSDHTIT
VVKRYINNNPVKESYHTCVALSGLPRCSGNRVALYNLVPESSAFKSVEVRCRCPSPNVYLNTGLNQVYTCAPAP
QLNVVG
>ML.ATF27771.1 conotoxin [Conus praecellens] 45
MLLMFAWTLMTAMMVMNASSKDCPLLDDSNPLKRRCLWNNAICGKSVSGKCTSLCNCRNGQKCSMNSTHT
ITVVPYYINGVPVKKRYYTCMDVAELGQCSSTQEALYSLIYEETELKNAKVYCECRSPKVYLRLFTPKRYICRRAEP
RTG
>ML.QFQ61139.1 superfamily Cerm-13 [Conus magus] 46
MLSMLAWTLMTAMVVMNAKSQYCPTMRDSYPKQHECLYKNALCGKEDSGGRCSSICNCKNGQMCSTDRD
HNVTIVIHENTLPKKKRYYTCLSLWRLNECSETEKALHRLTHRTKELKSVKVLCRCPYPKAYVTLESRYRYTCALVQ
VLGD
>ML.UBT01704.1 conotoxin precursor superfamily Tpra 06, partial [Conus ammiralis] 47

MLSMLAWTLMTAMVVMNAKSQTTCPTSTNIDSCVNDNNACGKDVSGNCSSLCNCGNGQTCSTDSSHTITLV
PYYTEDGPYEKKYYTCGDPSELDECYDIDKALEVNESDDPNSVEVLCHCPSDQIYLWIHLQYYICTPPPQPDI
>ML.UMA83598.1 conotoxin precursor Cver06 [Conus judaeus] 55
MLSVFTVVWVLTTVMMMTDVTFQSTCDTDNLELCSEATHMCGKRISWDGCNGLCKCRTQQACTTDADHTVQ
VIPAPFQSNKTYYTCRSLSTMGACQSNNEAMSGNSEDTYKILCKCDETYQPHSLNNRTFVCR
>ML.UMA82347.1 conotoxin precursor Cver06 [Conus ebraeus] 56
MLSVFTVVWVLTTVMMMTDVTFQSTCNTDNLELCSEATRLCGKGTSWDQCIGLCKCRNEQACTTDADHTVQV
IPAPFQSNKTYYTCRSLSTMDACQSNERAMSGNSENKYKILCKCDKTYQPRSLNDRKFVCQ
>ML.UMA82346.1 conotoxin precursor Cver06 [Conus ebraeus] 57
MLSVFTVVWVLTTVMMMTDVTFQSTCNTDNLELCSEATRLCGKGTSWDQCIGLCKCRNEQACTTDADHTVQV
IPAPFQSNKTYYTCRSLSTMGACQSNNEAMSGNSEDTYKILCKCDETYQPHSLNNRTFVCR
>ML.UMA82626.1 conotoxin precursor Cver06 [Conus ebraeus] 58
MLSVFTVVWVLTTVMMMTDVTFQSTCNTDNLELCSEATRLCGKGTSWDQCIGLCKCRNEQACTTDADHTVQV
IRAPFQNRETYYTCRSLSTMDACQSNERAMSGNSENKYKILCKCDKTYQPRSLNDRKFVCQ
>ML.UMA82627.1 conotoxin precursor Cver06 [Conus ebraeus] 59
MLSVFTVVWVLTTVMMMTDVTFQSTCNTDNLELCSEATRLCGNRTSWDRCNGLCKCPKDQACITDDDHTVQ
VIRAPFQNRETYYTCRSLSTMDACQSNERAMSGNSENKYKILCKCDKTYQPRSLNDRKFVCQ
>ML.GCVH01000124.1 TSA: Conus lenavati Cln_SF6_1 transcribed RNA sequence 61
MLSMFAWTLMTATVVVIAERQYCPIAGQTCTFGSDLCGKEESGSCSPRCNCKNERMCSRDSDHTITVVRVFRR
RRPVEERYYTCVALSGLEECSNQKALTDLVPETRELNSVEVHCKCSSPKVYGYHMYLKGYFCGTYERS
>ML.GCVH01000118.1 TSA: Conus lenavati Cln_SF2_1 transcribed RNA sequence 62
MLFVFTVVWILTMVMIITDVTFQSTCNTDNKPSCSEDTRLCGKNNSWGNCVALCKCPNQQACTTDTDHKVQV
KRGPFQLTETYYTCKNVSTMSDCQSNAKAMSGTSESTYKIMCKCDDTYKPSAPTNWKFICG
>ML.SRR1803937.C.tribblei.TRINITY_DN1174_c0_g1_i1_1 supFam_MMLFM tpm_313.02 91
IMLFVFTVVWILTMVMIITDVTFQSTCNTDNKPSCSEDTRLCGKNNSWGNCVALCKCPNQQACTTDTDHKVQV
KRGPFQLTETYYTCKNVSTMSDCQSNAKAMSGTSESTYKIMCKCDDTYKPSAPTNWKFICG
