## Supporting file 2 for "Structural similarities reveal an expansive conotoxin family with a two-finger toxin fold"

**Supporting File 2.** All sequences used for the analyses in Fig. 2.

>SRR15402271.C.textile.TRINITY_DN602_c0_g1_i1_1 supFam_MLSML
MLSMLAWTLMTAMVVMNAKSHTTCPTSTEIDSCSNDNNACGKDVSGSCSSLCNCGNGQTCFTDSNHTITLVP
YYTEDGPFEKKYYTCGDPSELDECYDIDKALEVNESDDPNSVEVLCHCPSDKIYLWIHRGYYICITPPQP
>coronatus.TRINITY_DN558_c0_g1_i1_1 supFam_MLSML
MLSMLAWTLMTAMVVMNARGQFCPTVTDECFYDNNLCGKKVSGSCTSLCNCKSGERCSRDSDHTITLVPSYT
NGRPDERRYYTCVALSSLNECSSTEEALHDLVPEAGERNNVKVLCRCPSPKVYLFVRNPQRYICALAP
>ebraeus.TRINITY_DN659_c0_g1_i1_1 supFam_MLSML
MLSMLAWTLMTAMVVMNAHGQVCPTMTDSYTGENECFYDNNLCGKEVSGSCSSRCYCKNGGRCSTDSDHT
ITVVQSYINGYPEKRYYTCVALSSLNECAVREKALFVLAPEAADPISVKVLCRCPFPKVYLQTGRSRPHICSYNPR
RRG
>ebraeus.TRINITY_DN659_c0_g1_i2_1 supFam_MLSML
MLSMLAWTLMTAMVVMNAHGQVCPTMTDSYARENECHYDHNLCGKEVSGSCSSLCNCKNSQECSTDSDH
TITVVKRYVNNQPERKRYYTCVALSSLNECSGTQEALYVLTPEANELKSVKVLCRCPSTKVYLLARNPQRYVCTTA
AQLSTRTT
>lividus.TRINITY_DN566_c0_g1_i2_1 supFam_MLSML
MLSLLAWTLMTAMVIMNARSQFCPTMTDSYPSENECHYDNALCGKEVSGSCSPICYCKNGQMCSMDSDHTI
TVVPYYVNYYPHEIRYYTCVALSSLHQCARTEKAIHNLVPEVAEFQSVKVLCKCPSPKVYLKVNDRRRYACVMAP
PLNG
>rattus.TRINITY_DN825_c0_g1_i1_1 supFam_MLSML
MLPIFACALMTLTLVASTEFCPTTDNMCLYDDDLCGKRDSSGSCTSRCNCKNRQLCARDSEHTITVVRQFINAR
MQRESYYTCTALSALSPCSDGQVALTDLVPDTLPLSSVEVLCACRPPRIYLRTVKPTRYICSRVI
>rattus.TRINITY_DN825_c0_g1_i3_1 supFam_MLSML
MLPIFACALMTLTLVVNASNRFCPITDNMCHYDADLCGKRESSGSCTSRCHCKNMRLCVRDSEHTITVVKQIIN
HFMKRESYYTCTALSALSPCSDGQVALTDLVHETVPLNSVEVLCACRPPKVYLRTVGQYICSRGI
>rattus.TRINITY_DN654_c0_g1_i1_1 supFam_MLSML
MTLSVIKLTLITMITAVAVTVCSSSDACPNTASEACGLNRHACGKKTPWGTCEHLCRCPNNQPCLRDSKHTFQ
WKRQFLRDKETYYTCNDMSTLPACGNNAGLLHTSGEVTILCECLPPSYYAMESSKYICFRSG
>virgo.TRINITY_DN10252_c0_g1_i1_1 supFam_MLSML
MLSMLAWTLMTAMVVMNAKSQYCPTMEEDSYPGEHRCFYANDLCGKDVSGSCSSICYCNDGQMCSTDSDH
TITIVPHYVNYFPVRQRYYTCVALSSLHECSRGERAIYDLVPETPELTSVEVLCKCSSPKVYLKVGTARYVCARPPR
LTPSARHLHDDPRRTS
>SRR11807493.C.infinitus.VG.TRINITY_DN59_c0_g2_i3_1 supFam_MLSML
MLSMLAWTLMTAMVVMNAKSQFCPAMKNGYRDEHKCLRDNSLCGKEVSGSCSSICYCRNLHMCSTDSDHTI
TVVPHYVNYYPVKKRYYTCVALSGLDECSNGQISLRDIIPGTAELKYAKVLCNCRSPKVYKRLINPERYICALAPRL
PH
>SRR11807495.C.miruchae.VG.TRINITY_DN82_c0_g1_i5_1 supFam_MLSML
RNQCFYDNNLCGKKVSGSCSSICYCKNGKMCYTESDHTITVVPYYVYHHPVKMRYYTCVALSGLEECSNDETA
LYNLVPIAAELKTAKVLCKCPSPKVYLSTGRNQRYICARAPPLKG
>SRR11807496.C.cuneolus.VG.TRINITY_DN3685_c0_g1_i1_1 supFam_MLSML
MLSMLAWTLMTAMVVMNAKSQFCPTMEDRYPDEHKCFYDNNLCGKKVSGSCSSICYCKNGKMCYTESDHTI
TVVPYYVYHHPVKMRYYTCVALSGLEECSNDETALYNLVPNAGELKTAKVLCKCPSPKVYLSTGRNERYICARAP
PLNG
>SRR11807498.C.verdensis.VG.TRINITY_DN2612_c0_g2_i1_1 supFam_MLSML
MLSMLAWTLMTAMVVMNAKSQFCPTMEDSYPDEHKCFYDNNLCGKKVSGSCSSICYCKNGKMCSTDDDHTI
TVVPYYENRHLVKKRYHTCVALSGLEECSNGETALYNLVPDAAELKTAKVLCKCPSPKAYLNTGKNQRYTCGRAS
P
>SRR13740844.C.ventricosus.TRINITY_DN1909_c0_g2_i1_1 supFam_MLSML

MLSMLAWTLMTAMVVMNAKSQFCPTMEDSYDNEHRCLYDNGLCGKEVSGSCSSICYCRNLQMCSTDSDHTI
TVVPYYVNYYPVKKRYYTCVALSGLNECSGTQNALYDLIAETTELIHAKVLCNCRSPKVYKRIINPVRYICAHARPL
NG
>SRR14407584.C.abbreviatus.TRINITY_DN11_c0_g1_i1_1 supFam_MLSML
MLPIFACALMTLTLVVNASDQFCPTTDNMNQCFYDDALCGKRDSSQGCTSRCYCKNMQLCVRDSEHTITIVKR
IISGHMIKESYYTCTALSALSECDRMQKALEDLVPWTLELDSAKVLCACRPPKIYLRTINPTRYSCFF
>SRR15402271.C.textile.TRINITY_DN602_c0_g2_i4_1 supFam_MLSML
MLSMLAWTLMTAMVVMNAKSHTICPTSTNLCVNDNDACGKDVSGSCSSICNCKNGQPCSTDSSHTIILVPRYT
EDGPYIKNYYTCVDPSELVGCAGAQKSVNVSVSEAQDPNSAQVLCYCPPSKINIWALNFQHYICVTPPQP
>SRR1544627.C.miliaris.TRINITY_DN3169_c0_g1_i1_1 supFam_MLSML
MLSMFAWTLMTAMVVMNAHGQFCPTMTDINYPGENRCFYDNNLCGKEDSGSCTSLCYCKSGQMCSRDTD
HTITLVPRITNSGPDERRYYTCVALSSLNECSGTEKALHILSPEAEEPISVEVLCRCPSKVYRFILYPRPQRYICAPA
S
>SRR17653514.C.judaeus.TRINITY_DN505_c0_g1_i2_1 supFam_MLSML
MLSMLAWTLMTAMVVMNAHGQFCPTMTDSRPGENKCLHDNYLCGKEDDSGSCTSLCYCKSGQMCFRDND
HTITAVPRIINDRPVEITYYTCVALSSLNECSGIQEALKVLSPEAEEGESSVEVLCRCPSPKKYRFVFIRPDINGDNP
RYVCANAPRRHG
>SRR17653514.C.judaeus.TRINITY_DN740_c0_g1_i3_1 supFam_MLSML
MLSVFTVVWVLTTVMMMTDVTFQSTCDTDNLELCSEATHMCGKRISWDGCNGLCKCRTLQACTTDADHTVQ
VIPAPFQSNKTYYTCRSLSTMGACQSNNEAMSGNSEDTYKILCKCDETYQPHSLNNRTFVCR
>SRR2124878.C.betulinus.TRINITY_DN401_c0_g1_i2_1 supFam_MLSML
MLSVFTVVWVLTTAMMMTDVTLQSTCDTDDLGLCSEDTRLCGKRTVWNRCNGLCKCPNQQACTTDTDHTVR
VRSAPFQLIQTYYTCRDVSTMDACQSNERAMDGHNEETYKILCKCDNIYQPNAPQNWYFICS
>consors.TRINITY_DN5487_c0_g2_i1_1 supFam_MLSML
MLSMLAWTLMTAMVVMNAKSQYCPTMRDSYPKQHKCLYENALCGEEDSGGRCSSICNCINGQMCSTDSDH
TITIAKYENNRLSRKRHHTCTALWGLKECNGTKKALYVLAPESEKLKSVEVLCRCPPSGVYIKIKNPEKYICVRIRG
AQG
>MLSMLGm1_gloriamaris.TRINITY_DN3174_c0_g1_i2_1 supFam_MLSML
MLSMLAWTLMTAMVVMNAKSDTTCPTSTNIVSCVNDNDACGKDVSGSCSSICNCKNGQTCSTDSNHTIILVPY
YTEYGPYIKNYYTCVDPSDLDGCSIAQKSVNVSVSEAEDPNSVEVLCYCPPSKINIWAVNFQYYLCTTPPQPDS
>MLSMLGm2_gloriamaris.TRINITY_DN3174_c0_g1_i3_1 supFam_MLSML
MLSMLAWTLMTAMVVMNAKSQTTCPTSTNIDSCSNNNNTCGKDVSGSCSSLCNCQNGQTCFTDSNHTITLV
PYYTEDGPFEKRYYTCRDPSELDECYDINKALEVSESDDPNSVEVLCHCPSDKIYLWVHLQYYVCVLPPQPDS
>DAZ86972.1_TPA_inf:_conotoxin_precursor_Tpra06_[Conus_judaeus] supFam_MLSML
MLSMLAWTLMTAMVVMNAHGQFCPTMTDSYPGENECHYDNNLCGKEVSGSCTSLCHCKSGQMCSTDSGH
TITLVPHYTNNRPKKTRYYTCVALSSLNECSGSETALYDLVPEAEERNNVKVLCRCPSPKVYRKVLNPKRYICANA
PPRRG
>DAZ86971.1_TPA_inf:_conotoxin_precursor_Tpra06_[Conus_judaeus] supFam_MLSML
MLSMLAWTLMTAMVVMNAHGQFCPTMTDSYPGENECHYDNNLCGKEVSGSCTSLCHCKSGQMCSRDND
HTIPVVPRNINDRPDESRYYTCVALSSLNECSGSETALYDLVPEAEERNNVKVLCRCPSPKVYRKVLNPKRYICA
NAPPRRG
>UMA83966.1_conotoxin_precursor_Tpra06_[Conus_judaeus] supFam_MLSML
MLSMLAWTLMTAMVVMNAHGQFCPTMTDSYPGENECHYDNNLCGKEVSGSCTSLCYCKSGQMCSTDSGH
TITLVPHYTNNRPKKTRYYTCVALSSLNECSGIQEALKVLSPEAEEGESSVEVLCRCPSPKKYRFVFIRPDINGDN
PRYVCANAPRRHG
>UMA82654.1_conotoxin_precursor_Tpra06_[Conus_ebraeus] supFam_MLSML
MLSMLAWTLMTAMVVMNAHGQVCPTMTDSYTGENECFYDNNLCGKEVSGSCSSLCNCKNSQECSTDSDHT
ITVVKRYVNNQPERKRYYTCVALSSLNECAVREKALFVLAPEAADPISVKVLCRCPFPKVYLQTGRSRPHICSYNP
RRRG
>UMA83969.1_conotoxin_precursor_Tpra06_[Conus_judaeus] supFam_MLSML
MLSMLAWTLMTAMVVMNAHGQFCPTMTDSYPGENECHYDNNLCGKEVSGSCTSLCYCKSGQMCSRDNDH
TIPVVPRNINDRPDESRYYTCVALSSLNECSGIQEALKVLSPEAEEGESSVEVLCRCPSPKKVPFRLHPS

>QFQ61141.1_superfamily_Cerm13,_partial_[Conus_magus] supFam_MLSML
MLSMLAWTLMTAMVVMNAKSQYCPTMRERYRGQRTCLYENNLCGKKGSGGSCSSICNCKNGQMCSTDSDH
TVTIVKYENGRPYRKRYHTCLSLSGLNECSRTEEALYRLSHEIEELKSVKVLCRCPSPKVYVKL
>AXL95365.1_conotoxinlike_precursor_unassigned_superfamily_13_[Conus_ermineus] supFam_MLSML
MLSMLAWTLMTAMVVMNAKSQFCPVLTDSYRFEHRCAYDNSLCGKEVSGSCTSTCYCRSGRMCLRNSDHTIT
VVKRYINNNPVKESYHTCVALSGLPRCSGNRVALYNLVPESSAFKSVEVRCRCPSPNVYLNTGLNQVYTCAPAP
QLNVVG
>UMA82974.1_conotoxin_precursor_Tpra06_[Conus_ebraeus] supFam_MLSML
MLSMLAWTLMTAMVVMNAHGQVCPTMTDSYTGENEEVSGSCSSRCYCKNGGRCSTDSDHTITVVQSYINGY
PEKRYYTCVALSSLNECAVREKALFVLAPEAADPISVKVLCRCPFPKVYLQTGRSRPHICSYNPRRRG
>UMA83965.1_conotoxin_precursor_Tpra06,_partial_[Conus_judaeus] supFam_MLSML
CPTMTDSRPGENKCLHDNYLCGKEDVSGSCTSLCYCKSGQMCFRDNDHTITAVPRIINDRPVEITYYTCVALSS
LNECSGSETALYDLVPEAEERNNVKVLCRCPSPKVYRKVLNPKRYICAIASPRRG
>ATF27771.1_conotoxin_[Conus_praecellens] supFam_MLSML
MLLMFAWTLMTAMMVMNASSKDCPLLDDSNPLKRRCLWNNAICGKSVSGKCTSLCNCRNGQKCSMNSTHT
ITVVPYYINGVPVKKRYYTCMDVAELGQCSSTQEALYSLIYEETELKNAKVYCECRSPKVYLRLFTPKRYICRRAEP
RTG
>QFQ61139.1_superfamily_Cerm13_[Conus_magus] supFam_MLSML
MLSMLAWTLMTAMVVMNAKSQYCPTMRDSYPKQHECLYKNALCGKEDSGGRCSSICNCKNGQMCSTDRD
HNVTIVIHENTLPKKKRYYTCLSLWRLNECSETEKALHRLTHRTKELKSVKVLCRCPYPKAYVTLESRYRYTCALVQ
VLGD
>UBT01704.1_conotoxin_precursor_superfamily_Tpra_06,_partial_[Conus_ammiralis] supFam_MLSML
MLSMLAWTLMTAMVVMNAKSQTTCPTSTNIDSCVNDNNACGKDVSGNCSSLCNCGNGQTCSTDSSHTITLV
PYYTEDGPYEKKYYTCGDPSELDECYDIDKALEVNESDDPNSVEVLCHCPSDQIYLWIHLQYYICTPPPQPDI
>UMA83317.1_conotoxin_precursor_Tpra06,_partial_[Conus_judaeus] supFam_MLSML
MLSMLAWTLMTAMVVMNAHGQFCPTMTDSRPGENKCLHDNYLCGKEDDSGSCTSLCYCKSGQMCSRDSG
HTITLVPHYTNKRPKKTRYYTCVA
>UMA82347.1_conotoxin_precursor_Cver06_[Conus_ebraeus] supFam_MLSML
MLSVFTVVWVLTTVMMMTDVTFQSTCNTDNLELCSEATRLCGKGTSWDQCIGLCKCRNEQACTTDADHTVQV
IPAPFQSNKTYYTCRSLSTMDACQSNERAMSGNSENKYKILCKCDKTYQPRSLNDRKFVCQ
>UMA82346.1_conotoxin_precursor_Cver06_[Conus_ebraeus] supFam_MLSML
MLSVFTVVWVLTTVMMMTDVTFQSTCNTDNLELCSEATRLCGKGTSWDQCIGLCKCRNEQACTTDADHTVQV
IPAPFQSNKTYYTCRSLSTMGACQSNNEAMSGNSEDTYKILCKCDETYQPHSLNNRTFVCR
>UMA82627.1_conotoxin_precursor_Cver06_[Conus_ebraeus] supFam_MLSML
MLSVFTVVWVLTTVMMMTDVTFQSTCNTDNLELCSEATRLCGNRTSWDRCNGLCKCPKDQACITDDDHTVQ
VIRAPFQNRETYYTCRSLSTMDACQSNERAMSGNSENKYKILCKCDKTYQPRSLNDRKFVCQ
>GCVH01000124.1_TSA:_Conus_lenavati_Cln_SF6_1_transcribed_RNA_sequence supFam_MLSML
MLSMFAWTLMTATVVVIAERQYCPIAGQTCTFGSDLCGKEESGSCSPRCNCKNERMCSRDSDHTITVVRVFRR
RRPVEERYYTCVALSGLEECSNQKALTDLVPETRELNSVEVHCKCSSPKVYGYHMYLKGYFCGTYERS

>coronatus.TRINITY_DN675_c1_g1_i1_1 supFam_MMLFM
MMLFMFAAIIFTMVSTTVTVDRCANEQKVCRTVDDSGIGEFCNCDWVPGGCIMDDDHKFVFAESTLYTCQPVL
DFSVCQGTQTVFVAKFPQFNCRCDPHEYKIENRKIVCA
>ebraeus.TRINITY_DN480_c0_g2_i1_1 supFam_MMLFM
MMTLSVIFFLLVTGVIVTKAQANSCGDTPTTSCSEPGKICAVLFPGGQCAQLCTCGNENNACSIDDEHRISTNPT
STPAWFLCGQIEAFPICQAGENPYNRSEELFHCRCGNGYSFITNKSVFVCA
>imperialis_MP.TRINITY_DN480_c0_g1_i101_1 supFam_MMLFM

FMFAAIISIAVSARLEQCLDNRYACARIDENGQRKQLCKCINGACIKKKDYKVTVHLGKDRRTAIMCIPIDDFARC
TGHPHEYAMDSDLYKLKCKCEDKYYQEGFHYDVKCGN
>imperialis_MP.TRINITY_DN480_c0_g1_i102_1 supFam_MMLFM
IAVSARLEQCLDNRYACARIDENGQRKRLCKCINGACIKKKDYKVTVHLGKDRRTAIMCIPIDEFPRCTGRPHEY
AMDSDLYKLNCKCEDKYYQEGFLYDVKCGN
>lividus.TRINITY_DN687_c0_g1_i1_1 supFam_MMLFM
MMLFVFAAIIFTMATATEDQWPCANNKAVCAWKEAPINNCDCSGTACQRDSAHSVNVAGSIFYTCQQISDFDE
CVGSGAVMNAAFTQLLCRCSSGVYIVNEDEAIVCG
>rattus.TRINITY_DN1956_c0_g1_i1_1 supFam_MMLFM
MMLFVFAVIIFTMGSTTVSVESAPETEKTRGPKVLSLTRGIRKTRESDNYQPDKCVTETAACSVIQGGVRTPVCKC
QETGTCPTEAQYSFPISQNGHHFTVYTCKRENAFHVCSTTQAQDVFTDGNINCRCSSNNIMIMRNGHVVCTPA
MS
>rattus.TRINITY_DN376_c0_g1_i1_1 supFam_MMLFM
MMLFIFAAIIFTMAYTTVTAQDQCKNDMEACAMVLPPPNQLRDVMCHCIGTGRCNFHPSHKIDVEGTLFYTCQ
QIVHFDPCDGSKPAMDKSLMQLDCRCPNNEYKFDFAGNVVCSPTRP
>SRR11807493.C.infinitus.VG.TRINITY_DN47_c0_g1_i2_1 supFam_MMLFM
MMLFIFAAIIFTMASTTVIEDMCENGGQSVCAWKGMGEMCICPDPYSCVQSDAYKAEVTLVNRTETFYTCQQIS
DFSLCAASDPAMQGQLQQLNCRCSNGIYVLGVGAIVCG
>SRR11807493.C.infinitus.VG.TRINITY_DN47_c0_g2_i1_1 supFam_MMLFM
MMLFMFAAIIFTMASRTVTDNTCAHNKKACAMVLGDYVDIHCDCASTGNCIIHPAHKIAVNDGTLFYTCQLISDF
NICKGSEPVMDAGLTQLNCRCPNGAYKYEGSAVVCS
>SRR11807496.C.cuneolus.VG.TRINITY_DN6_c0_g3_i1_1 supFam_MMLFM
MMLFMFAAIIFTMASRTVTDNTCAHNKKACAMVLGDLVDIHCDCASTGKCIRQPGHDITANDGTRFYTCQLISD
FNICEGSEPVMDAGLTQLNCRCPNGAYKFEGKAVVCS
>SRR11807498.C.verdensis.VG.TRINITY_DN268_c0_g3_i2_1 supFam_MMLFM
LLPTLSSATHRIVHVAYRICWRLRKARLTMMLFIFAAIIFTMALTTVTAEMCENGGRSVCAWKGLGEMCICPDPYS
CVQSDAYKAVVTLENRTETFYTCQQISRFSLCAASDPALQGQLQQLKCRCSNDIYVLGVGAIVCG
>SRR11807498.C.verdensis.VG.TRINITY_DN268_c0_g3_i4_1 supFam_MMLFM
MGIFVDGAPLLLHPCFSDGRFLSTLSSAIHRIVYVAYRICWRLRKARLTMMLFIFAAIIFTMASTTVTAERCASRKSV
CAWEVLGKTCSCPDPYSCSQRYHYKAELTVRVGKLSFYTCQMIAAFRRCTPRSRPAMELDAKLKQLNCRCLSGI
YKRVNDTVVCG
>SRR11807507.C.grahami.VG.TRINITY_DN23_c0_g2_i4_1 supFam_MMLFM
MMLFMFAAIIFTMASTTVTAISCELETDLCAIEIGSDLTEICTCTDGGCTRDDSHTLSVSDISLVTCQLISDFDDCTA
GQALVDDNFTQVYCKCPGDNYRIKGTQGIGCD
>SRR13740844.C.ventricosus.TRINITY_DN405_c0_g1_i6_1 supFam_MMLFM
MMLFMFAVIIFTMASTTVSDENTCEHNKKACAMVLGDYLDTHCDCASTGNCIMNPAHKIAVNDGTLFYTCQKIS
DFNICEGSQPVMDAGLTQLNCKCPNGAYKNEGNGVVCG
>SRR13740844.C.ventricosus.TRINITY_DN405_c0_g2_i1_1 supFam_MMLFM
MMLFIFAAIIFTMASTTVTAETCASGQSACAWASVGEVCSCPGSNSCTRDDAHKVKVTLKHGIYSIYTCQQISAF
RDCAGSEAAMDAQLSQLNCKCLSGTYTIVDDAVVCG
>SRR14407587.C.aristophanes.TRINITY_DN251_c0_g1_i9_1 supFam_MMLFM
MMLFMFAAIIFTMASTTVSLDRCANPTKVCRAYVDDEFHENCDCEGIPSGCIMDEDHKFVFTGTEHSGVLYTCQ
PVLDFHICDGIETVYVDIKPQLNCRCQSNVYKVGDHVFLCG
>SRR14407587.C.aristophanes.TRINITY_DN251_c0_g1_i13_1 supFam_MMLFM
MMLFMFAAVIFTMVSTTVTVDTCANEKQVCRFVADGGTYEMCNCDGVPGGCIMDDXHKFVYYEASLYTCQPV
LDFSVCQGTERVFESANQQFNCRCNPHEYKKENRKMVCA
>SRR14921183.C.ammiralis.TRINITY_DN9_c0_g1_i301_1 supFam_MMLFM
MMLFMFAAIIFTMASSTVNAHICEQNEDLCATLIDGVTTEICDCPNGACPRGNDYKITVGSVSLYTCQQISAFDD
CTRGQHLVDDDFTQIYCKCSNGMYTVGGSQGIVCV
>SRR1544627.C.miliaris.TRINITY_DN2606_c0_g1_i1_1 supFam_MMLFM
MMLFMFAAIIFTMASTTVTEDTCDNNQKACAMMLPGVRDIHCNCASTGNCIINPAHKIFVNPHTSFYSCQQISA
FDICDGSQPVMDASLTQINCRCANNEYKFEGTDVVCG

>SRR1544627.C.miliaris.TRINITY_DN2622_c0_g7_i1_1 supFam_MMLFM
GISGGCIMDDDHKFFFSRSTLYTCQPVLDFHICDGTETVFEAYHPQFNCRCVPHEYKVENTKIVCA
>SRR1544627.C.miliaris.TRINITY_DN2622_c0_g8_i2_1 supFam_MMLFM
MMLFMFAAIIFTMVSTTVTVDTCANEQKVCRTVDDNGIGEFCNCDGVPGGCIMDDDHKFVFLRSTLYTCQPVS
DFHICDGTETVFVAYHPQFNCRCDPHEYKVEKTKILCA
>SRR1544627.C.miliaris.TRINITY_DN2622_c0_g8_i3_1 supFam_MMLFM
MMLFMFAAIIFTVVSTTVTVNTCANEKKVCRAVVSDRAYKFCDCDGIPGGCIMDDDHKFVYSTVTLYTCQPVLDF
SVCQGTETLYEPVDPQFNCRCNPHEYKEENTIIVCA
>SRR1544627.C.miliaris.TRINITY_DN2622_c0_g8_i4_1 supFam_MMLFM
MMLFMFAAIIFTMVSTTVTLDRCANEKKVCRSVVEGRISEICDCDGIPGGCIMDDDHKFFFSKSTLYTCQPVLSF
HVCQGTQIVFTARNPQFNCRCVPHKYKIENKKIVCA
>SRR17653514.C.judaeus.TRINITY_DN187_c0_g1_i4_1 supFam_MMLFM
MMLFMFAAIIFTMASATVTLDTCASAKGVCAWAGQGESCSCSATHSCTQDDAHKVEVNMEEGIQALYTCQQIS
DFDVCTGTGAVMPADWSQLNCRCSSNKYTLENGAVVCG
>SRR17653514.C.judaeus.TRINITY_DN187_c0_g1_i5_1 supFam_MMLFM
MMLFMFAAIIFTMASATVTLDTCASAKGVCAWAGQGESCSCSAPHTCTQDDAHKVEVNMEGGVQALYTCQQI
SDFDVCTATGAVMPADFSQLYCRCSNNKYTIENDDVVCG
>SRR1803937.C.tribblei.TRINITY_DN1174_c0_g1_i1_1 supFam_MLSML
IMLFVFTVVWILTMVMIITDVTFQSTCNTDNKPSCSEDTRLCGKNNSWGNCVALCKCPNQQACTTDTDHKVQV
KRGPFQLTETYYTCKNVSTMSDCQSNAKAMSGTSESTYKIMCKCDDTYKPSAPTNWKFICG
>SRR6378469.C.litteratus.TRINITY_DN341_c0_g2_i2_1 supFam_MMLFM
MMLFMFAAIIFTMATATVSATQCNSVQTLCSWEGMGELCNCSATGSCPQNDEHKIVVGTQDIYVCQQISDFEV
CTGSVAIDSSLTELNCRCSSNAYKIEGSQVVCD
>DAZ86921.1_TPA_inf:_conotoxin_precursor_Cerm08_[Conus_judaeus] supFam_MMLFM
MMLFMFAAIIFTMASTTVSLKTCANEKKVCVAFIADRMDDICNCPGIPRGCTLKDDHAFYLYTSEYYTCQPVLDF
SVCQGTERVFVPANAQFNCRCNPHEYKVENTKIVCA
>DAZ86569.1_TPA_inf:_conotoxin_precursor_Cerm08_[Conus_judaeus] supFam_MMLFM
MMLFMFAAIIFTMASTTVSLDTCANAKKVCRAVVDGRTDEICDCDGIPGGCIMDDRHKFFFSKSTLYTCPVSAFR
ICDGSETVFQAKFPQFNCRCASNVYRGVNNKAVCG
>DAZ86920.1_TPA_inf:_conotoxin_precursor_Cerm08_[Conus_judaeus] supFam_MMLFM
MMLFMFAAIIFTMASTTVSLKTCANEKKVCVAFIADRMDDICNCPGIPRGCTLKDDHAFYLYTSEYYTCQPVLDF
HICNGSEPVFQGENTQFNCRCASNVYIVENYEVVCG
>DAZ86919.1_TPA_inf:_conotoxin_precursor_Cerm08_[Conus_judaeus] supFam_MMLFM
MMLFMFAAIIFTMVSTTVSLGTCANEQKVCLAVHMGWKDELCDCPGILGGCIMEGHHEFSLYESLQFYTCQPV
SDFRICHGSETAFDPANRQFNCRCESRKYILAANADVVCG
>UMA82335.1_conotoxin_precursor_Cerm08,_partial_[Conus_ebraeus] supFam_MMLFM
TCGNEKKVCRDVVEGRMHELCDCPEIPSRGCIMDEDHKFVFAGTTLYTCQPVKDFPICNGSETTVYEPLNPQF
NCRCESHVYKVVHYEVVCG
>UMA82334.1_conotoxin_precursor_Cerm08_[Conus_ebraeus] supFam_MMLFM
MMLFMFAAIIFTMASTTVTADTCASAKSVCAWGGQGESCSCSAPHTCTQDDAHKVEVNMGEGVQALYTCKQIS
DFNVCTGTDAVMDAQFSQLNCRCSSNTYIIQNSAVVCG
>B0L0Y6.1_RecName:_Full=Putative_conotoxin;_Flags:_Precursor_[Conus_caracteristicus]
supFam_MMLFM
MKLFMFAAIIFTMASTTVRAEQCANNRKVCTWDGQGDTNCDCIGTACHEDDAHKVSVAGSAFYTCQPISAFRV
CDGSEDVMDAGFSELYCRCSGGSYTISNGEVVCD
>UMA83589.1_conotoxin_precursor_Cerm08_[Conus_judaeus] supFam_MMLFM
MMLFMFAAIIFTMASTTVTADMCASAESVCAWKGQDGAGCSCSAPHSCIQDDDHKVEVTMGEGAQVLYTCKQ
ISDFTECTGTGDVMDTDLSQLNCRCSSNKYVLDNSAVVCG
>DAZ86917.1_TPA_inf:_conotoxin_precursor_Cerm08_[Conus_judaeus] supFam_MMLFM
MMLFMFAAIIFTMASTTVTAEQCASNQNVCKWAEQEGQNCDCIGTPCITDNAHKVSIADSAFYTCQLISAFRDC
AESEVVADGEFSQLNCKCSSGTYVINNGAVVCG
>UMA82620.1_conotoxin_precursor_Cerm08_[Conus_ebraeus] supFam_MMLFM

MMLFMFAAIIFTMASTAVTLDTCVSAESVCTWEGQAEPGCNCSAPHTCTKDDDHKIEVTLAEGTQAIYTCKKISD
FTGCSETGDVMATDFSQLNCRCSSNKYAIENGTVVCG
>UMA83587.1_conotoxin_precursor_Cerm08_[Conus_judaeus] supFam_MMLFM
MMLFMFAAIIFTMASTTVTADTCASAKSVCAWGGQGESCSCSATHTCTEDDAHKVEVNTEEGVQALYTCQQIS
DFDECTGTGAVMPADLSQLNCRCSSNKYVLDNSAVVCG
>ATF27507.1_conotoxin_[Conus_andremenezi] supFam_MMLFM
MMLFMFAAIIFTMASTTVTAETCTSDKRVCAWQGQNDTCSCSNSNSCTRDDAHKVVVTMTQGGQSFFTCKKIS
DFSACDGSQSVMNSGMSQLNCRCSGDTYKIENGVVVCG
>UMA83586.1_conotoxin_precursor_Cerm08_[Conus_judaeus] supFam_MMLFM
MMLFMFAAITFTMASATVTLDTCASAKVVCAWAGQGESCSCSATHSCTQDDAHKVEVNMERGVQALYTCQQI
SDFDKCTATGAVMPADLSQLYCRCSNNKYTIENDDVVCG
>AXL95466.1_conotoxinlike_precursor_unassigned_superfamily_08_[Conus_ermineus]
supFam_MMLFM
MKLFMFTAIIFTMASTTVTETTCENDKKACGMVLGNLRDTHCDCASTGNCIFHPAHKITVDDTLFYTCRKNSDFD
ICDGSQPVMNAALTQLNCRCPGAAYTFEGSDVICG
>UMA83588.1_conotoxin_precursor_Cerm08_[Conus_judaeus] supFam_MMLFM
MFAAISFTMASATVPLDTCASAKGLCAWAGHGESCSCSAPHTCTQDDAHKVEVNTEEGVQALYTCQQISDFDE
CTGTGAVMPADLSQLNCRCSSNKYVLDNSAVVCG
>AXL95570.1_conotoxinlike_precursor_unassigned_superfamily_08_[Conus_ermineus]
supFam_MMLFM
MKLFMFTAIIFTMASTTVTARTCRNNMEVCAIQYDEMYDYCSCSNGGCIKDDPYKVAVPGMSVFTCQQISDFNI
CGQSQFVMKDPAYQINCRCRDGRYKVVHGGIKCE
>UMA83276.1_conotoxin_precursor_Cerm08_[Conus_judaeus] supFam_MMLFM
MKLFIFAAIIVAMATAEQCPNNRAVCAWQDQEGVNCDCIGTSCTKDSAHSINAAGSTLYTCQEISAFRECVGSE
AALNSDFSQMNCRCASGVYTISGTAVVCG
>ATF27506.1_conotoxin_[Conus_andremenezi] supFam_MMLFM
MKLFVFAAIIFTMVSTTSTCEDVTSVCHVTVGSQNSSVCNCAGSSSCPKDADYAFPADQSGQDATLYACQKKAD
FSDCTNGQDVLTDMKINCKCTSGKIKLSADPEVCG
>UMA82937.1_conotoxin_precursor_Cerm08,_partial_[Conus_ebraeus] supFam_MMLFM
ANSNTCDNNQKACAMIVSSVRDIHCNCASTGNCIIDPAHKIVVNSDTLFYSCQQISAFDICVGSQPVMDASLTQ
INCRCANNKYTF
>DAZ85928.1_TPA_inf:_conotoxin_precursor_Cerm08,_partial_[Conus_ebraeus] supFam_MMLFM
AETCASAKSVCATIEQGEICNCTAPHSCRQNYDHEVEIISKGGVQTLYTCQQISDFDVCTGLDNDVMDGELKEL
KCVCQSGTFVQCDFGVECF
>GCVH01000117.1_TSA:_Conus_lenavati_Cln_SF1_1_transcribed_RNA_sequence supFam_Unknown
MMVLSVSIVVVFTKESATPLVPCSKPAEFCTTVGGTCGEYYRWGDRNYCFLFCECDVGTSCPTDEAHAIVLRNV
THPLTHYTCQPISEFPVCQVNDTAIVFRGPSLLRLVHCICHGVYDDVEVGKLVCRD
>GBRA01000064.1_TSA:_Gemmula_speciosa_TSA:Gsp_64_transcribed_RNA_sequence
supFam_MMLFM
MRMSLCLLIATILVTAGKGVSTTDVESQAHGYSADRDANIGPCQDGNGICGAQIVSLPDPGPLCTCPSGGCPLD
NDHRVPITNPQSITELYACRPINRYEQCSGSGAVAVTDALTEMKCRCTGMNYMLDEVYMRVTCAL
>GCVM01000091.1_TSA:_Conus_tribblei_Ctr_SF1_1_transcribed_RNA_sequence supFam_Unknown
MKSAVFMMVLSTSIFVVFTKDSATPLVPCSKPAEFCTTVGGTCGEYYRWGDRNYCFLFCECDVGTSCPTDEAHA
IVRRNVTHPLTHYTCQPISEFPVCQVNDTAIVFLGPSLLKLVHCTCHGEYDDVGNGRVVCRD

>SRR11807492.C.raulsilvai.VG.TRINITY_DN68_c0_g2_i2_1 supFam_Unknown
MKSAVFMMALSMSIFIDFTMESATPDVPCSVPDELCTTTGGTCGLFYHFGGRNYCFRFCKCDIGMGCPTDEAH
AVVSGNVTDPFTYYTCQPISEFPVCNVNDVAIVIPEHGHPLLHEVKCICPWVYANDGSWTYICKE
>SRR11807493.C.infinitus.VG.TRINITY_DN1512_c0_g1_i2_1 supFam_Unknown

MKSAVLMMALSMSILIDFTIESATPDGPCSDPYELCTTTGGTCGEYYHWSGRNYCFRFCECDIGMGCPTDEVH
AIIRGNVTHPLTHYTCQPISEFTACKVNDVAMLNPGPGLLLVVKCLCPWVYTYDGNGSYICKD
>SRR14407584.C.abbreviatus.TRINITY_DN3078_c0_g1_i1_1 supFam_Unknown
MKSAVFMVALSVSIFIDFTMEYAVPDELCNDLENLCSEPDHTCGEYYHMDDDLHCFRFCSCDVGMRCPIDTD
HAIVLRNSTARPWAVYSCTSLNEIPECDAGDAAIVNAPIPYRQIRCICPGEYRFIEEGDGWFKCIY
>SRR14407587.C.aristophanes.TRINITY_DN85_c0_g1_i1_1 supFam_Unknown
MKSAVFMVALSMSIFTDFTMASATPDVPCIFVTGQLKGELCTTIGETCAEYAYWNGKIHCLRLCDCGVGVACPT
DEAHAIVRGTKLDPLTVYTCKPIREFPECEVGDAAIVKYGTHLGRVVKCICPWKYAFDGNRTDVCQD
>DAZ86968.1_TPA_inf:_conotoxin_precursor_Tand01_[Conus_judaeus] supFam_Unknown
MKSAVSMMALSMSIFTDFTMASATPDVLCITPVGQLSELCTTIGGTCGEYYHWKGKYYCFRFCECDVGMPCAT
DEAHAIVRGTETHPLTHYTCQPISEFPVCEVDQVAIVNNGSGLLRVVKCICPWVYAYDSNSSHICKD
>DAZ86970.1_TPA_inf:_conotoxin_precursor_Tand01_[Conus_judaeus] supFam_Unknown
MKSAVSMMALSMSIFTDFTMASATPDVLCITPVGQLSELCTTIGGTCGEYYYREGKYYCFRFCECDVGMPCATD
EAHAIVRGNETHPSTSYTCQPISEFPVCEVDDVAIVTADPGLHRVVKCICPWVYAFDGNRSHICKD
>UMA83314.1_conotoxin_precursor_Tand01_[Conus_judaeus] supFam_Unknown
MKSAVSMMALSMSIFTDFTMASATPDVLCITPVGQLSELCTTIGGTCAEYYHWKGKNYCFRFCECDVGVACPTD
EAHAIVRGTVKRPLTHYTCEPITEFPECEVGDVAIVNSGPGLRRVVKCICPWTYIYDGNSSHICKD
>UMA82381.1_conotoxin_precursor_Tand01_[Conus_ebraeus] supFam_Unknown
MKSAVSMMALSMSIFTGFTMASATPDVLCITPVGQLSELCTTIGGTCAEYYHWKGKNNCSRFCECDVGVACPT
DEAHAIVRGTVKRPLTHYTCEPITEFPKCEVDDVAIVNSGPGLRRVVKCICPWKYMYDGNSSHICKD
>ATF27517.1_conotoxin_[Conus_andremenezi] supFam_Unknown
MKSAVFMMALSMSISIVFTKESAIPPGTRCSDPKGTCPFTDGTCGAYYRSGNANYCFRVCKCKGTNDCLIDEDH
ALVRGNKTFPSTVFTCKPIREFRECKDGETALVYPQPGLIRQIRCLCSRGWRYEYAGNGKVKCLRLT
>ATF27515.1_conotoxin_[Conus_andremenezi] supFam_Unknown
MKSAVFMMALSMSISIVFTKAEESVVKDVDCSDPKGTCPFTDGTCGAYYRSGNANYCFRVCKCKGTNDCLIDE
DHALVRGNKTFPSTVFTCKPIREFRECKDGETALVYPQPGLIRQIRCLCSRGWRYEYAGNGKVKCLRLT
>ATF27766.1_conotoxin_[Conus_praecellens] supFam_Unknown
MKSAVFMMALSMSISIVFTKAEESVVKDVDCSDPKGTCPFTDGTCGAYYRSGNANYCFRVCKCKGTNDCLIDE
DHALVRGNKTFPSTVFTCKPIREFRECADGEIALVYPQPGLIRQILCLCSRGRRYDKAGNGKYKCFD
>ATF27767.1_conotoxin_[Conus_praecellens] supFam_Unknown
MKSAVFMMALSMSISIVFTKAEESVVKGVDCSVPKGTCPFTDGTCGAYYYWGGAIHCFRVCKCEGTNDCLIDE
DHALVRGNKSFPDTVFTCKPIREFTECDNGDIALVYTQPGLIGQILCLCSRGRRYEQAGSGIYKCFD
>QFQ61149.1_superfamily_Tand01_[Conus_magus] supFam_Unknown
MRSAIFMMALSMSISIGFATEWKPPTVPCSEPVKQCPATDATCGTYYYWSVDNKNFCFLVCKCGDQNTCLTDE
DHAIVLGNETHPNTIYTCKPISKIPECDKDKTTLLPLGGLLMKVPCKCLGGYQRLRDGSYKCR
>ATF27770.1_conotoxin_[Conus_praecellens] supFam_Unknown
MKSAVFMMALSMSISIVFTMESAIAPGTHCTKPDKGTCPFTNGTCGEYNRLGPFTYCDHYCTCEVTNICPTDKD
HALEVETRIFLTCQPISKIRECDTGEVSLDFRRLGIIGEVKCICSHGRKYVQVQQDGKVHYECS
>ATF27768.1_conotoxin_[Conus_praecellens] supFam_Unknown
MKSAVFMMALSMTISIVFTMESAIAPGTRCSKPEKGRCPFISGTCGQYNRLDDFIYCDHDCTCEGTNICPTDKD
HALKVETSILFTCHPISKIRKCDNGEVSLVFFQPGSIGLVRCICSGGRKNVQVEQDGKFHFVCS

>terebra.TRINITY_DN3292_c0_g1_i1_1 supFam_E
MMTRVFITMFFLLALTQGWPQLYDEDCERGPNIHLTCGRFKQCGRIEKRNGQLKCRVLCTCPNGNSCLNGEVI
DWDDRGLKFYTCPRD
>terebra.TRINITY_DN3292_c0_g1_i2_1 supFam_E
MMTRVFIAMFFLLALTQGWPRLYDEDCVREPLADDPCIGAYQCGRAVKSNNALRCYLKCTCSHGNRCLNGEY
VNWEERSVKIFSCP
>terebra.TRINITY_DN3292_c0_g1_i3_1 supFam_E
MKTRVFIAMFFLLALTQGWKRLYDVDCVREPLADNTCIGANQCGRAIKENGNLKCYLKCTCSNGNCLNGENV
DWDDRSVKHYSCV

>14357X3.C.furvus.TRINITY_DN8547_c0_g1_i1_1 supFam_E
MMTRVFLAMFFLLVLTEGWPRLYDDDCTRGPNMHITCFKDETCGVIRKRGGSLECNLTCKCRRNESCLHGENI
DWDNRGVKIQICPKPWF
>lividus.TRINITY_DN294_c1_g1_i1_1 supFam_E
MITRVFIAMFFLLALTEGWPQLQDSDCEKGPDYHTTTCRRTKECGRIEKINGHLTCTVWCTCQKGRSCLTDEHV
EWDRPNIELQFFSCPSKWFS
>marmoreus.TRINITY_DN1638_c0_g1_i101_1 supFam_E
MMTRVFFAMFFLVALTEGWPRLYDSDCVRGRNMHITCFKDQTCGLTVKRNGRLNCSLTCSCRRGESCLHGEYI
DWDSRGLKVHICPKPWF
>marmoreus.TRINITY_DN1638_c0_g1_i102_1 supFam_E
MMTRVFFAMFFLMALTEGWPRLYDSDCVRGRNMHITCFKDQLCGLTEKRNGRLNCSLTCSCRRGESCLHGEY
IDWDSRGIKVHICPRPWF
>rattus.TRINITY_DN672_c0_g1_i1_1 supFam_E
MTRVFITMFFLLALTEGWPRMYDKNCENGPNMHPDFTCRAKEQCGTIRKRDGQLSCKLKCKCAPTGHCLNGE
DIDWDDITVKTYTCP
>sponsalis.TRINITY_DN396_c0_g1_i1_1 supFam_E
MMIRVFLAMFFLLALTEGEWERLVDEQCTKGPNMHNTCYKDTTCGRIEKKKNKLKCHLTCKCRRGESCLRGDK
IDWDYKNRNIKIYSCPLPWF
>virgo.TRINITY_DN2728_c0_g1_i1_1 supFam_E
MMTRVFITMFFLLALTQGWPRLYDSDCVREPTADNTCIGIYQCGRAVMNNGVLTCYLKCTCSNGNGCLHGEYV
DWDDRSVKHYYCS
>SRR11807493.C.infinitus.VG.TRINITY_DN82_c0_g1_i3_1 supFam_E
MITRVFIAMFFLLALTEGWPRLTDSDCELGRNMHITCKQLDQCGVIDKKDGQLTCKLRCKCKPGKRCLRKENID
WSDITTRIYHCPWP
>SRR11807502.C.guanche.VG.TRINITY_DN472_c0_g1_i2_1 supFam_E
MMTRVFIAMFFLLALTEGWPRLNDWDCELGRNMHITCKQLEQCGFIEKKDGQLTCKLRCRCKPGKRCLRKEKI
DWSDITTRIYHCPWP
>SRR11807506.C.trochulus.VG.TRINITY_DN1934_c0_g1_i1_1 supFam_E
MTRVFIAMFFLLALTEGWPRLHDDNCENGPNMHYSYTCDTQQQCAIIRKRNGQWTCKLMCKCEPGVDCLHG
EYIDWDDRTAKIIYCP
>SRR13740844.C.ventricosus.TRINITY_DN197_c0_g3_i2_1 supFam_E
MMTRVFIAMFFLLALTEGWPRLHDWDCEVGRNMHLTCTQLEQCGLIEKNDGQLTCKLRCKCKPGKRCLREEN
IDWSRITTRIYHCPWP
>SRR13740844.C.ventricosus.TRINITY_DN197_c0_g2_i1_1 supFam_E
MMTRVFIAMFFLLALTEGWPRLYDRNCQNGLNMHYSYTCDTQRQCAIIRKRKGQLTCKLMCKCEPGVDCLRG
EYIDWDDRTAKIIYCP
>SRR14407584.C.abbreviatus.TRINITY_DN65_c0_g1_i1_1 supFam_E
MMTSVFFAMFFLLALVEGWPRLKDTDCTKQGSYDQGRQCFKNEQCGRIEKRDGKLKCYVRCKCIRGNSCLQ
DDDLDWDGIRTIKIYSCPRPWF
>SRR14407584.C.abbreviatus.TRINITY_DN96_c0_g1_i2_1 supFam_E
MMTSVFFAMFFLLALVEGWPRLKDTNCTKQGSYDQGRQYFKNEQCGRIEKRDGKLKCYVRCKCIRGNSCLQD
EDLDWDGTRNVKIYSCPRPWF
>SRR14921183.C.ammiralis.TRINITY_DN806_c0_g1_i1_1 supFam_E
MMTRVFLVMFFLLVLTEGWPRLTDSDCERGRNMHITCFKDQSCGLIVKRNGRLSCSLNCKCRRNESCLPSEEV
DWDHRDMKIVICPKPWF
>SRR1544627.C.miliaris.TRINITY_DN2006_c0_g1_i1_1 supFam_E
MRRVFIAMFFLLALTEGWPSLYDQNCENGPNMHYSYTCPSGEKCAIIRKRDEQLTCKLMCKCEPGVDCLHGE
DINWDDTSAKMIYCP
>SRR17653514.C.judaeus.TRINITY_DN1415_c0_g1_i1_1 supFam_E
MTRVFLAMFFLLALTEGWPRLYDRDCERGRNMHYTCTQLEQCGVIRKMNGQLTCELRCRCTPGKSCLHGENI
DWDNMNTKIYHCPWP
>SRR2609537.C.quercinus.TRINITY_DN77_c0_g1_i2_1 supFam_E

MMTRVFIAMSFLLAITEGWPILYDSDCEPSRNMHLTCASYEQCGRIMKNSGELTCILTCSCPNGDSCLPGDYIDWSDQTQQFYTCP
>SRR6378469.C.litteratus.TRINITY_DN279_c0_g1_i2_1 supFam_E
MVRVFIAMFFLWALTEGWPRLHDRNCQNGPNMHHTYRCRSRQRCAIIRKRNGQLTCELKCKCESVGDCLQGEVVDWDVRTVKTYTCP
>EGm1_gloriamaris.TRINITY_DN688_c0_g1_i1_1 supFam_E
MMTRVFLAMFFLLVLTEEWPELSDSDCERGPNMHITCFKDQSCGLIVKRNGRLSCTLNCKCRRNESCLPSEEVDWDHRNMKIVICPKPWF
>UMA82449.1_conotoxin_precursor_E_[Conus_ebraeus] supFam_E
MTRVFIAMFFLLALTEGWPRLYDRDCERGRNMHYRCKQLEQCGIIRKKDGKLTCELRCSCKPGKSCLHGENIDWDDMTTKIYHCPWP
>W4VS70.1_RecName:_Full=Conotoxin_Vc22.1;_AltName:_Full=E_Vc1;_Flags:_Precursor_[Conus_victoriae] supFam_E
MMTRVFLAMFFLLVLTKGWPRLYDGDCTRGPNMHITCFKDQSCGLIVKRNGRLSCTLNCKCRRNESCLPSEEVDWDNRNMKIVICPKPWF
>AMP44647.1_conotoxin_[Conus_betulinus] supFam_E
MTRVFVAMFFLLALTEGWPPLYDRNCENGPNMHYSYTCGTKRKCAIIRKRNGQSTCKLMCKCEPGVDCLHGENIDWDDRTAKIIYCP
>UMA82732.1_conotoxin_precursor_E_[Conus_ebraeus] supFam_E
MMTRVFFAMFFLMALTEGWPRLYDSDCVRGRNMHITCFKDQSCGLIVKRNGRLSCSLNCKCRRNESCLPSEEVDWDHRDMKIVICPKPWF
>AXL95534.1_conotoxin_precursor_superfamily_E,_partial_[Conus_ermineus] supFam_E
MMTRVFIAMFFLLALTEGWPRLYDRNCQNGLNMHYSYKCDTQQQCAIIRKRKGQLTCKLMCKCEPGIDCLHGEYIDWDYRTAKIVY
>A0A125S9D9.1_RecName:_Full=Conotoxin_Im22.1;_AltName:_Full=Conopeptide_im005;_Flags:_Precursor_[Conus_imperialis] supFam_E
MMMRVFIAMFFLLALVEAGWPRLYDKNCKKNILRTYCSNKICGEATKNTNGELQCTMYCRCANGCFRGQYIDWPNQQTNLLFC
>GCVH01000032.1_TSA:_Conus_lenavati_Cln_E_1_transcribed_RNA_sequence supFam_E
MMMRVFIAMLFLLALTEGWPRMYDKNCTTDLNYANGIQCFRAEQCGAIRKVNGELTCYLKCNCVRGDDCLTGEYINWDGNKNIKIFSCPKPWQ

>terebra.TRINITY_DN2465_c0_g1_i1_1 supFam_MARFL
MARFRSILLCIAMAVALAAGIYYPSSEEEIGPCSTNDISKTETYTDDDGGYVLDYSYCTCASGEVHFAADDTTTSSSTVPHKIYVCGAPTQSCTGGTLPVTDPDYDGPRRMQCTCGEYKYFVSHFGWHVRCK
>arenatus.TRINITY_DN677_c0_g1_i1_1 supFam_MARFL
MARFLSILLCIAVAVAPAAGIYYPNGNPVGPCSRNDISKTEVYSERYERYVLHYSYCSCASGEVHFQADDTTSSHNTAFYKIYECGAPTFSCSGRAMFPVTDSDQGTRRMLCRCGAYKYFVSRLGWHVRCK
>ebraeus.TRINITY_DN1775_c0_g1_i1_1 supFam_MARFL
MARLLSMLLCIAVAVVLAAGIQYSAESPMQQCSTHAVSKMEVYSNNQGKYELLDSYCSCDSGEAYFEAQDTTTSSSTVSSYKIYVCGMPTESCTGGTLPVTDSDNGPRRMQCTCGQYKYLVSASLGWHVRCK
>imperialis_MP.TRINITY_DN183_c0_g1_i1_1 supFam_MARFL
MARFLSILLCFAMATGLAAGIRYPDRVLGRCSTHDLSKMEIDTNLDGVYSPHRSFCTCGSGEVYFTAKDRRNHSNYRVYVCGMPTEFCTAENPVRDPKKGNRWLQCRCRQYKMVIYRDWLVLCE
>lividus.TRINITY_DN908_c0_g1_i1_1 supFam_MARFL
MARCLSSLLCIAMAVALAAGTYYPSVETIEPCSSHDVSKTETYSDQKEGYVVDRIYCTCDSGEVHFTADDTITSSTTVSHKIYVCVAPTHSCTGTSPVTDPDDEGPRRMQCTCEEYKYLISNWGWHVLCSTQPRNVWWSRTTA
>lividus.TRINITY_DN908_c0_g1_i3_1 supFam_MARFL
AVALAAGIYYPSETIEPCSSHDVSKTETYSDRDGVYVVDHIYCTCDSGEVHFTADDTTTSSSTVSHKIYVCVAPTYSCTGGTLPVTDPDQGTRRMQCTCEEYKYFISNWGWHVRCRSD

>rattus.TRINITY_DN88_c0_g1_i1_1 supFam_MARFL
AMFLFILLGILTATKGLYVPELCIGDPSNVPCDPCQISQTQWYSTVQEKTCVRHSCRCTGTKHVFEAVYQTSTTYT
LTACCSEQARCTGTSYAVSEPGTSRVMLCSCQNYKYIINAHGGGGDYHIRCDN
>sponsalis.TRINITY_DN828_c0_g1_i2_1 supFam_MARFL
MARFLSILLCIAVTVAPTAGIFHPPSESSVGPCGTNDISKTEIYSNREQRYKPLYTYCNCASGEVHFEAVDTVSSYQ
HEYYKIYECGAPTYSCSGRMKPVTDRDDAGPREMQCTCEHYKYFLYPHLGWMVRCK
>virgo.TRINITY_DN2656_c0_g1_i1_1 supFam_MARFL
MARFLSILLCIAMAVALAAGIYYPVPEPIESCSTNDVSRTEVYSDLYGVYKPQHTYCTCASGEVYFAAEDTITSSTTS
SYKIYVCGAATHSCTGGTLPVTDSDDEGPRRMQCTCGEYNYFISRLGWHVRCK
>SRR11807498.C.verdensis.VG.TRINITY_DN5636_c0_g1_i1_1 supFam_MARFL
MARFLSILLCIAVAVALAAGIYYPDNRPIGLCETYDISKTEVYSDRFDKWVSGSTYCTCASGEEHFAAEDTTSSSKG
PYKIYVCGAPTYSCPGAMIPVTDSGDGTRRMQCTCGQYKYFVSNRLGWHVRCK
>SRR11807500.C.galeao.VG.TRINITY_DN575_c0_g1_i1_1 supFam_MARFL
MARFLSILLCIAVAVALAAGIGYPDVSKPIGPCETNDISKTEVYSNRFGRWKSHYTYCTCASGEEHFAADDTTSSN
AVVSYRIYVCGAPASPCSGATIPVTDSDDGTRRMQCTCGQYKYFVSSRLGWHVRCK
>SRR11807507.C.grahami.VG.TRINITY_DN16_c0_g2_i1_1 supFam_MARFL
MARFLSILLCIAVVMALAAGIGYHLGDRPGVGPCNTNDISKTERYSNHFSKWEIDHAGWKSHDTYCTCASGEE
HFTAIDTTSTYTSYRIYVCGAPTSPCSGAKPVTDSNDGTRRMQCTCGQYNYFVTYRLGWHVRCK
>SRR13740844.C.ventricosus.TRINITY_DN1_c0_g1_i2_1 supFam_MARFL
MARFLSLLLCIAVAVALAAGIGYPDDLFRRIGPCKMNDISKTEFYSYRYDRWQPGYTYCICYDVREDGFPAEDTTS
NADFAFYVCGAPTSPCSGATIPVTDPDDGTRRMQCTCGQYKYFLRKHVGWHIRCK
>SRR13740844.C.ventricosus.TRINITY_DN1_c0_g2_i3_1 supFam_MARFL
MARFLSLLLCIAVVMALAAGIYYPPVDKPIGPCNTNDISKTEAYSNRFGRWESHNTYCTCASGEEHFLANDTISS
YATRSYRIYVCGAPTSPCSRAIPVTDSNEGTRRMQCTCGQYKYFVSNRLGWHVRCK
>SRR17653514.C.judaeus.TRINITY_DN52_c0_g1_i1_1 supFam_MARFL
MARLLSILLCIAVAVGLAAGIQYPTVRPIGPCSTNAISKTELYSNRYGRYVLHYSYCSCASGEVYFEAQDTTTSYTTV
LYKIYACGAPTHSCSGETLPVTDPDDRPRKMQCTCEQYKYLLYPRLGWLVRCK
>SRR17653514.C.judaeus.TRINITY_DN52_c0_g1_i2_1 supFam_MARFL
MARLLSILLCIAVAVGLAAGRQYPTVRPVGPCSTNAISKTEVYSNRQEGYVLHYSYCSCASGEVYFEAQDTTKSST
TSSTTESSYKIYVCGAPTQSCTGGTFPVTDSDDSGPRRMQCTCGQYKYFVYSWGWHVRCK
>SRR2609537.C.quercinus.TRINITY_DN470_c0_g1_i1_1 supFam_MARFL
MACCLSSLLCIAMAVALATGIYYPSDETIGPCNTNDISKTETYSGRQGKYVVDRTYCTCASGEVHFAADDTTSSFT
TVPHKIYVCITPTYSCTGGTSPVTDPDDEGPRRMQCTCEEYKYFVSNWGWHVRCK

>rattus.TRINITY_DN814_c0_g1_i1_1 supFam_unk2
MLGASFILICFLWATTFLLINSSDQWDDSDRESRMLPTCISNEVCATQTHDEFKHRCACDNGCHVYNGSIPDS
DFWLWRLYTCDPVHFCQQGQNPVNAYSDQWQMLCQCLCGLYRLHMLEEIPAFECYDAHA
>rattus.TRINITY_DN814_c0_g1_i5_1 supFam_unk2
NSSDSRGESRRELTDCRDNQVCAIRSHHTTNHLCKCEGNCVDDVVVPDSPSNLARIYTCNTVDGCEGDEDPL
TVFEGEWLKLNCQCGWFLISNVNGQYEAECC
>SRR11807492.C.raulsilvai.VG.TRINITY_DN2196_c0_g1_i2_1 supFam_unk2
MLGASFTLIYFLLATTFLLNNATGSRDESRRVLSTCVPSQVCAIRSHRRTNLLCECVDECPVDDFDIPDSPHGLA
RLYTCDTVRGCEGGENPLRVVVGERLELNCQCGLVLVSLVGGECVA
