## Supporting file 4 for "Structural similarities reveal an expansive conotoxin family with a two-finger toxin fold"

**Supporting File 4.** Sequences used for the analyses performed in Fig. S11.

The following sequence was used as input for the PSI-BLAST search identifying the protostome 2FTX proteins clustered in Fig. S11A:

>ELU02186.1 hypothetical protein CAPTEDRAFT_218812 [Capitella teleta]
MRDGRPLRRLHQTLLRLLCGGGGGGDGRGGDCGAVRRRRWRPAFAHCLLCLTVLTAECIQPMCAHPCKSDY
EVCNAVLQVGSNLIEDNPKGERPYCECSDTNACPKRWSNRTGQTLTWFHHERRDWLVQYQFCSDVIERGPQ
CTMTTEVVITVKTEQSLWRPELKHIHCHCPDHIYYLQSWRLNSTGQEWYYSYNCYRRNCKLKEPCVKHYLDRG
GSRTMGYHFLCSCPHGQHCPIGNKDQRIQSYEVDDSDDIGPFISGYCEDVPGLQSNNRKSFTYESREPIRDSI
DKS*

The following sequences (all shown devoid of their predicted signal sequences) were used as input for the AlphaFold predictions of protostome 2FTX proteins shown in Fig. S11B:

Cluster A:
>KAL3848361.1_hypothetical_protein_ACJMK2_019226_[Sinanodonta_woodiana]
YQYALPECSEDAVCSEIVAISSGSMYEQTYYPVQKCKCPSFMYCPKEPGSQTISVNKDRWYGLCRPVSEIKTCQ
KGEVAEEVLFNARDIGDRIYTTIHCTCPSQLYPVLNDVGQSALGTYTKDNDIQPKLLYQFICMEDGNLFTKRGNR
GSGRRFYFY*

Cluster B:
>XP_069936290.1_U-scoloptoxin(11)-Sm6a-like_[Cherax_quadricarinatus]
TPLVAGNVATNELVEPPPMQEEADLPLCRSGSACGYLQVNAYGVSTQQFCRCARTDGSCDMHWDPQDGHSI
TMGSDQFKFCRSPPPLHVCGRDETAYSAAFIFNKATFSMAGQQHLLHCFCPPPLAHVRADVVEDVVDGELLLA
TLHSCSRLEVCTEEAPCKEVSMGGGTPLVNPKCRCPRKTSCPTLGTTVAPHSNTYPQGSAYSVFCHPGPA*

Cluster C:
>GBM48507.1_Protein_giant-lens__Araneus_ventricosus
AMVSRFPLNFRHNHKMAFRIFYQIGNSESDLPECSEMAVCNRVDTYSTPWIERQCRCPGKKVCSMSVDPRDG
YTVVDKSRQLKLCEPVNQLPTCRYFRDVAWTYATYPDNTTKQTMHCVCPKNSVAYIFKHEIYNTPDGVAVLYFLA
CSPQSKMRCQRKEPCRLFTVKKRPDVEEVSTNTLCQCPRGHYCPEHHSQPSVISSPSSFTEEHIRTYSGYCTSE
LS*

Cluster D:
>XP_068622463.1_D_Battus_philenor
MTRLIFPVLALAAVTAVASSDQTGPRILDFNPWSWLPSSNRSPSVMDYFFPPSRTPKNSNLDPPADRSPPKTTST
VVGETLTTAYQERPTTLARKSANKSKTPLSTYHTTVSPKTKKNALKTIDINATEDIKTTQSVRGDDVTQDLFKAAVTT
STARPTKHYTIRPIVRTSTATAEETESTLNTNSDKTETVAVETVATTVEDAKGTAPMFSTPESRSTDADTASTGTESST
LVDSTIAEDLDSREGYLPLRDKPQQTHVKINDQTKKETVPLPAGSTSVLAKPTKYHYYPHNQHIYLLPECAIQQVC
NAVYVRLNYTQPLCACPARYRDPCSASLNADDLHTTRLTTDSKKKYAKKAVTLVKTCEAVSEMRECRAPRDWSL
LALQNIRTGKSHYLVICRCPESHILEGPMTHDQPTYASVPGIRVYGMMCVRPAPSYQSGYKGNRYTSAQSTTSAY
KVDYSQPHSYPDKPVYNRFQSHSYLDSNYYNRRSRAARMRRMAQGNPPFPWYKVTELAQSLHWTN*

Cluster E:
>XP_012240454.1_kappa-scoloptoxin(11)-Ssd1b-like_[Bombus_impatiens]
SYVIISSKLNNTTFKYWDGINGESDLHECVINAVCSVTHNRFWVSSLTERLCRCSNGKECPWQWTKELGNSSIS
LNNKSHMKFCAPITELSTCKYNQEGIEIHGKSDRNNSYLIPYNVILNCNCPGLHYWRLKKYTYLENDFIIQTFKCV
KRRMCNTYEFCGHIRSDLYSTYYRCTCPENHLCIFQDRNKENVQELLYSGSAYKGYCLPFNNA*

Cluster F:
>PAA79353.1_hypothetical_protein_BOX15_Mlig014663g1_[Macrostomum_lignano]
AFTDPSGMRCPCAYLERGGFWDSPTAVSLGVSDRVYRERRCSCQHEVNCDWSWITTDKSITLTWLGSEDIRD
SEIQLQFCAPIFRPGAPFCPTSGGGAFALRQTGWWIDLHEAPRCRCAAETGPASIRRFRYAEPLEMGERLRNQI
VEFACGRMPPCDAAQACYGESLNKQGRRTGGQVLCTCPPDTYCPVYTGQGARTVPADPTPETSSNSTAEVAEE
LSDTGQRIELLCVTAGP*

Cluster G:
>XP_025090922.1_uncharacterized_protein_LOC112562116_[Pomacea_canaliculata]
DGDKGPKPCSSPNHVCTTLMRYPDGYTADEGPNCFCGETTACSSEWEENSDNTLTWQHYERGHWRVQYKF
CKPVFAEMRQCEPDERAVTINTEVKTWKPYIDQARCRCPHSTGTSTLYQLSGWKREGAQWKYHYNCHKVECK
YYPLGTELNKRGSECAKVYIDQYKTSDKVQEVIFLCACPEGQMCPARLRKDNKDEVDLKEDDHGIFVMKYCES
NHQTENNK*

Cluster H:
>XP_005110430.1_uncharacterized_protein_LOC101852515_[Aplysia_californica]
FVPPGYGGNRRPAICPLPSKPCRLQITMRHLNGTAAGEMETAPCTCNGRVECPSDWNDRNHVISRNLISTTSI
MTLNMMFCRPIQPPNYCSLGQEALVVSGAMTIPNNVEDFNCRCYGDSPLYLHNRRYAENHLIYHTYVCDYHK
PTCPNYVTRCMVMREDSVDYTCKCAQGLHCRPQEDDWTYPMYGYCRP

Cluster I:
>XP_003107017.1_hypothetical_protein_GCK72_022181_[Caenorhabditis_remanei]
FTSTMTLKRHYRRPVCEPSQSCSYSYQQFTFELCDCPQSKSCPMDNGVQLKGITYQFCGARELPECMPDEIAA
EINFLQTSIYCVCPLDQIYVKQKESSNASVKYVCEEKEMCEVGQMCGVGNPIVGIKQTCQCAANSRCQVTAPN
VFNPISIQNATCQPI*

Cluster J:
>CAI9739673.1_Hypothetical_predicted_protein_[Octopus_vulgaris]
HQARVPYRVSNLMNVVLPQRYIPVCPTLDHVCGRVWRKGIYIQHICSCPGKTQCSMEWDDSVTGRSFQVKKY
VQDKYCGPAKVERVCAPKEISFSVVSAIKPQREVYRQGVVCLCKSPYGVHQNAIYELRDRNDLKYVHVFSCGKL
PKCTRYEPCAQYFTNIENGRRTENFIARVNLCTCPRPQHCIKPRTKVRQEELVGPGNIPYDLHYCSA*

Cluster K:
>CAH1780065.1_unnamed_protein_product_[Owenia_fusiformis]
TSEPIPPGVHHYGSDDEPCSSDMSICASVEHWEAFDGELPTTVITKKCGCQGNLKCPVSDRFDPNDGHTVIKG
GLYYKTCAPLTSTRRCRRSERALTMSGQKEVVEVRCQCRRGKVPTVYATWKPDEWTYDNPVFPFTNHATCGEL
RQCDIQTIRRNGKTQPCSKYYPRMNTDGPIPVPVTDFKGEVEMCTCPDSKFCDGRLESEMTAYTMGRDKNGFY
DFRYCMGL*

Cluster L:
>XP_013384870.1_uncharacterized_protein_LOC106154879_[Lingula_anatina]
ATTSRNPLTIVLPEGVDSQTDLPTCTPTQVCSEVRPYLPAWLSTARSSQMRDVTWLPRNATLFCKCPDEASPCAT
SFGFDSLSGQIYNGAKLHTCGAPSPVPYCAFSTPGVLKKEPVKNAAGKDTVKIELSCRCPGDSRAVFSDPKLEIT
TFENEIKTVDISCERVSEICSIENDAESPQCLDVSYEYGDESQIVLGDVSRPCRCPGGNTCLQYYRHWSRLMQE
ARTAIRRNRKTISPSMMQRFNYRVKRKELLPLCFPGSLENSTEYFNEGVNSDDQLPHCSHPNQLCSVVKPHSL
VLGRKTMTKTVTKYCRCSGDKPCRDDFTPDPVRSLIIGGKQYHTCKGNAGVPSEERPYQAMVCEVKTVYPNTEF
TWTEARAGLSSDQLKPYDVSKYVTLDCKPQGSRYGATFRTGELSRYGAGMAGVDGNLTLTTSRKTDYISVNHCR
YKKESDVCGTVTLNYQHLYVTTQKPAWEFAKAEQDCMCPSDSICPIEKWVKDVHTGNVTDLTPRYNGPVTVTIR
CQKKPE*
